# Divergent Inclusion Body Structures and Stabilities Emerge from Native Monomer Properties

**DOI:** 10.64898/2026.06.01.729379

**Authors:** Bruna Siebeneichler, Xiaoyue Liu, Pedro E. Rodriguez Cruz, Dalia Naser, Grace DelMistro, Julia Steckner, Anna Schaefer, Elizabeth M. Meiering

## Abstract

Protein aggregation is of broad importance in biotechnology and disease, yet the structural heterogeneity of cellular aggregates has confounded high-resolution structural analysis. Inclusion bodies (IBs) formed in *Escherichia coli* are an attractive, controllable system for unravelling the complexities of protein aggregation in a cellular context. Here, a multimodal analysis integrating residue-resolved quenched amide hydrogen–deuterium exchange (qHDX), proteolysis, FTIR, Congo red binding, and chemical denaturation is applied to IBs formed by proteins encompassing stable β- and α-globular folds, a partially structured protein fragment, and intrinsically disordered low complexity domains (LCDs). Remarkable conformational diversity is observed: IBs formed by well-folded proteins are extensively structured and include substantial local native-like features, whereas proteins with decreased access to stable native conformations form more heterogeneous and dynamic aggregates increasingly shaped by intrinsic sequence features. Strikingly, qHDX protection of TDP-43 LCD IBs strongly aligns with the core of cryo-EM structures of *ex vivo* pathological fibrils; however, peripheral regions that appear fully hydrogen-bonded in the cryo-EM structures exhibit little protection. The results reveal that individual protein IBs contain distinct mixtures of native-like, disordered, and amyloid-like conformers, informing the prediction and control of cellular aggregate structure and stability.

**Entry for the Table of Contents:** Inclusion body conformational heterogeneity. Left: aggregates retaining local native-like structure. Middle: assemblies populated by α-helical intermediates coexisting with antiparallel β-sheet oligomers. Right: initially disordered states undergo helix–helix interactions and progress towards β-sheet–rich conformations, illustrating distinct routes of cellular aggregation.

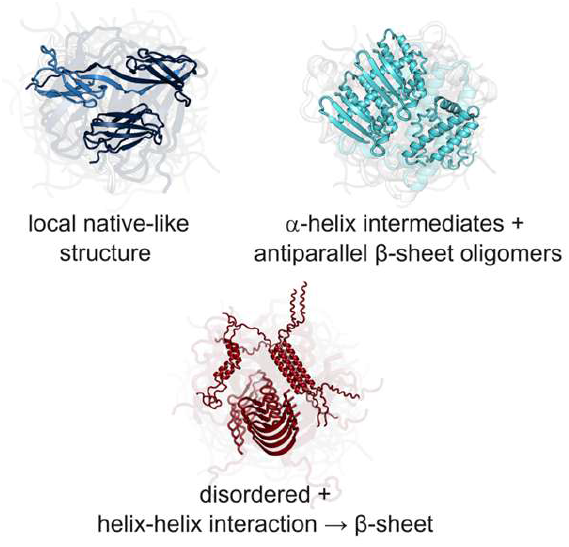

## Introduction

Protein aggregation is an area of intense current research owing to its importance in many fields, ranging from protein production and materials in biotechnology to the formation and therapeutic targeting of toxic protein assemblies in disease.^[1–10]^ Confounding conformational heterogeneity is a major challenge for defining the structure and stability of such assemblies.^[7,8,10– 14]^ Examples of these assemblies include functional protein condensates that form during liquid–liquid phase separation (LLPS), inclusions associated with stress and disease, and inclusion bodies (IBs) formed during recombinant protein production in research and technology.^[5,7,11,13–15]^ Currently, the specific molecular organization of these assemblies *in vivo* remains obscure, motivating new approaches for their characterization.

The formation of IBs in *Escherichia coli* provides a valuable system for elucidating cellular protein aggregation. Beyond their broad significance for protein production,^[1,16,17]^ IBs offer a controlled system for analyzing how intrinsic protein properties determine aggregate structure and stability.^[13,18,19]^ Importantly, much of the current structural understanding of protein aggregation derives from *in vitro* studies of purified proteins; however, recent studies are highlighting that aggregates formed *in vitro* can be very different from those that form *in vivo*.^[7,20]^ Recently, we reported detailed analysis of IBs based on quenched amide hydrogen–deuterium exchange (qHDX) applied to superoxide dismutase 1 (SOD1) mutants associated with amyotrophic lateral sclerosis (ALS).^[12,21]^ In contrast to previous studies that reported prominent amyloid structure in IBs,^[22]^ SOD1 IBs also contain substantial disordered and notably native-like conformations, revealing that IB structures do not conform to a common amyloid architecture.

We demonstrate here how IB properties are broadly governed by the native structural and physicochemical features of the constituent protein monomers (Figure 1, Table S1). The studied proteins encompass diverse native states: Adnectin, a highly stable β-sandwich scaffold protein from a newly approved therapeutic protein class,^[23]^ all-α Apomyoglobin and its partially folded N-terminal 77 residue fragment (ApoMb153 and 77),^[24]^ and intrinsically disordered low-complexity domains (LCDs) from heterogeneous nuclear ribonucleoprotein A2 (hnRNPA2^LCD^) and TAR DNA-binding protein 43 (TDP-43^LCD^), which undergo LLPS naturally and can transition under stress or disease conditions to more solid-like or fibrillar states.^[7,25,26]^ High-resolution qHDX measurements combined with FTIR, Congo red binding, proteolytic digestion, and chemical denaturation reveal that IBs do not adopt common predominantly amorphous or amyloid structures, but instead exhibit major differences in structure and stability linked to the properties of the constituent protein monomers.

**Figure 1.**
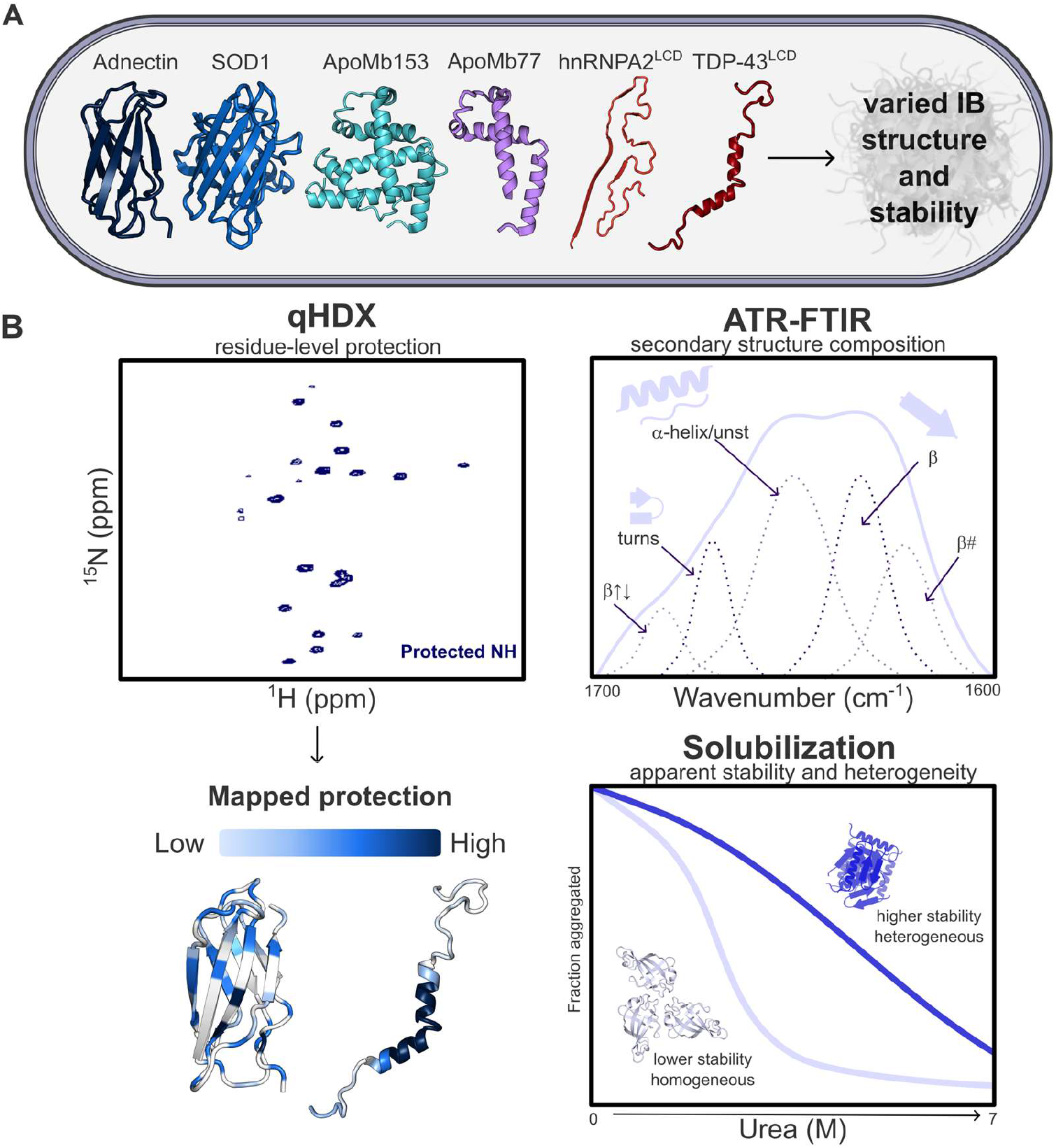
Overview of proteins and main methods for investigation of inclusion body (IB) structure and stability. A) The panel includes well-folded globular proteins with distinct secondary structure: β-sheet-rich proteins (Adnectin; reduced apo SOD1) and an α-helical protein (Apomyoglobin, full-length, ApoMb153). Additionally, proteins with reduced stability or intrinsic disorder were examined, including N-terminal 77 residue fragment of Apomyoglobin (ApoMb77) and low complexity domains of hnRNPA2 (hnRNPA2^LCD^, fused to a TEV-cleavage tag) and TDP-43 (TDP-43^LCD^). Structures are represented using (left to right): PDB codes 1FNA, monomer of 2GBU, 1VXF, AlphaFold3 model,^[50]^ 2N3X. B) Complementary biophysical methods for IB characterization: qHDX NMR reveals protection in IB of individual amides against exchange with D_2_O; FTIR reveals secondary structure composition; urea solubilization reports on stability and heterogeneity of IBs.

## Results and Discussion

### Quenched Hydrogen/Deuterium Exchange (qHDX) NMR Uncovers Diverse Protein Conformations in IBs

qHDX NMR is a powerful yet little applied method for obtaining a high-resolution view of IB structure by measuring the protection of individual backbone amide hydrogens against exchange with solvent deuterons. In general, such protection is conferred by hydrogen bonding and/or burial of amide groups due to native and/or non-native protein structure in aggregates.^[12,13,21]^ The qHDX experiment involves incubating IBs in D_2_O, followed by quenching of exchange, sample solubilization in DMSO, and amide protection measurement using solution NMR. For these experiments, we first made sequence-specific resonance assignments in DMSO, establishing many amide probes throughout the sequence of each protein (Figure S1 and Tables S2-S6). Measurements of qHDX protection of these amides were then compared with protection in the native soluble protein and with protein properties that may influence aggregation, including hydrophobicity and primary sequence-based aggregation-prone regions (APRs) predicted by Tango, AGGRESCAN, PASTA, and Zipper DB.^[27–31]^

Notably, across the proteins examined below, we observe extensive evidence for protection conferred by local native-like as well as amyloid-like structures. Here, we define *local* native-like structure in IBs as secondary structure and/or packing in restricted regions of the protein, which resemble the native state and may share features with folding intermediates.

#### Adnectin

Adnectin IBs exhibit an extensive and uniform qHDX protection pattern throughout the protein (Figure 2A, F). Many protected residues coincide with native β-strands and partially overlap with residues protected in the soluble native monomer and with hydrogen bonding in the crystal structure of the highly homologous WT ^10^FN_3_ (Figures 2B, C, S2).^[32]^ Also, residues in the BC and FG loops exhibit higher protection than the native monomer. Some loops, including BC and FG, are relatively hydrophobic and coincide with APRs (Figure 2D, E). In addition, the FG loop forms new hydrogen bonds and packs against the BC loop in predicted domain-swapped structures for this Adnectin (Figure S3) and homologs,^[33,34]^ which may contribute to increased protection.^[35]^ Further support for retained native structural features is provided by IB proteolysis monitored by mass spectrometry (MS) (Figures 2F, S4). Although the major IB species observed by MS is the full-length protein, proteolysis is observed in the BC loop (L26) and strand E (Y55), which are relatively exposed sites in both the monomer and domain-swapped model structures (Figure S3). During expression, Adnectin is initially observed in both soluble and insoluble fractions and later only in the insoluble (Figure S10A).^[36]^ These observations, together with the fast folding,^[37]^ and high stability of Adnectin (*T*_*m*_ of 78°C),^[36]^ suggest folded Adnectin monomers can be recruited into IBs, as also proposed for other proteins.^[18]^

**Figure 2.**
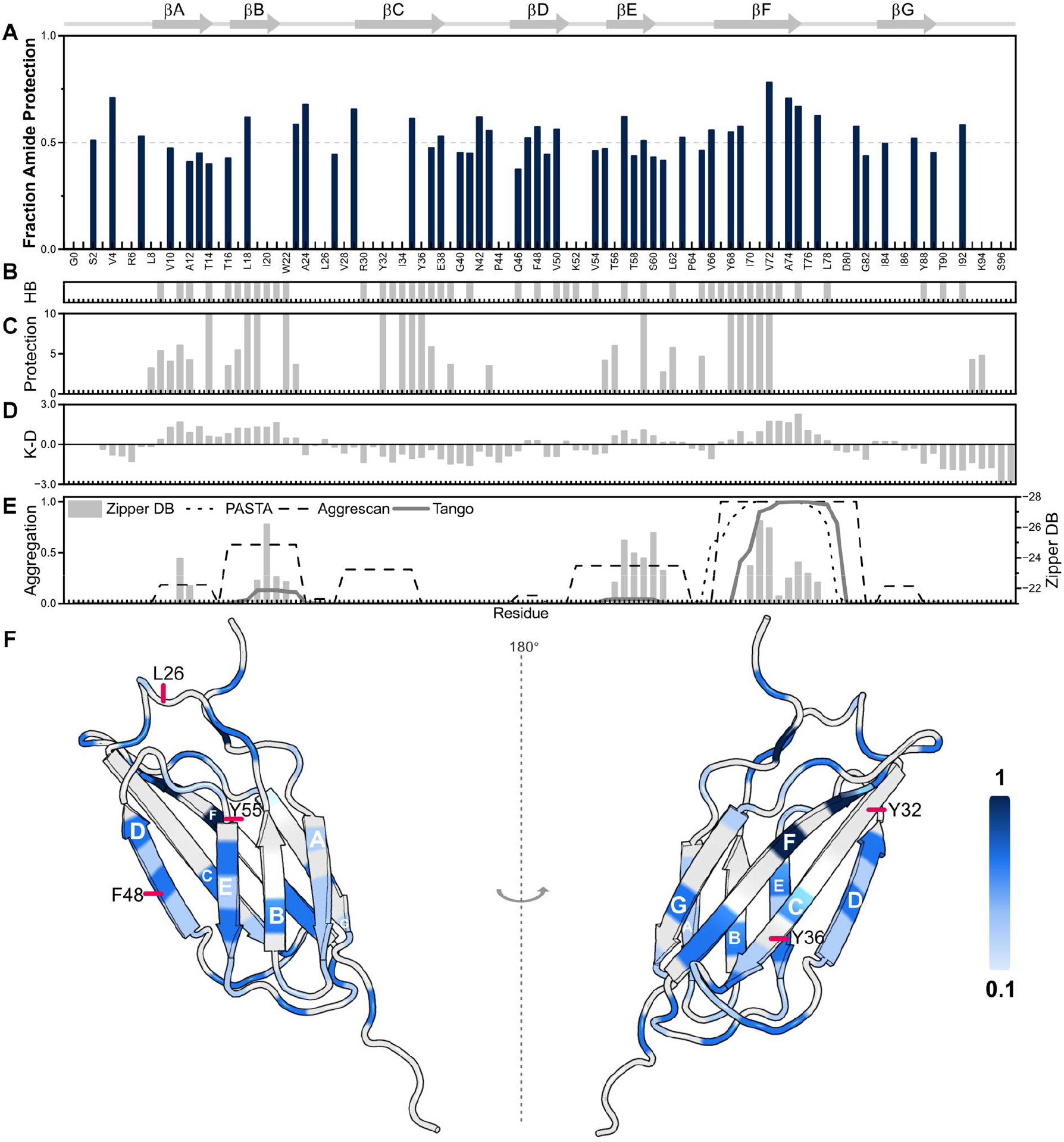
qHDX analysis of Adnectin IB. A) Fraction of amide protected against exchange with solvent. Bars are average values of three biological replicate measurements. Gaps indicate areas of no data. B) Hydrogen-bonded backbone amides in the crystal structure of wild-type 10^th^ fibronectin type III domain (1FNA), from which Adnectin is derived. C) HDX protection of monomeric native Adnectin. Measured amide exchange rate constants, k_ex_ (s^−1^), at pD 4.6 are plotted as −log k_ex_. Rate constants smaller than 1×10^−8^ s^−1^ are set to 10.^[32]^ D) Kyte-Doolittle hydrophobicity, positive values are hydrophobic, negative are hydrophilic.^[27]^ E) Sequence-based Aggregation propensity predicted by: TANGO, AGGRESCAN, and PASTA, normalized from 0 (no aggregation propensity) to 1 (highest aggregation propensity); and ZipperDB.^[28–31]^ F) qHDX fraction amide protection from A) shown on the AlphaFold3 predicted structure;^[50]^ values from 0.1 to 1 coloured as light to dark blue, respectively. Light grey indicates no data. Red lines indicate proteolysis sites determined by MS (Figure S4).

#### SOD1

Similar to Adnectin, monomeric reduced apo SOD1 exhibits an IB qHDX protection pattern indicative of significant local native-like structure, as reported previously and considered further here for the ALS-associated A4V mutant (SOD1_A4V_, Figure S5).^[12]^ The observed protection is consistent with contributions from native-like monomers, dimers, and non-native conformations, including some amyloid-like structures. Many of the protected amides occur in strands β1-β8 of the β-barrel, while protection in some loops and N- and C-terminal strands is associated with native dimer structure.^[12]^ The protection in the long loops (4 and 7) is greater in IBs than in native dimers, suggesting that non-native dimers previously observed for SOD1 may also contribute to protection.^[38,39]^ Such associations may still constitute relatively accessible regions, as also suggested by proteolysis observed in loop 4 (Figures S4, S5). Comparisons with cryo-EM structures of SOD1 amyloid produced *in vitro* show partial agreement with qHDX protection observed in IBs; differences may arise due to the increasingly recognized high sensitivity of aggregation to experimental conditions.^[6,7]^ It is notable that SOD1 and Adnectin IBs have similar qHDX behaviour in IB cellular aggregates, with extensive and uniform protection indicating local native-like structure as well as non-native interactions.

#### ApoMb153 and 77

We next examine the predominantly helical, globular, 153-residue protein ApoMb and its N-terminal 77 residue fragment (Figure 3). For ApoMb153 IBs, residues in the relatively hydrophobic helices αA, αE and αG exhibit the highest protection, while the more hydrophilic helices, αC, αD, and αH, show moderate protection, similar to native myoglobin amide exchange (Figures 3A-D, S2).^[40]^ These results again point to the presence of local native-like structure in IBs. As for Adnectin and SOD1, proteolysis is observed in solvent accessible turns and loops (Figures 3F, S4), consistent with retention of locally ordered structure within IBs. Proteolysis is also observed in αF, which is poorly folded in native ApoMb153, and in parts of αH that have relatively low stability in the native protein.^[41,42]^ Despite lacking bound heme, ApoMb153 has substantial thermodynamic stability, and during folding, it forms multiple intermediates containing structured αA, αB, αG, and αH, which can be well packed.^[43–45]^ Together, these observations suggest the presence of multiple species with native-like features in ApoMb153 IBs.

**Figure 3.**
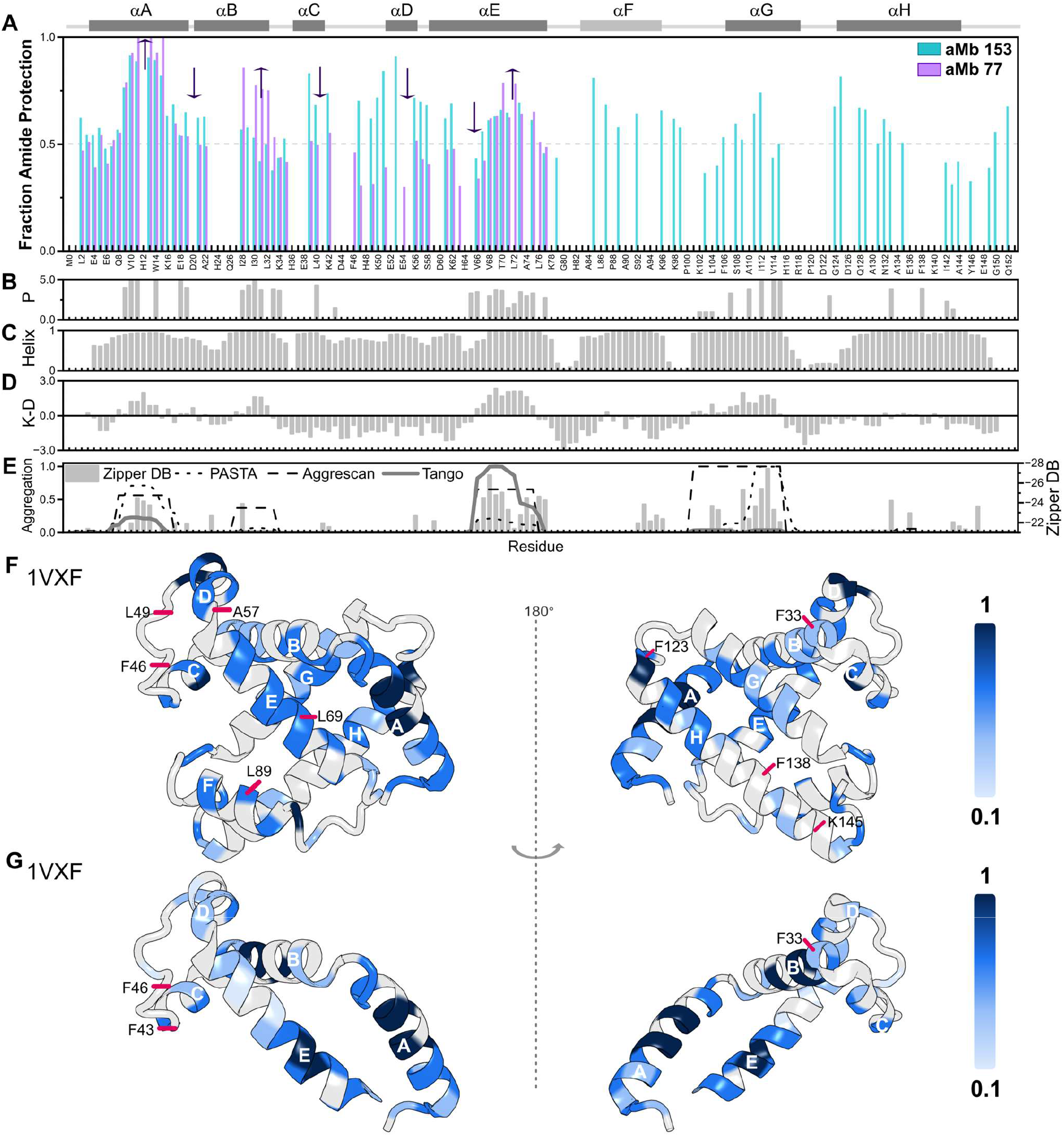
qHDX analysis of ApoMb153 and ApoMb77 IBs. A) Fraction of amide protected against exchange with solvent for ApoMb153 and ApoMb77, with arrows indicating an increase or decrease in protection for ApoMb77 relative to ApoMb153. Bars are average values of three biological replicate measurements. Gaps indicate areas of no data. B) HDX protection factor, P, of monomeric native ApoMb153.^[40]^ C) Helical propensity of ApoMb153 from PASTA.^[30]^ D) Kyte-Doolittle hydrophobicity, positive values are hydrophobic, negative are hydrophilic.^[27]^ E) Sequence-based Aggregation propensity predicted by: TANGO, AGGRESCAN, and PASTA, normalized from 0 (no aggregation propensity) to 1 (highest aggregation propensity); and ZipperDB.^[28–31]^ F) qHDX fraction amide protection from A) shown on the crystal structure of ApoMb153 (1VXF), values from 0.1 to 1 coloured as light to dark blue, respectively. Light grey indicates no data. Red lines indicate proteolysis sites determined by MS (Figure S4). G) Results for ApoMb77 shown as in F) using residues 1-77 of 1VXF because ApoMb77 is known not to be fully helical.^[24]^

In contrast, the partially folded and aggregation-prone ApoMb77 forms IBs with a markedly different protection pattern. ApoMb77 forms fluctuating helices and β-rich oligomers in solution and its IBs show increased protection in αA, αB, and the C-terminal of αE together with decreased protection in αC, αD, and the N-terminal of αE (Figure 3A).^[24]^ These changes are consistent with the loss of stabilizing interactions contributed by the C-terminal half of ApoMb153, leading to increased exposure and subsequent burial of hydrophobic and APRs of the fragment (Figure 3D, E). Consistent with this interpretation, qHDX protection for ApoMb77 correlates with hydrophobicity, unlike ApoMb153, Adnectin, or SOD1 (Figure S2B). ApoMb77 also shows proteolysis in the C-D loop similar to ApoMb153, but lacks cleavage in helix E, supporting a role for this hydrophobic region in aggregation. Thus, whereas ApoMb153 IBs retain substantial native-like features, the ApoMb77 protection pattern appears to be driven more strongly by burial of otherwise exposed hydrophobic residues.

#### hnRNPA2^LCD^ and TDP-43^LCD^

Further insight into how disordered proteins may contribute to IBs was obtained by examining intrinsically disordered regions from hnRNPA2 and TDP-43. The hnRNPA2_LCD_ construct contains residues N266-Y341, preceded by an N-terminal hexa-histidine tag and TEV protease site (SI 1.2).^[46]^ hnRNPA2_LCD_ exhibits particularly low amide protection (< 0.5, Figure 4) and readily undergoes extensive proteolysis (SI 1.1, Figure S4, S10). Increased protection observed near the tag is consistent with MS evidence for protease-resistance in this region. Aggregation predictors identify high aggregation propensity near the tag (Figures 4C, S2C), while AlphaFold3 predicts an ordered β-strand within the tag (Figure 4D), which may promote aggregation.^[11,47]^ TEV sites have also been reported to decrease protein solubility.^[48]^ Although hnRNPA2_LCD_ has been observed to form fibres, these contain low-complexity aromatic rich kinked structure (LARKS) with short β-strands,^[25]^ characteristic of functional amyloids.^[11]^ Mapping IB qHDX onto the cryo-EM structure of *in vitro* hnRNPA2 fibrils lacking the tag reveals generally dispersed low protection, consistent with the lower stability of functional relative to pathogenic amyloids.^[11,25]^ Together, these observations suggest that the hydrophobic, β-prone tag contributes disproportionately to ordered structure formation in hnRNPA2^LCD^ IBs, while much of the LCD region remains relatively disordered.

**Figure 4.**
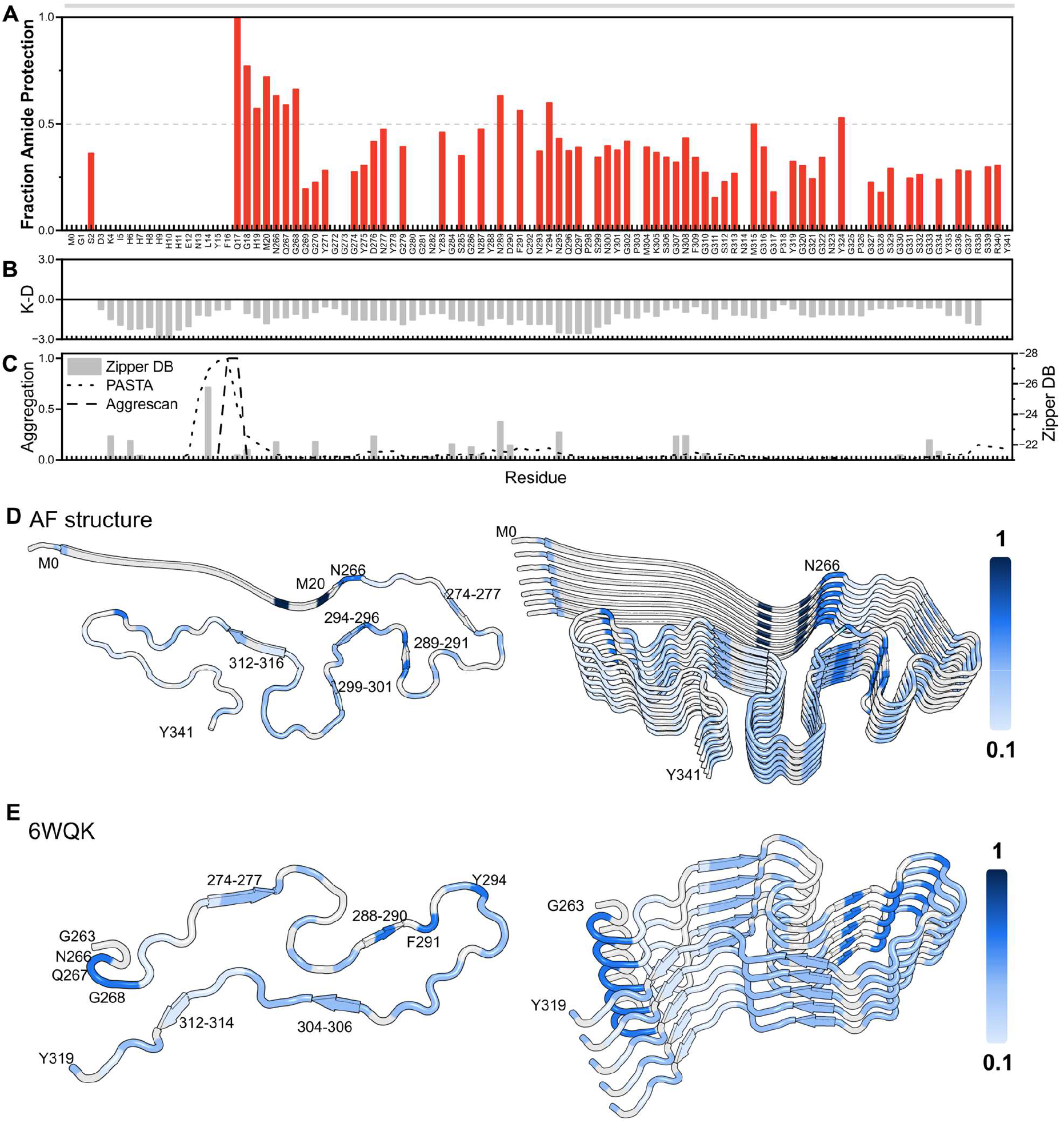
qHDX analysis of hnRNPA2^LCD^ IB. A) Fraction of amide protected for hnRNPA2^LCD^. Bars are average values of three biological replicates. Gaps indicate areas of no data. B) Kyte-Doolittle hydrophobicity, positive values are hydrophobic, negative are hydrophilic.^[27]^ C) Sequence-based Aggregation propensity predicted by: TANGO is 0; AGGRESCAN and PASTA, normalized from 0 (no aggregation propensity) to 1 (highest aggregation propensity); and ZipperDB.^[28–31]^ D) qHDX fraction amide protection from A) shown on AlphaFold3 predicted multimer structure,^[50]^ values from 0.1 to 1 coloured as light to dark blue, respectively. Light grey indicates no data. Predicted structure confidence is very low (average pLDDT 34). E) Protection from A) shown as in D) on fibril structure of hnRNPA2^LCD^ (6WQK).

The TDP-43^LCD^ construct consists of residues 281–360 with a C-terminal hexa-histidine tag (SI 1.2). In contrast to hnRNPA2_LCD_, TDP-43_LCD_ has a strongly protected region spanning A321–M339 (Figure 5), and no proteolysis is observed for A324–M337 (Figure S4). This conserved region forms transient α-helices in the soluble monomer and contains the most hydrophobic stretch in the sequence, which drives helix-helix assembly.^[26]^ The qHDX pattern agrees with previous *in vitro* studies in which the native helices initiate aggregation through hydrophobic self-association before transitioning to β-sheet.^[26,49]^ In contrast, the remainder of the TDP-43^LCD^ sequence shows relatively little protection, indicating greater disorder or solvent accessibility.

**Figure 5.**
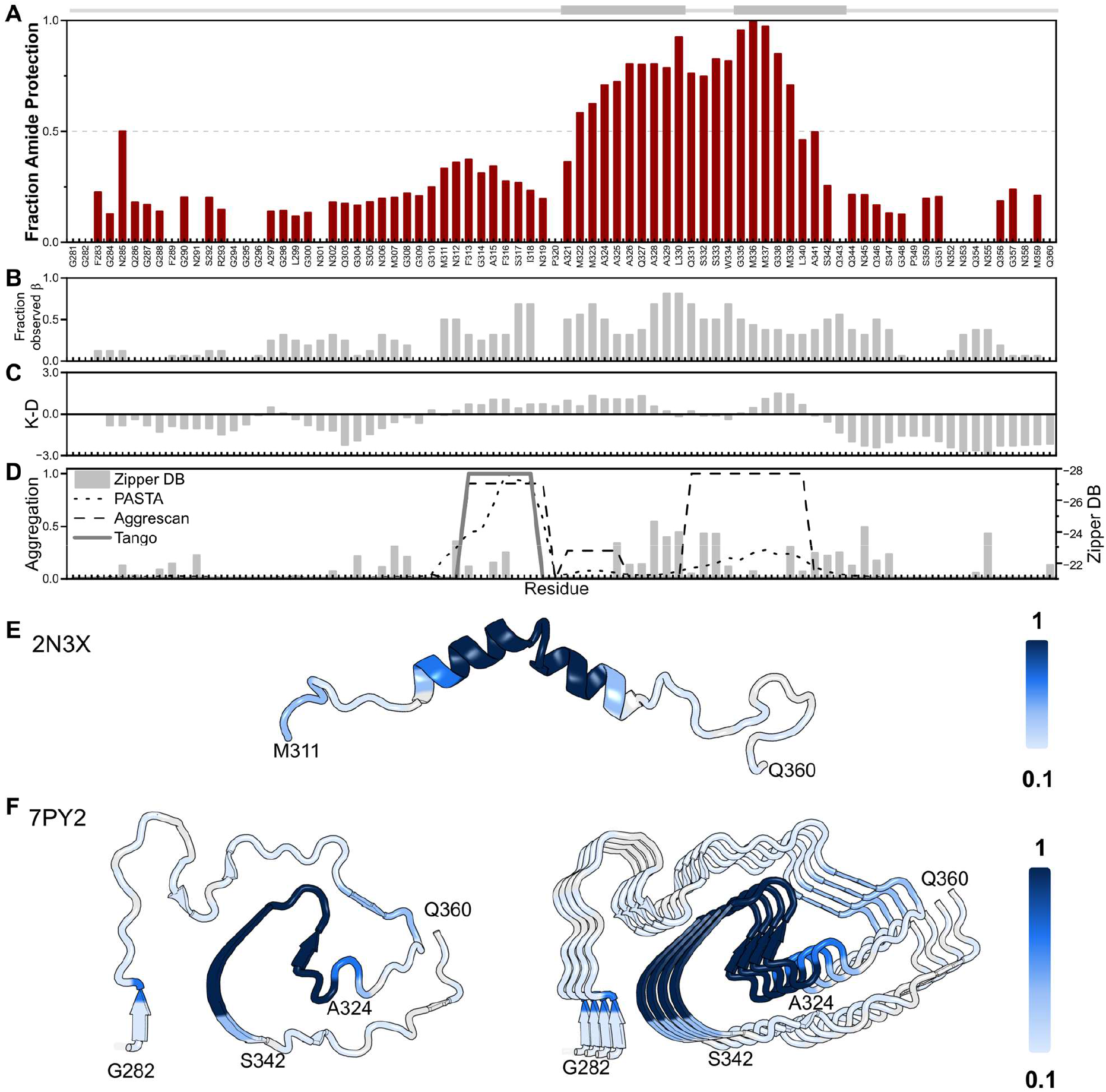
qHDX analysis of TDP-43^LCD^ IB. A) Fraction amide protection for TDP-43^LCD^. Bars are average values of three biological replicates. Gaps indicate areas of no data. B) Fraction observed β-conformation in amyloid-like structures available for TDP-43^LCD^ (more information given in Figure S6). C) Kyte-Doolittle hydrophobicity, positive values are hydrophobic, negative are hydrophilic.^[27]^ D) Sequence-based Aggregation propensity predicted by: TANGO, AGGRESCAN, and PASTA, normalized from 0 (no aggregation propensity) to 1 (highest aggregation propensity); and ZipperDB.^[28–31]^ E) qHDX protection shown on TDP-43^LCD^ (residues 311–360) NMR solution structure, with values from 0.1 to 1 coloured as light to dark blue, respectively. Light grey indicates no data. F) Protection from A) shown as in E) on TDP-43^LCD^ (residues 282-360) fibril structure from a patient having ALS with FTD (7PY2).

Mapping this qHDX protection onto cryo-EM structures of amyloid-like fibrils (Figures 5F, S6) reveals a striking correspondence with an *ex vivo* pathological TDP-43_LCD_ filament structure (7PY2) with the highest protection concentrated in the inner long β-strands of the filament core.^[49]^ Even more remarkable is the comparatively low protection of the adjacent structure, despite being fully hydrogen-bonded in the cryo-EM structure.^[49]^ As noted by Arseni *et al*., due to the high glycine and neutral polar residue content, the 7PY2 structure has many β-turns and lacks the β-sheet stacking typical of cross-β amyloid structure. Furthermore, pathological *ex vivo* structures tend to contain longer β-strands than *in vitro* aggregates. Thus, unlike the more limited similarities observed for *in vitro* aggregates,^[6,51]^ the correspondence of TDP-43_LCD_ IB protection with *ex vivo* patient-derived filaments reveals unexpected structural similarities among cellular aggregates.

Overall, qHDX reveals that IB structure is strongly shaped by the conformational tendencies of the precursor protein (Table S1). Adnectin and ApoMb153 IBs most closely retain protection patterns resembling their native monomers, whereas ApoMb77 and TDP-43_LCD_, which cannot form stable native structures, exhibit protection patterns that correlate more strongly with residue hydrophobicity (Figure S2). Across the entire protein set, residues located in structured regions of the native monomers show a modest tendency toward increased protection in IBs relative to residues from unstructured regions (Figure 6A). The average fraction of protected amides decreases in the order ApoMb77, Adnectin, ApoMb153, hnRNPA2^LCD^, SOD1_A4V_ and TDP-43^LCD^ (Figure 6B), potentially reflecting differences in monomer stability, as native SOD1_A4V_ has a substantially lower T_*m*_ than Adnectin and ApoMb153 (Table S1). To gain further insight into the basis of these protection patterns, we next compare qHDX with complementary optical measurements of secondary structure composition and relative IB stability.

**Figure 6.**
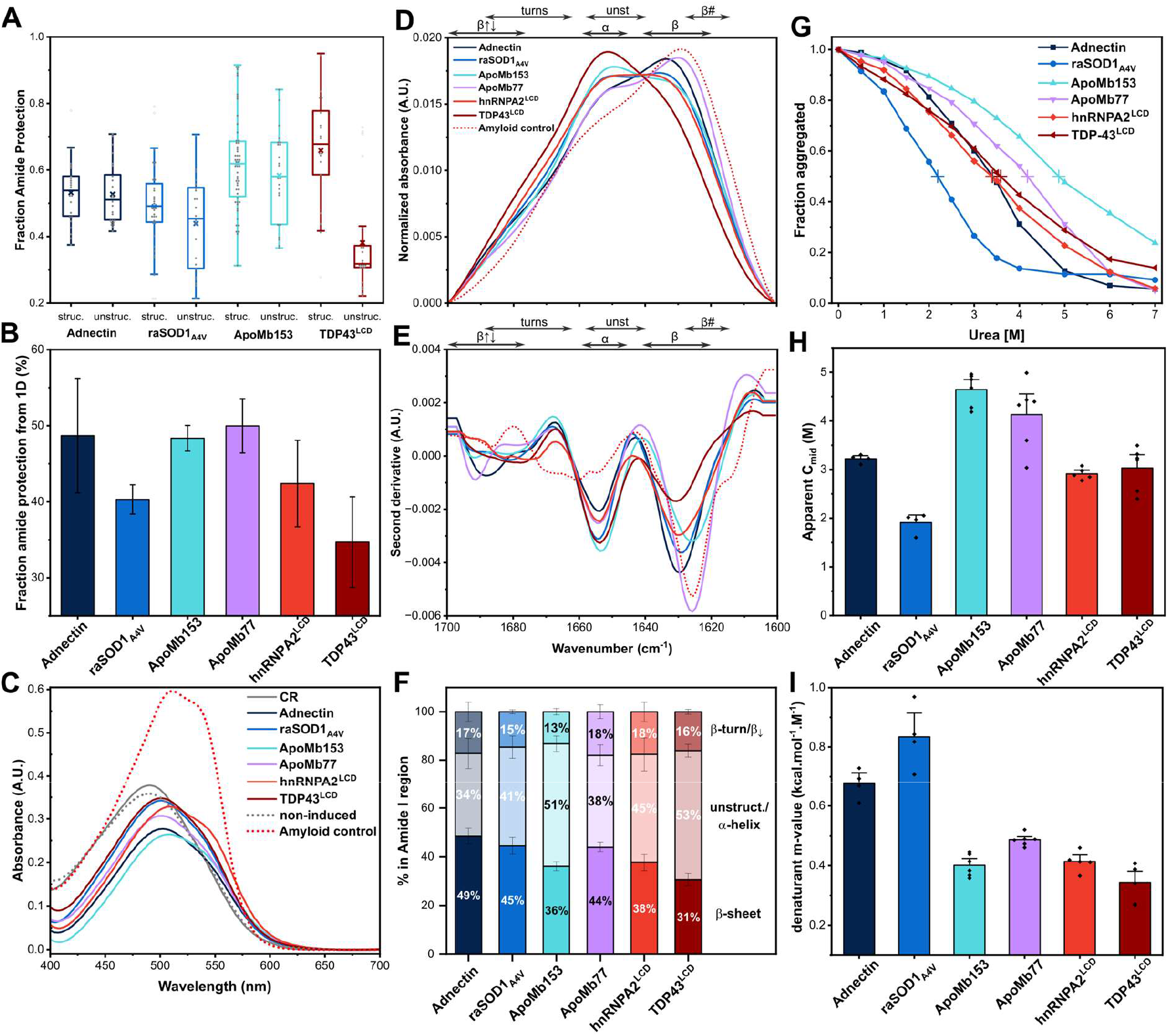
IB stability and structure by amide protection against exchange, CR, FTIR and chemical solubilization. A) Fraction protection for individual amides grouped as structured (H-bond) or unstructured (unstruc) based on native protein monomer structure (Adnectin: 1FNA, raSOD1: 2GBU, ApoMb153: 1VXF and TDP-43^LCD^: 2N3X). B) Fraction protection of all amides from qHDX results calculated from amide region of 1D ^1^H spectra. C) CR shifted absorbance spectra of IBs (average of two biological replicates and one technical). Sonicated pure SOD1 is the amyloid control.^[12]^ D) FTIR spectra (average of at least two biological replicate samples, each with three technical replicate samples) and E) corresponding FTIR second derivative spectra for the amide I region. Spectral regions for antiparallel β-sheet (β⇵), turns (including loops), α-helix (α), unstructured (unst), β-sheet (β), and β-amyloid with strong intermolecular hydrogen bonds (β#) are indicated. F) Integrated area of secondary structure bands from curve-fitting of spectra using 3 components: β-turn/antiparallel β-sheet (~1680 cm^−1^), unstructured/ α-helix (~1654 cm^−1^) and β-sheet (~1630 cm^−1^). Further details for fitting are given in the SI (Figures S7, S8, Tables S13, S14). G) Urea chemical solubilization of IBs (average of at least two biological replicates, each with two technical replicates), and corresponding fitted values: H) apparent *C*_*mid*_ and I) apparent denaturant dependence of solubilization (*m*-value).

### Distinct IB Secondary Structure Compositions and Amyloid Content Revealed by FTIR and Congo Red

FTIR absorbance spectroscopy is an established method for qualitative analysis of secondary structure of aggregates, such as amyloid, oligomers, and IBs (see 3.1 SI R&D 1).^[12,13,52–54]^ Absorbance features in the amide I region provide insight into β-sheet organization, disordered and α-helical content, and the contributions of native-like and amyloid-like conformations. Aligned with the qHDX results, FTIR shows clear differences between the investigated proteins (Figure 6D-F, Table S1).

The Adnectin IBs exhibit a high proportion of β-sheet and β-turn/antiparallel β-sheet structure (49% and 17%, respectively), together with unstructured/α-helical content (34%). Importantly, the FTIR spectrum shows a distinct low-intensity band at ~1690 cm^−1^ (Table S13), which is characteristic of antiparallel β-structure.^[54–56]^ A similar feature is observed in the native protein (Figure S7). Parallel β-sheet, common in mature amyloid, has negligible absorbance at these higher wavenumbers, while also exhibiting a strong lower-wavenumber β-band similar to antiparallel β-sheets (SI R&D 3.1).^[57]^ Accordingly, the strong β band alone cannot definitively distinguish parallel from antiparallel β-structure. While sharing features with native Adnectin, the IBs also show increased signal for unstructured/α-helix (34% vs 25%) (Figure S7, Table S14). Thus, the FTIR results indicate that Adnectin IBs retain significant native-like secondary structure while also exhibiting increased disorder relative to the native monomer. Similar results have been reported previously for SOD1 IBs, which also have prominent β-structure, including antiparallel features, and even stronger correspondence to the soluble monomer (Figure S7, Tables S13, S14).^[12]^ Together, these results indicate that neither Adnectin nor SOD1 IBs are dominated by the parallel β-sheet structure characteristic of mature amyloid,^[13,52,55,58,59]^ and they retain structurally heterogeneous ensembles including native-like features.

For ApoMb153, the IB FTIR spectrum shows a prominent signal in the α-helix/unstructured region together with features characteristic of antiparallel β-sheet (Figure 6D, E, Table S14).^[54,55]^ The spectrum has similarities to that of native ApoMb153, but with reduced helical signal and increased intensity in the lower and higher wavenumber β-sheet bands (Figure S7). Although α-helix and unstructured absorbance overlap, the ApoMb153 IB spectrum clearly has a stronger signal (51%) in this region than the less helical ApoMb77 (38%, Figures 6F, S7), suggesting higher helical content in ApoMb153 IBs. Concomitantly, ApoMb77 IBs exhibit a higher β-sheet signal (44%) than ApoMb153 (36%); this may be related to the increased protection identified by qHDX in hydrophobic regions (Figure 3). Features of antiparallel β-sheet (1695, 1624 cm^−1^) are also increased in ApoMb77, both for IBs and purified protein (including oligomers),^[24]^ which show remarkably similar FTIR spectra (Figure S7 and Table S13). Collectively, these results indicate that ApoMb153 IBs retain some native-like helical structure, whereas ApoMb77 IBs are more β-enriched and resemble oligomeric assemblies formed by the pure protein.^[24]^

In contrast to the more structured IBs described above, the low complexity domain IBs exhibit a strong unstructured/α-helical FTIR signal together with comparatively weaker β-sheet signal (Figure 6D-F). hnRNPA2_LCD_ (TDP-43_LCD_) shows a mixture of 45% (53%) unstructured/α-helix and 38% (31%) β-sheet. TDP-43_LCD_ has the highest proportion of unstructured/α-helix signal of all the IBs, as well as the lowest qHDX protection, which includes only a relatively small stretch of protected amides (Figures 5, 6B). Previous studies showed that TDP-43^LCD^ aggregates do not exhibit typical amyloid properties, such as strong CR binding or high stability.^[60]^ Also, some fibrillar TDP-43 constructs contain substantial unstructured/α-helix content by FTIR despite possessing an ordered core observed by ssNMR and cryo-EM.^[61,62]^ These observations suggest that TDP-43^LCD^ IBs contain aggregates with intermediate properties on the assembly landscape, including a small core component with ordered structure (amyloid-like/LARKS) flanked by disordered regions.

Complementing FTIR, Congo red (CR) absorbance spectroscopy is a standard method for qualitative assessment of the presence of amyloid in aggregates.^[13,63,64]^ The IB samples show small CR spectral shifts compared to the amyloid control, with the largest response for hnRNPA2^LCD^ (Figure 6C). These small shifts may indicate the presence of some amyloid or β-structure, as well as structural heterogeneity in IBs. While IBs are generally porous structures,^[65]^ CR may underestimate amyloid or β-structure due to the limited accessibility of the dye to tightly packed aggregates.^[66]^ This limitation does not apply to FTIR, which offers a distinct view of IB secondary structure features. Together the FTIR and CR measurements complement the qHDX results, showing that IBs are structurally diverse assemblies whose properties depend strongly on the precursor protein.

### Urea Solubilization Reveals Differences in IB Stability and Heterogeneity

Urea solubilization monitored by turbidity (Figure 6G) was used to assess IB stability and structural heterogeneity.^[19,67,68]^ The midpoint of solubilization (*C*_*mid*_) is a measure of the apparent stability of IBs against solubilization by urea while the denaturant-dependence of solubilization (*m*) is an indicator of aggregate heterogeneity (SI 1.9, 3.2, Figures 6G-I, S9, Table S1).

The urea solubilization profiles reveal significant variations in *C*_*mid*_ and *m* among the different protein IBs (Figure 6H,I). SOD1_A4V_ has the lowest apparent stability, as it is fully solubilized after 3 hours at a relatively low urea concentration, while ApoMb153 is the most stable, remaining partially insoluble after 24 hours in 7 M urea (Figure S9). The native β-proteins, SOD1 and Adnectin, exhibit higher denaturant-dependence (0.83 and 0.67 kcal mol^−1^ M^−1^, respectively), suggesting more uniform aggregate species that solubilize at similar urea concentrations (Figure 6I). The relatively low *C*_*mid*_ values for Adnectin and SOD1_A4V_ (3.2 and 1.9 M, respectively) suggest that these IBs contain relatively weakly associated native-like species (SI R&D 2).^[12]^ Similar behaviour has been reported for non-classical IBs, such as for asparaginase IBs, which retain activity and have low C_*mid*_, further evidence for the presence of native-like conformers in IBs.^[1,2,19,69]^ By comparison, *in vitro* and pure amyloid structures are typically much more resistant to urea solubilization.^[11,68,70]^ Taken together, these results suggest that SOD1_A4V_ and Adnectin IBs contain native-like structure within relatively uniform assemblies.

ApoMb153 has the highest *C*_*mid*_ (~4.6 M, Figure 6H) and lowest *m* (0.4 kcal mol^−1^ M^−1^), indicating an overall more stable and structurally heterogeneous IB structure. The ApoMb153 monomer has substantial global stability and forms various partially folded intermediates.^[43,45]^ Strong interactions among various hydrophobic helices in ApoMb153, analogous to the behaviour of human growth hormone IBs,^[71]^ may likewise stabilize ApoMb153 IBs. ApoMb77 IBs have somewhat lower apparent stability (*C*_*mid*_ 4.1 M) and heterogeneity (*m* 0.49 kcal mol^−1^ M^−1^). Relative to ApoMb153, the absence of the C-terminal half of the protein weakens native structure, while increased exposure of hydrophobic residues may promote formation of β-oligomers.^[24]^ These interactions can be expected to confer higher urea resistance to ApoMb77 IBs compared to the more disordered LCD IBs, considered next.

The hnRNPA2^LCD^ and TDP-43^LCD^ IBs have comparatively low stability (*C*_*mid*_ 2.9 and 3 M, respectively) and high heterogeneity (*m* 0.41 and 0.34 kcal mol^−1^ M^−1^, respectively, Figure 6H,I). The native disorder and relatively high polarity of these LCDs may be expected to result in assemblies that are less stable and less uniform. Previous measurements likewise support relatively labile intermolecular organization in these systems: hnRNPA2_LCD_ forms hydrogels at 45 °C, that lose structure at 55 °C, while TDP-43_LCD_ shows weak and dynamic self-association, with only transiently populated dimeric species on the pathway to tetrameric and larger assemblies.^[25,26]^ For hnRNPA2_LCD_, while much of the IB ensemble appears to be structurally disordered, there is still evidence for a small fraction that is highly resistant to urea solubilization (Figure S9). This interpretation is consistent with the low overall qHDX protection for both proteins (Figure 6B), their extensive proteolytic susceptibility, and the strong unstructured/α-helical FTIR signal (Figure 6F), collectively suggesting structurally variable assemblies that are loosely packed and include smaller locally ordered cores.

### Precursor Native-State Properties Drive Unexpectedly Diverse Inclusion Body Structures

The converging multimodal evidence presented here reveals a remarkably broad spectrum of IB structures and stabilities, which are linked to properties of the precursor native state. This coherent view emerges from high-resolution quantitative qHDX NMR of IBs augmented by FTIR, CR, proteolysis, urea solubilization, and sequence and structural analyses. An overview of the determined IB structural features is given in Figure 7, with the corresponding experimental details summarized in Table S1. For well-folded globular proteins (Adnectin, SOD1, ApoMb153), there is strong evidence for substantial locally native-like conformations in IBs (Figure 7B). Adnectin IBs show qHDX protection that overlaps most of the native β-strands; additional protection in the BC/FG loop regions may arise from intermolecular association, such as domain-swapping (Figure S3).^[33,35]^ This interpretation is reinforced by FTIR, as spectra for Adnectin IBs retain features of the native protein. SOD1 IBs also exhibit extensive protection throughout the β-barrel and loop regions, reflecting native-like monomers forming native and non-native dimers, again confirmed by FTIR.^[12]^ ApoMb153 IBs display the strongest protection in hydrophobic helices and lower protection in regions that are less stable in native ApoMb153 and folding intermediates.^[45]^ FTIR shows notably high signal for α-helix/unstructured compared to other IBs, particularly ApoMb77, consistent with significant native-like helical content.

**Figure 7.**
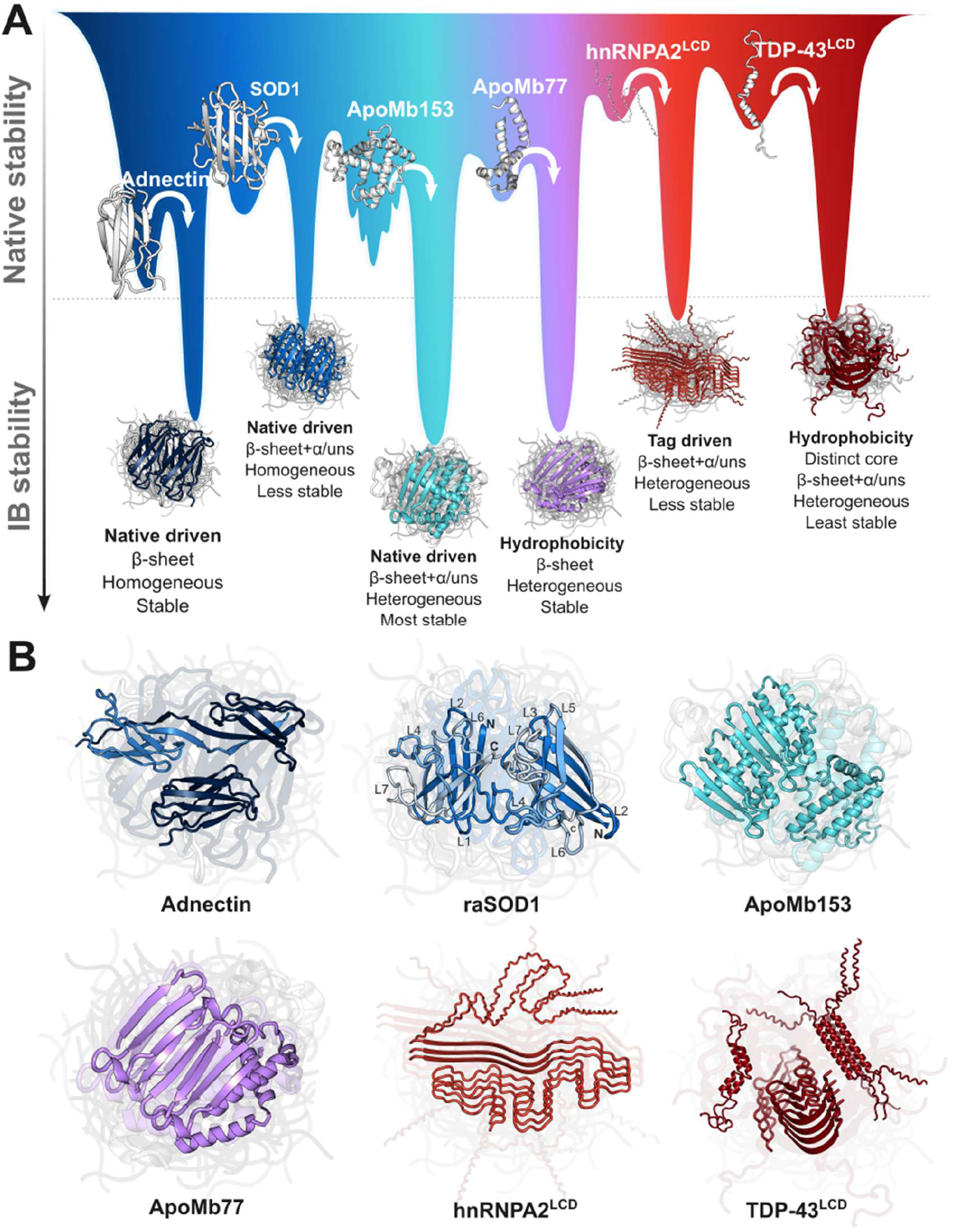
Schematic energy landscape for structure and stability of soluble monomer and IB. Corresponding specific experimental results from all methods are summarized in Table S1. A) *Top*: Schematic of ribbon structures and relative stabilities of protein monomers, from a very stable globular protein (left, Adnectin) to less stable monomers and increasing disordered regions (right, LCDs). Well width and roughness reflect structural heterogeneity and presence of folding intermediates. *Bottom*: Relative stabilities of IBs. The quantitative relationship between native protein stability and IB stability is unknown. Schematics summarize key features of IB architecture, with well depth representing IB stability from urea solubilization and amide protection, and well width representing structural heterogeneity inferred from qHDX, FTIR, and CR binding analyses. B) Summaries of IB properties, which depend on folding and aggregation pathways as well as stability and tertiary structure of the native state. Proteins that have well-structured native states, such as antiparallel β-folds (Adnectin, and to a lesser extent raSOD1, which includes long, flexible loops) and α-helical globin fold (ApoMb153) form IBs with native-like characteristics (protected amides and secondary structure in IBs resembling native). Native-like IB structures may include domain-swapped oligomers for Adnectin,^[33,34,79]^ that maintain native strands and protect loops (Figure S3),^[33,34]^ as well as nonnative dimers for raSOD1, among other species.^[38,80]^ ApoMb153 IBs are the most stable and heterogeneous, but with native-like features from intermediates^[45]^ and antiparallel β-sheet oligomers. As native stability decreases, IB organization shifts toward intrinsic sequence properties. ApoMb77 IBs are stable and heterogeneous, but their structure is driven by hydrophobicity. hnRNPA2^LCD^ and TDP-43^LCD^ IBs are less stable assemblies and include more localized amyloid-like β-cores arising from tag and hydrophobic helix-helix interactions, respectively. This schematic representation emphasizes that the complex IB architecture is strongly shaped by the native fold, sequence composition, and stability of the monomer.

As the stable native structure is further decreased in ApoMb77 and the LCDs, IB structure is increasingly driven by local characteristics of the primary sequence. ApoMb77, lacking the C-terminal half of the protein, shows increased protection of hydrophobic residues (Figure 3A) exposed by truncation and capable of burial within a β-oligomer/aggregates,^[24]^ while residues previously stabilized by the missing region may become less protected. The highly hydrophilic sequence of hnRNPA2^LCD^ confers generally low protection, which increases towards the N-terminal tag, a region with increased hydrophobicity and predicted β-structure/aggregation propensity (Figure 4). TDP-43_LCD_ also has extensive hydrophilic stretches, but these flank a central conserved hydrophobic nascent helical stretch (A321–M339); the highest protection is concentrated in a hydrophobic core, with relatively minimal protection elsewhere. Similarly, Chiti and coworkers have found for other proteins lacking a stable native conformation, such as Aβ, α-synuclein, and tau, that the regions most often forming the aggregate core have high hydrophobicity.^[47]^ Overall, the qHDX data obtained here complement and are consistent with results obtained using the other methods (Table S1), providing deeper molecular insight into the complexity of cellular aggregation. The general trend in these results is that when native-like conformations are more accessible (Adnectin, SOD1, ApoMb153), they are reflected in increased protection and structure in IBs, and as native disorder increases (from ApoMb77 to hnRNPA2^LCD^ and TDP-43^LCD^) so does disorder in IBs, as determined by intrinsic sequence properties.

Support for this native-based aggregation *in vivo* extends well beyond IBs.^[13]^ Aggregation of folded proteins involved in disease, such as transthyretin (TTR) in amyloidosis and crystallin in cataracts, as well as in the functional assembly of TasA bacterial biofilms, have also been reported to include native-like structure, despite initially being reported as dominantly amyloid.^[35,72,73]^ As noted, IBs have been mostly considered rather similar across different proteins, consisting of amorphous assemblies of folding intermediates or amyloid.^[13,22]^ The current results represent an important advance in defining the wide range of structures that can form and the rational basis thereof.

### qHDX NMR Bridges Static Structures and Dynamic Ensembles

An important result here is that amides in ordered amyloid-like assemblies determined by cryo-EM are not necessarily protected against exchange. This is particularly notable for TDP-43_LCD_, where strong amide protection in IBs is localized to the conserved hydrophobic region forming the core of multiple cryo-EM structures, while peripheral regions show much lower protection (Figures 5F, S6). In these cryo-EM structures, essentially all residues form intermolecular hydrogen bonds and, consequently, protection against exchange would be expected, even on the surface of a structure, as long as it is well ordered and the hydrogen bonds persist.^[74]^ These qHDX results indicate that the periphery is substantially less ordered than the core. Studies of polymorphic amyloid-β have likewise found that HDX can detect conformational populations differing from dominant cryo-EM fibril structures.^[75]^ These findings may have far-reaching significance, since the structures in the Amyloid Atlas, now numbering over 700 and mainly determined by cryo-EM, largely comprise highly ordered stacked polypeptides in which amides are very extensively hydrogen-bonded with the adjacent chains.

This apparent discrepancy likely reflects the distinct views of the aggregation landscape provided by cryo-EM and qHDX. Many high-resolution amyloid cryo-EM structures have reported highly ordered subsets of aggregation ensembles, typically corresponding to mature or end-stage fibrillar states.^[7,76]^ qHDX, however, reports on the entire sample, including partially ordered segments, flexible loops, and dynamic regions. Interestingly, for IBs the amides generally exhibit partial protection, uncovering conformational heterogeneity and structural dynamics. The partial protection of the globular proteins studied here, together with their native and unstructured FTIR signals, raises the intriguing possibility of oligomeric species retaining partially destabilized native folds.^[8]^ Likewise, solid-state NMR has identified oligomeric, heterogeneous and dynamic aggregate species.^[77,78]^ Owing to the intrinsic structural heterogeneity and polymorphism of protein aggregation, no single technique can fully capture aggregate complexity. qHDX is demonstrated here as a powerful complementary high-resolution tool for elucidating IB structure and the determinants of cellular aggregation.

## Conclusion

The results reported herein illuminate how IB structure varies widely between proteins and is determined by the properties of the soluble precursor protein. The studied IBs span a continuum from assemblies with prominent locally native-like structure (Adnectin, SOD1 and ApoMb153) to aggregates increasingly governed by local sequence features and hydrophobic interactions as native structure and stability decrease (ApoMb77, hnRNPA2^LCD^ and TDP-43^LCD^). In this context, qHDX provides residue-level measurements of protection across both ordered and dynamic regions. Integrating qHDX with FTIR, CR binding, proteolysis, urea solubilization, and sequence and structure analyses, this study establishes relationships between precursor-state properties and IB packing, stability and heterogeneity. These findings advance understanding of IB structure in contexts ranging from protein production to functional biomaterials and intracellular aggregation studies. Overall, this work identifies key determinants of aggregation *in vivo* and provides a framework for engineering aggregate structure and stability.

The ability to understand and control IB structure has important implications across biotechnology and materials science. Examples include designing efficient recovery of soluble protein from native-like IBs, controlled release of active protein from IBs, and the use of IBs as functional biomaterials, such as immobilized catalysts and biocompatible materials. As a specific case, functional IBs can retain substantial activity while enabling cost-efficient production and recyclability.^[1–3]^ Moreover, because IB formation is rapid and tunable (expression conditions, co-expression, or sequence variants), IBs provide a useful system for further defining the determinants of aggregation *in vivo* and their relationships to disease.^[1,7,13,14]^

## Supporting Information

## Acknowledgements

We would like to thank Andreas Barth for helpful discussions regarding FTIR and Silvia Cavagnero for guidance on apomyoglobin constructs. We also thank Sameer Al-Abdul-Wahid for technical assistance with NMR spectroscopy. This research was supported by funding from NSERC Discovery Grant RGPIN-2022-05139 to E.M.M., NSERC CREATE to B.S. and D.N., and OGS to P.R.C.

