## Supplementary Information for "Divergent Inclusion Body Structures and Stabilities Emerge from Native Monomer Properties"

### 1. Materials and Methods

#### 1.1. IB production and sample preparation

Experiments were performed as described previously,[1,2] with some modifications, as described below. Starter cultures for preparing IB samples were grown overnight at 37 °C in Luria Broth (LB) media (Table M1). 30 mL of starter culture was then centrifuged (5 min, 4000 × g), 20 mL of the supernatant discarded, cells resuspended in the remaining 10 mL and used to inoculate 1 L of minimal media (50 mM $Na_2HPO_4$, 25 mM $KH_2PO_4$, and 500 mg/L NaCl, 0.4 % Glucose, 100 μM $CaCl_2$, 2 mM $MgSO_4$, 500 mg/L $NH_4Cl$). For $^{15}N$, $^{13}C$ double-labelled samples, the media contained 0.3% $^{13}C$ glucose and 0.5 g/L $^{15}NH_4Cl$. The inoculated media were grown at 37 °C to an optical density at 600 nm (OD600) of 0.6, then protein expression was induced with isopropyl β-D-1-thiogalactopyranoside (IPTG, 1 mM) for 4 to 18 hours (overnight). The induction times are summarized in Table M1, along with cell lines, vector, and expression conditions. After induction, the cell culture was centrifuged at 5000 × g for 20 min, and the pellet was flash frozen in liquid nitrogen and stored at -80 °C.

Next, cell pellets were thawed and resuspended in 30 mL of TEN buffer (20 mM Tris, 1 mM EDTA, 100 mM NaCl, pH 8.0), then lysed using multiple freeze-thaw cycles (Table M1). Phenylmethylsulfonyl fluoride (PMSF, 1 mM) was also included during lysis for ApoMb, $hnRNPA2^{LCD}$ and $TDP\text{-}43^{LCD}$ to mitigate degradation of inclusion bodies by cellular proteases, while Adnectin and SOD1 were prepared as before without PMSF (Figure S10)[1,2]. After lysis, samples were centrifuged (21000 × g, 20 min). The pellet containing IBs was resuspended in 8 mL of TEN buffer, flash-frozen in liquid nitrogen as 1 mL aliquots, and stored at -80 °C.

For samples with low protein levels relative to the other systems (such as $hnRNPA2^{LCD}$ and $TDP\text{-}43^{LCD}$), a washing step for IBs was performed for qHDX NMR and FTIR as a control. This step removes the dominant contaminant protein, outer membrane protein (OMP). The 1 mL IB sample was centrifuged at 21000 × g for 20 minutes, and the pellet was resuspended in 3.3 mL of TEN buffer with 1% octyl glucoside (n-octyl-β-D-glucoside, nOG). The sample was incubated in the nOG solution for 4 minutes, and nOG was removed by centrifugation for 4 min at 9400 × g, with various fractions collected for SDS-PAGE. The final pellet was resuspended in 1 mL of TEN buffer. Successful removal of OMP can be seen in the gel Figure S11.

For NMR samples, 1-2 mL aliquots were used. After centrifugation for 20 minutes at 21000 × g, the pellet was resuspended in 5 mL of $H_2O$ or $D_2O$, flash frozen and then lyophilized for 48 hours. Next, the lyophilized IB was dissolved in DMSO-d6 with 25 mM DTT, 5% $H_2O$ and a volume of 5% DCA to reach pH 5.5 with a final volume of 600 μL.

**Table M1 – Summary of IB production and sample preparation conditions.**

| Protein | Cell line | Vector | Tag | Antibiotic | Extra nutrients | Expression conditions | Lysis conditions |
|---|---|---|---|---|---|---|---|
| **SOD1$^{A4V}$** | BL21(DE3) pLysS | pET21 | - | Chlor+Amp | Thiamine 0.1% | t4, 37 °C | 2x freeze-thaw, 40min with DNase 0.1mg/mL, 3x freeze-thaw |
| **Adnectin** | BL21(DE3) pLysS | pET-9d | 6xHis | Chlor+Kan | Metals$^{\times}$ | tON, 37 °C | 2x freeze-thaw, 20min with DNase 0.1mg/mL, 3x freeze-thaw (resuspend pellet right away) |
| **ApoMb** | BL21(DE3) pLysS | pET17b | - | Chlor+Amp | Thiamine 0.1% | t4, 37 °C | 3x freeze-thaw, 40 min with DNase 0.1mg/mL, 5x freeze-thaw$^{+}$ |
| **TDP-43$^{LCD}$** | Rosetta (DE3)* | pET28a | 6xHis | Chlor+Kan | | tON, 37 °C | 20min with lysozyme 0.1mg/mL and DNase 0.1mg/mL, 5mM $MgCl_2$, 5x freeze-thaw, 20 min incubation, 5x freeze-thaw$^{+}$ |
| **hnRNPA2$^{LCD}$** | Rosetta (DE3)* | pJ411 (addgene #118821) | 6xHis + TEV | Chlor+Kan | Metals$^{\times}$ and Vitamins$^{\times}$ | t6, 37 °C | |

*Unlike BL21(DE3) pLysS cells, Rosetta(DE3) cells do not carry the pLysS plasmid and so do not constitutively express T7 lysozyme, which makes the BL21(DE3) pLysS cells lyse more easily. For this reason, for samples derived from Rosetta(DE3) lysozyme was added during lysis at a final concentration of 0.1 mg/mL.

$^{+}$ApoMb, TDP-43$^{LCD}$, and hnRNPA2$^{LCD}$ samples were subjected to additional freeze–thaw cycles compared to SOD1 and Adnectin. To assess whether the increased number of cycles affected the samples, IBs with a higher number of freeze–thaw cycles were compared with samples prepared using fewer freeze–thaw cycles, as described in SI R&D 3: Lysis results.

$^{\times}$Metals: 3 µM $(NH_4)_6Mo_7O_{24}$, 400 µM $H_3BO_3$, 30 µM $CoCl_2$, 10 µM $CuSO_4$, 80 µM $MnCl_2$, 10 µM $ZnCl_2$

$^{\times}$Vitamins: 1µg/mL Biotin, 0.4 µg/mL Choline Chloride, 0.5 µg/mL Folic acid, 0.05 µg/mL Riboflavin, 0.5 µg/mL Pantothenic acid, 0.5 µg/mL Thiamine, 0.5 µg/mL Niacinamide, and 1 µg/mL Myo-inositol.

#### 1.2. Primary amino acid sequences of proteins studied

Adnectin:
GVSDVPRDLEVVAATPTSLLISWSARLKVARYYRITYGETGGNSPVQEFTVPKNVYTATI
SGLKPGVDYTITVYAVTLLRDYGPISINYRTEIDKPSQHHHHHH

$SOD1^{A4V}$:
ATKVVCVLKGDGPVQGIINFEQKESNGPVKVWGSIKGLTEGLHGFHVHEFGDNTAGCTS
AGPHFNPLSRKHGGPKDEERHVGDLGNVTADKDGVADVSIEDSVISLSGDHCIIGRTLV
VHEKADDLGKGGNEESTKTGNAGSRLACGVIGIAQ

ApoMb 153:

MVLSEGEWQLVLHVWAKVEADVAGHGQDILIRLFKSHPETLEKFDRFKHLKTEAEMKA
SEDLKKHGVTVLTALGAILKKKGHHEAELKPLAQSHATKHKIPIKYLEFISEAIIHVLHSR
HPGNFGADAQGAMNKALELFRKDIAAKYKELGYQG

ApoMb 77:

MVLSEGEWQLVLHVWAKVEADVAGHGQDILIRLFKSHPETLEKFDRFKHLKTEAEMKA
SEDLKKHGVTVLTALGAILK

$TDP\text{-}43^{LCD}$:

GGFGNQGGFGNSRGGGAGLGNNQGSNMGGGMNFGAFSINPAMMAAAQAALQSSWG
MMGMLASQQNQSGPSGNNQNQGNMQHHHHHH

$hnRNPA2^{LCD}$:

MGSDKIHHHHHHENLYFQGHMNQGGGYGGGYDNYGGGNYGSGNYNDFGNYNQQPS
NYGPMKSGNFGGSRNMGGPYGGGNYGPGGSGGSGGYGGRSRY

#### 1.3. qHDX NMR

Experiments were performed as described previously,[1,2] with some modifications, as described below.

##### *1.3.1. Sequence Specific Resonance Assignments*

Standard NMR experiments based on standard pulse sequences and parameter sets were acquired on a Bruker Avance 600 MHz or 700 MHz (with cryoprobe) NMR spectrometer: HNCO, HN(CA)CO, HNCACB, CBCA(CO)NH, and HN(CA)NNH.[3–7] Computer aided resonance assignment (CARA) was used for making assignments.[8]

##### *1.3.2. qHDX NMR sample preparation and experiment*

Each qHDX experiment requires 2 samples, one protonated and a corresponding sample that has undergone exchange with $D_2O$. For each sample, 1 mL aliquot of cell pellet was quickly thawed and then centrifuged (21000 × g, 20 min), the supernatant was removed, and the pellet was resuspended either in 5mL of $H_2O$ (protonated sample) or 5 mL of $D_2O$ (exchanged sample). After allowing 1 hour for exchange at room temperature, samples were flash frozen in liquid nitrogen and then lyophilized for 48 hours.

For the protonated (exchanged) experiment, the lyophilized samples were dissolved in 0.5 mL of buffer solution containing 95% DMSO-d6, 5% $H_2O$ ($D_2O$), 25 mM DTT, and a volume of 5% DCA (in DMSO) to achieve a final pH of 5.5 (pD=5.5=pHread + 0.4). The volume of DCA was predetermined using an unlabelled sample prepared in parallel with the $^{15}N$-labelled sample. This procedure allowed for an experimental dead time for sample measurement by NMR of under 10 minutes.

All qHDX NMR experiments were conducted at a temperature of 19 °C (292 K) using a Bruker Avance 600 MHz or 700 MHz spectrometer locked to DMSO-d6. For the exchanged samples, a $^1H$ 1D NMR spectrum was acquired with 8 scans, for a total acquisition time of 1 min 40 s. This was followed by a series of nine consecutive $^1H$-$^{15}N$ HSQC spectra, each with an acquisition time of 20 min. For ApoMb, TDP-43$^{LCD}$ and hnRNPA2$^{LCD}$ IB samples, each HSQC spectrum was acquired using 4 scans and 256 increments in the indirect $^{15}N$ dimension, with a spectral width of 23 ppm. For Adnectin and SOD1 samples, each HSQC spectrum was acquired using 8 scans and 128 increments in the indirect $^{15}N$ dimension, with a spectral width of 44 ppm.[1,2] The sample was then kept at room temperature for a week. After that, $^1H$-$^{15}N$ HSQC spectra were taken to measure the equilibrium H-D sample signal levels. For a given protein, qHDX exchange experiments were performed at least in duplicate, and the reported protection is the average of the two independent culture growths (i.e. biological replicates).

##### *1.3.3. qHDX NMR analysis*

Spectra were processed with Topspin software (Bruker) and analyzed with CCPNMR software (V2.4) to measure amide cross-peak intensities. First, the ratio (R) of the intensity of the $^1H$ 1D amide region (8.1-8.5 ppm) of the $D_2O$ sample divided by the intensity of the corresponding $H_2O$ sample (from the same growth) was determined, where these $^1H$ spectra were scaled for protein concentration using the methyl region (0.5-0.95 ppm). Second, cross-peak decays were fit to an exponential equation (Eq 1) where $Signal\ Intensity_i$ is the intensity for cross-peak $i$, at time $t$ since dissolution of the sample in DMSO, $A$ is the amplitude of the decay, $k_{int}$ is the intrinsic rate constant of exchange for amide $i$ in DMSO, and $C$ is the offset value of the fit to obtain smoothed, fitted values for peak intensity for each amide in the first $^1H$-$^{15}N$ HSQC spectrum corresponding to 20 minutes (t=20) or corresponding to the time of sample dissolution in DMSO (t=0).[1,2]

$$Signal\ Intensity_i = A \times e^{(-k_{int}\,t)} + C \qquad \text{Eq 1}$$

Next, signal ($Peak\ Intensity_i$) was normalized for concentration of the protein in the NMR samples by scaling to the average intensity of all peaks ($Average\ Intensity_{All}$), both for protonated (Eq 2) and exchanged (qHDX) peaks (Eq 3):

$$Protonated\ Intensity_i = \frac{Peak\ Intensity_i}{Average\ Intensity_{All}} \qquad \text{Eq 2}$$

$$qHDX\ Intensity_i = \frac{Smoothed\ Intensity_i}{Average\ Intensity_{All}} \qquad \text{Eq 3}$$

where *Smoothed Intensity$_i$* is the fitted intensity at t=20 or t=0. Lastly, the fraction amide protection for each peak was obtained by combining the smoothed and scaled values:

$$Fraction\ Protected_i = R \times \frac{qHDX\ Intensity_i}{Average\ Protonated\ Intensity_i} \quad \text{Eq 4}$$

where *Average Protonated Intensity$_i$* is the average of *Protonated Intensity$_i$* across all samples (at least two biological replicates) and *R* is the average ratio of the intensity of the $^1$H 1D amide region (8.1-8.5 ppm) of $D_2O$ sample divided by the intensity of the corresponding $H_2O$ sample (from the same growth), where these $^1$H 1D spectra were scaled for protein concentration using the most upfield peak in the methyl region (~0.81 ppm). The average ratios, R, for the proteins studied here are shown in Table S1 and Figure 6B.

#### 1.4. Mass spectrometry

Limited proteolysis has been used extensively as a probe for protein structure. Here, we show that proteolysis by cellular proteases can provide valuable insights into IB structures. Freeze-thaw lysis is used for IB preparation (production and isolation), and PMSF plays an important role in this process. Without adding PMSF, clear degradation of IBs is observed on the gel for ApoMb, hnRNPA2$^{LCD}$, and TDP-43$^{LCD}$ IBs (Figure S10). Matrix-assisted laser desorption/ionization time-of-flight (MALDI-TOF) mass spectrometry (MS) was used to identify possible fragments. To facilitate sequence assignment, parallel samples of one unlabeled and another one $^{15}$N-labelled were analyzed. The expected mass shift between the unlabeled and $^{15}$N-labelled spectra supported the identification of fragment sequences by comparing the mass differences of corresponding peptide peaks. As the different fragments contain distinct amino acid compositions and therefore different numbers of nitrogen atoms, the fragment identification relied on both mass and the number of nitrogen atoms.

To enable MALDI-TOF analysis, IB samples were diluted 25-fold in MQ water. The samples were then mixed with freshly prepared saturated α-cyano-4-hydroxycinnamic acid (HCCA) matrix in acetonitrile/$H_2O$+0.1% trifluoroacetic acid diluted 1:1 with acetonitrile/$H_2O$+0.1% trifluoroacetic acid solution. After fully mixing, 1 μL of the mixture was spotted on the stainless-steel plate for MALDI analysis. Linear positive ion mode with a detection range of 2-20 kDa was used for the data acquisition. The calibration was done using $[(CsI^3)_nCs]^+$ cluster ion series, generated using freshly prepared $CsI_3$ and 2-[(2E)-3-(4-tert-butylphenyl)-2-methylprop-2-enylidene] malononitrile (DCTB) in tetrahydrofuran (THF). A range of 2-13 kDa can be calibrated with m/z accuracy <10ppm. For each IB sample, a sum of 5000 laser shots was obtained. The MS spectrum was generated and analyzed on *flexAnalysis 3.4 (Bruker)*. Using scan MS results, a list of potential fragment sequences was exported. The calculated theoretical $^{15}$N-labelled masses were compared with $^{15}$N-labelled experimental data, and fragments corresponding to the m/z values were identified.

#### 1.5. Bioinformatics analysis

Various biophysical quantities of the IB proteins were analyzed using bioinformatics tools. The hydrophilicity and hydrophobicity of the proteins were assessed using the Kyte and Doolittle scale, with a window of 7 amino acids.[9] Sequence-based predictors, such as ZipperDB,[10] PASTA,[11]

Aggrescan,[12] and Tango,[13] were used to identify aggregation-prone regions (APRs). To facilitate a comparison among the predictors, the outputs of PASTA, Aggrescan, and Tango were normalized to a scale of 0 to 1. For ZipperDB, as the values are energy-based (kcal/mol), they were plotted from values of -23 to -26, with values below -24 considered indicative of aggregation propensity. For SOD1, 6 fibril structures (7VZF, 9JBO, 8IHV, 8IHU, 9IYD and 9IYJ) were used to estimate the percentage in β-conformation. Similarly, for TDP43 LCD, 14 structures found in the Amyloid Atlas (6N37, 6N3B,6N3A, 7KWZ, 8QX9, 8QXA, 8QXB, 7PY2, 8CG3, 8CGG, 8CGH, 7Q3U, 9FOR and 9FOF) were used.[14] In the case of hnRNPA2, because only one experimental structure is available (6WQK), AlphaFold 3 multimer was used to predict a potential structure as well.[15]

#### 1.6. ATR-FTIR Spectroscopy

Secondary structure of IBs was measured with attenuated total reflectance Fourier transform infrared spectroscopy (ATR-FTIR) using a Tensor 37 FTIR spectrometer with LN-MCT detector and Bio-ATRII cell with ZnSe crystal (Bruker Optics). Measurements consisted of acquiring four spectra successively, each with 256 scans in the spectral range of 4000-1200 $cm^{-1}$ and a resolution of 4 $cm^{-1}$; the last of these spectra was used for quantitative analysis (measured after 5 minutes in the cell for consistent thermal equilibration). A Thermo HAAKDC water bath (Haake) was used to control the temperature at 25 °C.

Before adding the IB sample, background (air) and buffer (TEN) readings were acquired. After taking initial spectra, 25 µL of IB was loaded onto the sample cell and measured as described above. After removing the IB sample, the crystal was washed using 3 x 25 µL of TEN buffer (drawing and ejecting solution from the pipette 20 times for each 25 µL). Then 25 µL of TEN buffer was loaded, and post-sample spectrum was collected and used to subtract the contribution of material that may have been altered due to strong adsorption to the crystal.[16] To wash the crystal in between IB samples, it was soaked in 25 µL of 5% SDS for 5 minutes and then fully washed with water.

#### 1.7. FTIR Spectral Analysis

FTIR spectra were processed and analyzed as before using the OPUS 6.5 software (Bruker) for atmospheric compensation and buffer subtraction.[1,2] Baseline correction was then applied in Origin 2025 by subtracting a straight line between 1700 $cm^{-1}$ and 1600 $cm^{-1}$, and each spectrum was normalized to the area of the amide I band. The final average spectrum for each IB sample was calculated by averaging at least four replicate measurements (2 separate aliquots of a given sample, i.e. technical replicates; for at least 2 different growths, i.e. biological replicates). For second derivative analysis, the amide I band was normalized to 1. The second derivative was calculated using Savitzky-Golay filter for smoothing with order 2 and a window of 9 points. Peak-deconvolution was performed using a Voigt function. For a given spectrum, initial peak position estimates for the fitting were obtained based on the corresponding second derivative spectrum. The Gaussian width was limited to a maximum of 30 $cm^{-1}$ to keep the components within a physically reasonable range and the setting to 3 components avoids overfitting.[17] Fits had an $R^2$ greater than 0.99. To evaluate possible background contribution, a sample background correction was performed as described in 3.1 (SI R&D 1). For these corrections, spectra after normalization to the area of amide I were used.

#### 1.8. Congo Red

A 300 μM Congo red (CR) stock solution was prepared in TEN buffer containing 10% ethanol and filtered three times using a 0.2 μm Acrodisc filter.[2,18] CR-IB assays followed a protocol adapted from Klunk *et al.*[19] Absorbance measurements were taken with a Cary 50 spectrophotometer (300 nm - 700 nm) using TEN buffer as a blank. Each sample involved diluting the IB to obtain a maximum absorbance below 1, resulting in IB concentrations averaging around ~20 μg/mL. As controls, 20 μM CR and diluted IB were measured separately before each binding experiment. Each diluted IB was mixed with 300 μM CR to reach a final concentration of 20 μM CR and incubated for 10 minutes before being scanned three times. The plots represent the IB+CR minus the IB to identify changes in spectral shape and intensity (CR bound protein spectrum = IB+CR - (diluted IB)). Sonicated SOD1 was utilized as an amyloid control.[18]

#### 1.9. Chemical denaturation

Urea solubilization is an established method for measuring the apparent stability of IBs and may also report on their heterogeneity.[20–22] This method treats IB protein as existing in one of two states: an aggregated state that scatters light (measured at 350 nm, $A_{350nm}$) and a soluble state that does not contribute to light scattering. Data fitting (details below) provides apparent $C_{mid}$ value reflecting the denaturant concentration at which half of the IBs are solubilized, with a higher $C_{mid}$ indicating greater resistance to solubilization. This parameter has been used previously to report differential IB stability in response to changes in growth temperature,[20,21] and is applied here to compare stability across the different protein model systems examined in this study.

The data fitting also provides an *m* value, which describes the steepness of the solubilization profile and is proposed here as an indicator of IB structural heterogeneity. In the protein folding literature, the *m* value is commonly interpreted as reflecting the denaturant-dependence of the unfolding free energy, which is related to the change in solvent-accessible surface area upon unfolding.[23,24] Relatively low apparent *m* values have been associated with less cooperative or multistate transitions.[25] In the current study, a higher *m* value suggests a comparatively more homogeneous IB population that solubilizes within a narrow range of denaturant concentrations, whereas a lower *m* value indicates a broader solubilization profile, consistent with a heterogeneous population of aggregated species with varying resistance to solubilization.

Urea stock solution (9-10 M) was prepared in Trizma buffer (20 mM, pH 8.1) and diluted to standard concentrations ranging from 0-7 M. Generally, IB samples were first diluted 50-fold using TEN buffer and initial optical density measured at 350 nm using a Cary 500 Scan UV-Vis Spectrophotometer (Varian). Based on this $OD_{350nm}$, samples were diluted with TEN buffer to ensure similar absorbance readings for samples measured using a 96-well plate. Then, 40 μL of diluted IB was plated in duplicates, and 160 μL of standard urea solution was added. IB solubilization was monitored using a SpectraMax Plus 384 microplate reader (Molecular Devices) at 350 nm and 27 °C. The plate was shaken for 3s before the first read (for 40s), and 5s between each subsequent read. Kinetic readings were measured continuously for 3 hours from the time of plating. For data analysis, for each urea concentration, the last 20 minutes of absorbance readings (which changed relatively slowly) were averaged. The fractional solubilization values for all urea concentrations were obtained by normalizing absorbance values with respect to the corresponding first reading for 0 M urea. The average solubilization profile for each IB was calculated using at least 2 independent sample growths, each measured in duplicate (Figure 6G). Background scattering was subtracted using measurements of corresponding IBs treated with proteinase K for 24 hr and then solubilized overnight. For these background samples, the amount of insoluble material measured by turbidity correlates with that measured by SDS-PAGE (Figure S9).

The background corrected fractional solubilization values were fit to Eq. 5 for a two-state model using Origin2018Pro to obtain apparent $C_{mid}$ and *m* values based on the urea-dependence of Y with limiting optical signal for the aggregated (A) or denatured (D, i.e. solubilized) states, respectively (Figure 6H, I).[20–22]

$$Y = \frac{Y_A-(Y_A-Y_D)*(e^{\frac{-(C_{mid}*m)+m[urea]}{RT}})}{1+e^{\frac{-(C_{mid}*m)+m[urea]}{RT}}} \qquad \text{Eq.5}$$

For measuring the extent of IB solubilization, samples were incubated 24 hours in urea and then centrifuged (21000 × g, 20 min). Pellets were visualized using SDS-PAGE (Figure S9).

#### 1.10. SDS-PAGE

Sodium dodecyl sulfate–polyacrylamide gel electrophoresis (SDS-PAGE) was used to visualize IB formation, proteolysis during IB isolation, protein composition in IB samples, washing efficiency, and IB stability in urea. Two gel systems were used: 15% SDS-PAGE and 16% tricine-SDS-PAGE. The 16% tricine-SDS-PAGE was used to better resolve smaller proteins and peptide fragments. The 15% SDS-PAGE gels were used to visualize urea solubilization of Adnectin, raSOD1$^{A4V}$, and ApoMb153, as well as Adnectin IB washing (Figure S9G–I and Figure S11A). All other gels were 16% tricine-SDS-PAGE (Figures S9J–L, S10, S11B–D, and Figure S12).

### 2. Supplementary Results

The $^{1}H$-$^{15}N$ HSQC spectra for the IB proteins solubilized in DMSO (Figure S1) provide many well resolved cross-peaks for individual amide groups, allowing for high-resolution qHDX analysis of IB structure.

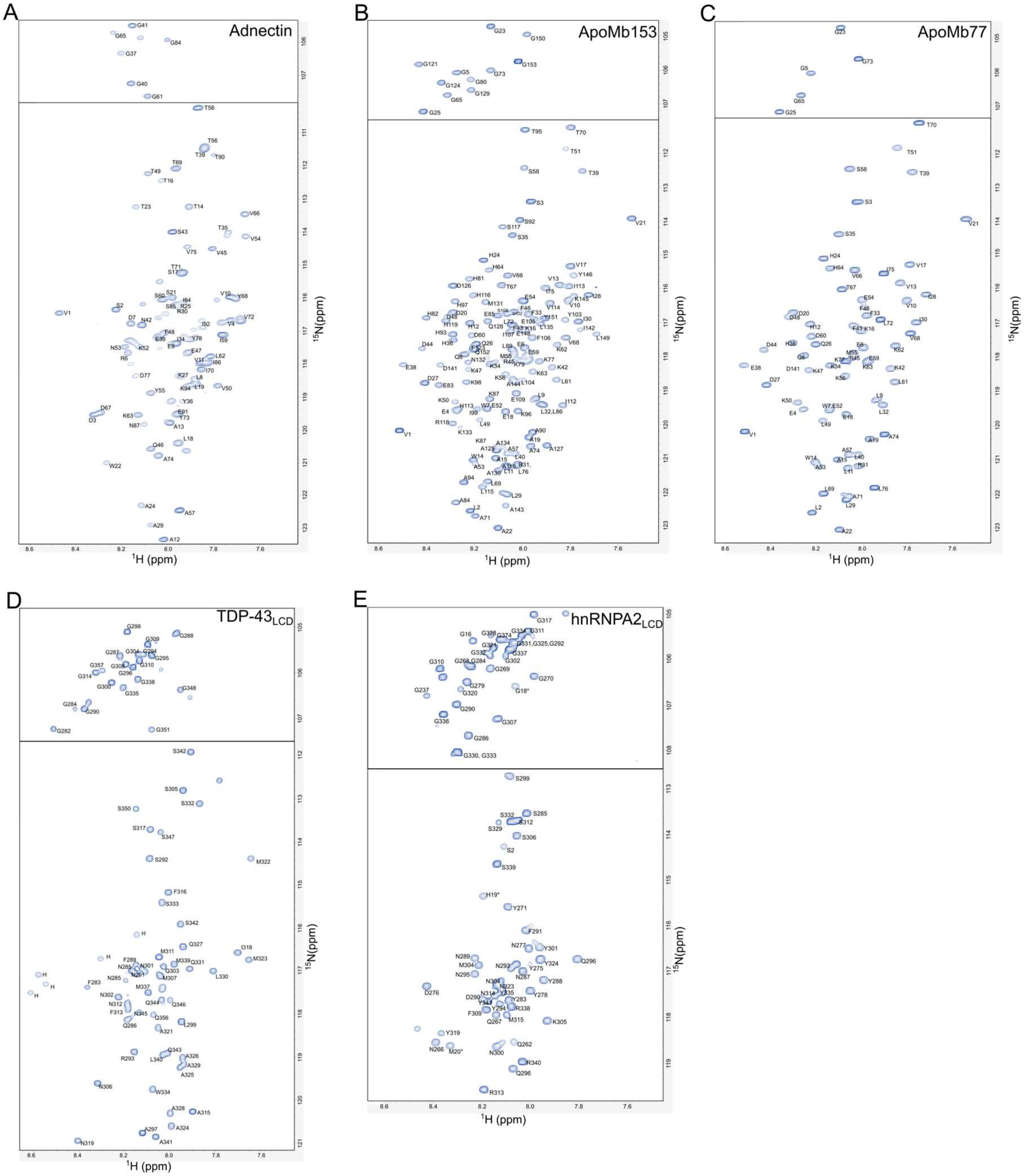

**Figure S1.** $^{1}H$–$^{15}N$ HSQC spectra of IBs denatured in 95% DMSO, 5% $H_2O$ with 25 mM DTT. Spectra are shown for: A) Adnectin, B) ApoMb153, C) ApoMb77, D) TDP-43$^{LCD}$, and E) hnRNPA2$^{LCD}$. Cross-peaks are labelled by residue, and resonance assignments are summarized in Tables S2-S6. Details for SOD1 have been reported previously.[2]

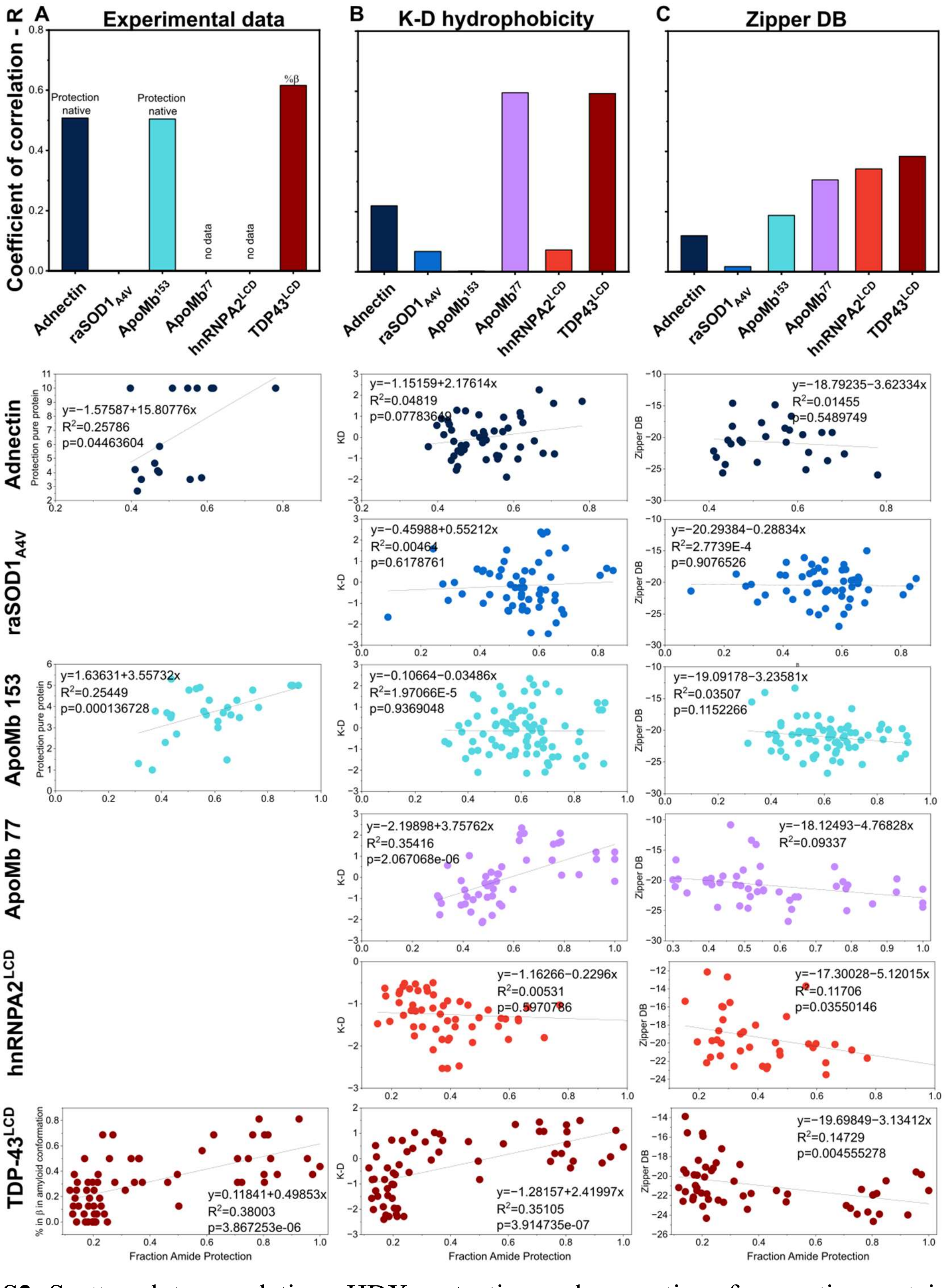

**Figure S2.** Scatter plots correlating qHDX protection and properties of respective proteins. The qHDX protection (at t=0) is on the x-axis, and experimental data or property is on the y-axis. P-values were calculated using R. A) Left column corresponds to experimental property, as available: amide exchange rate in pure protein folded monomer in solution (Adnectin),[26] and ApoMb153,[27] or Fraction in β-conformation in amyloid structures (6N37, 6N3B,6N3A, 7KWZ, 8QX9, 8QXA, 8QXB, 7PY2, 8CG3, 8CGG, 8CGH, 7Q3U, 9FOR and 9FOF). B) Middle column corresponds to Kyte-Doolittle hydrophobicity.[9] C) Right is the aggregation predictor Zipper DB.[10]

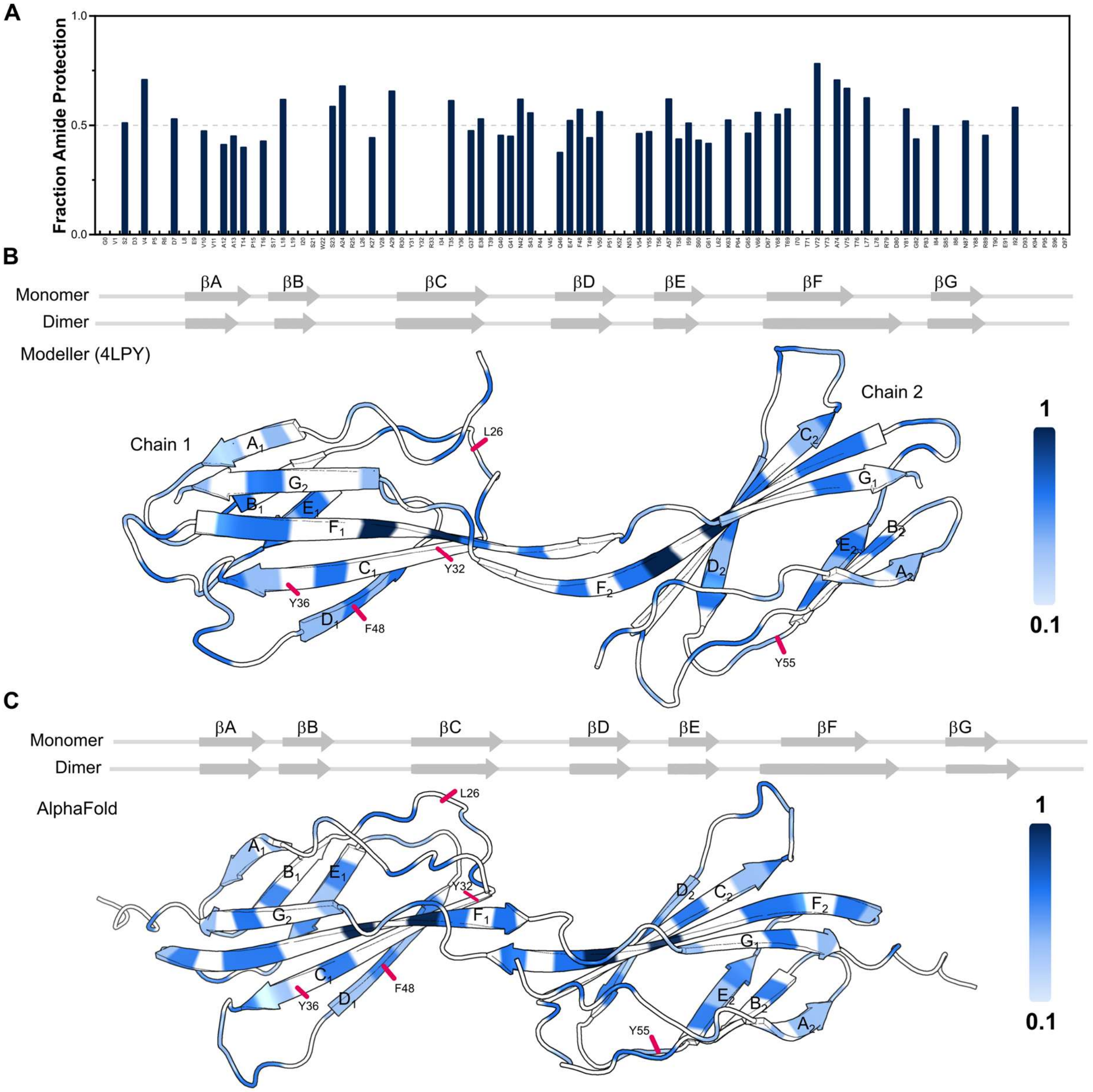

**Figure S3.** Adnectin analysis in the context of domain-swapped model structures. qHDX and proteolysis data are mapped onto strand G-swapped Adnectin dimers predicted using Modeller[28] and AlphaFold3[15]. A) Fraction of protection against solvent exchange. Each bar is an average of three biological replicates. Gaps indicate no data. B) Adnectin monomer structure predicted by AlphaFold3 and a domain-swapped dimer model generated using Modeller based on the crystal structure of an Adnectin homologue (PDB 4LPY). Protection values from 0.1 to1.0 are mapped onto the domain-swapped model and coloured from light to dark blue. C) AlphaFold3 models of the Adnectin monomer and domain-swapped dimer with protection mapped as in panel B. Red lines indicate cleavage sites identified by MS (Figure S4). The mutual proximity of multiple sites is consistent with destabilization in this region in the IB and consequent accessibility to cellular proteases during sample preparation.

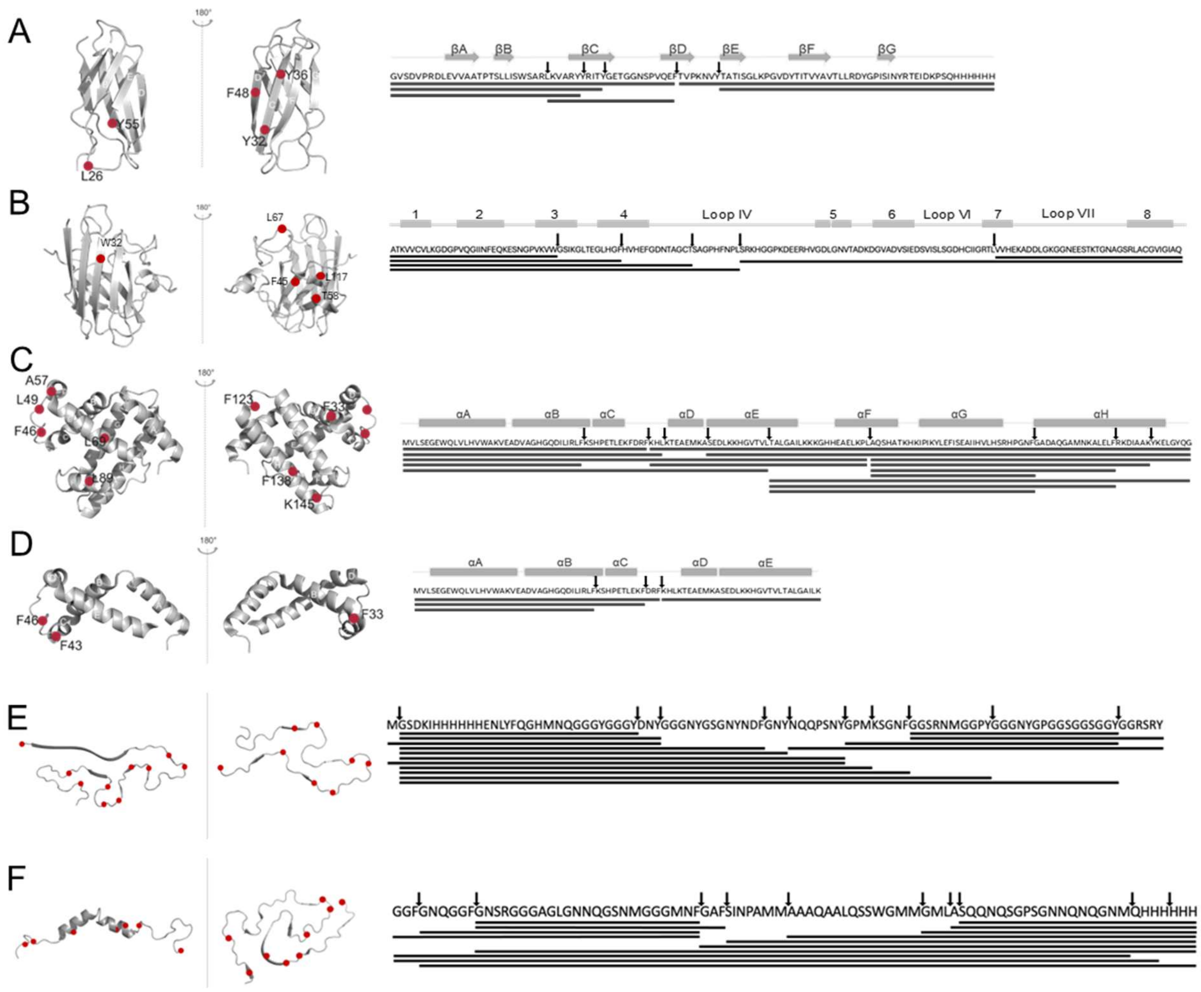

**Figure S4.** Mass spectrometry analysis reveals exposed proteolysis sites. Proteolysis sites (red dots on ribbon representations of native monomers at left and at right corresponding arrows and fragments shown as black bars along with primary sequences and secondary structure elements) caused by cellular proteases during IB sample preparation without protease inhibitor PMSF (see Methods 1.1). A) Adnectin B) raSOD A4V (2GBU), C) ApoMb153 (1VXF), D) ApoMb77 (1VXF corresponding length), E) hnRNPA2$^{LCD}$ (AlphaFold3 was used to predict native structures for hnRNPA2$^{LCD}$,[15] while fibril structure corresponds to 6WQK) F) TDP-43$^{LCD}$ (2N3X, 7PY2). Representative MALDI-TOF raw data in panels G-L illustrate: G) ApoMb 77 IBs prepared without PMSF. Black arrows show ApoMb 77 fragments resulting from cellular proteases. H) ApoMb 77 IBs prepared with 1 mM PMSF. I) TDP-43$^{LCD}$ IBs prepared without PMSF. J) TDP-43$^{LCD}$ IBs prepared with 1 mM PMSF. Data for Adnectin IBs K) unlabelled, fragment sequences corresponding to representative high intensity peaks are given on the spectrum, and L) $^{15}$N-labelled, prepared without PMSF. The given mass shifts from unlabelled to $^{15}$N-labelled correspond to the number of nitrogen atoms in a given peptide. Peaks for undigested monomers are marked with * for singly charged and ** for doubly charged.

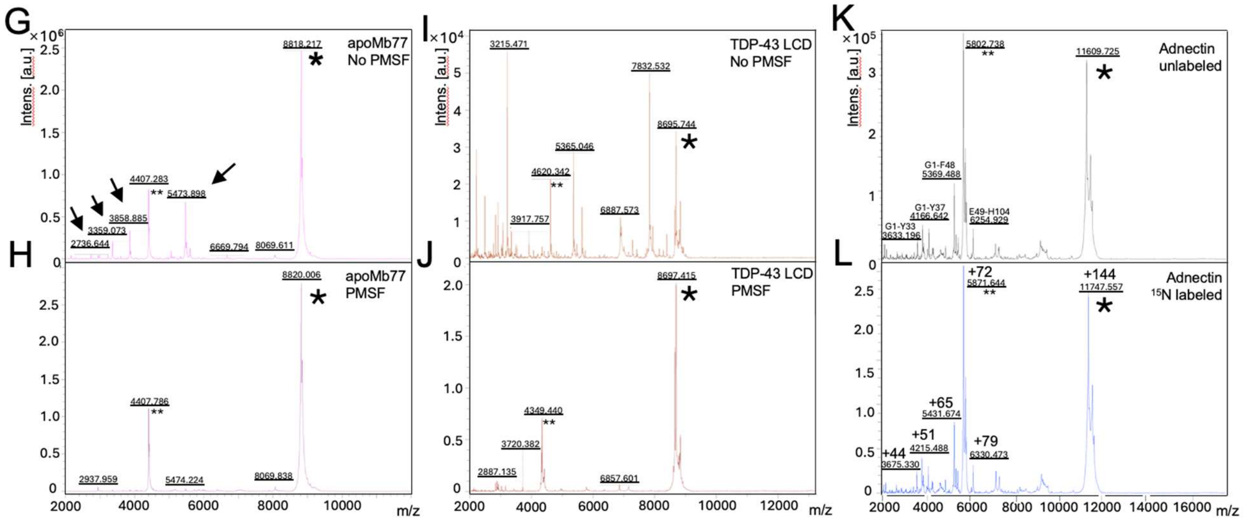

**Figure S4.** (continued)

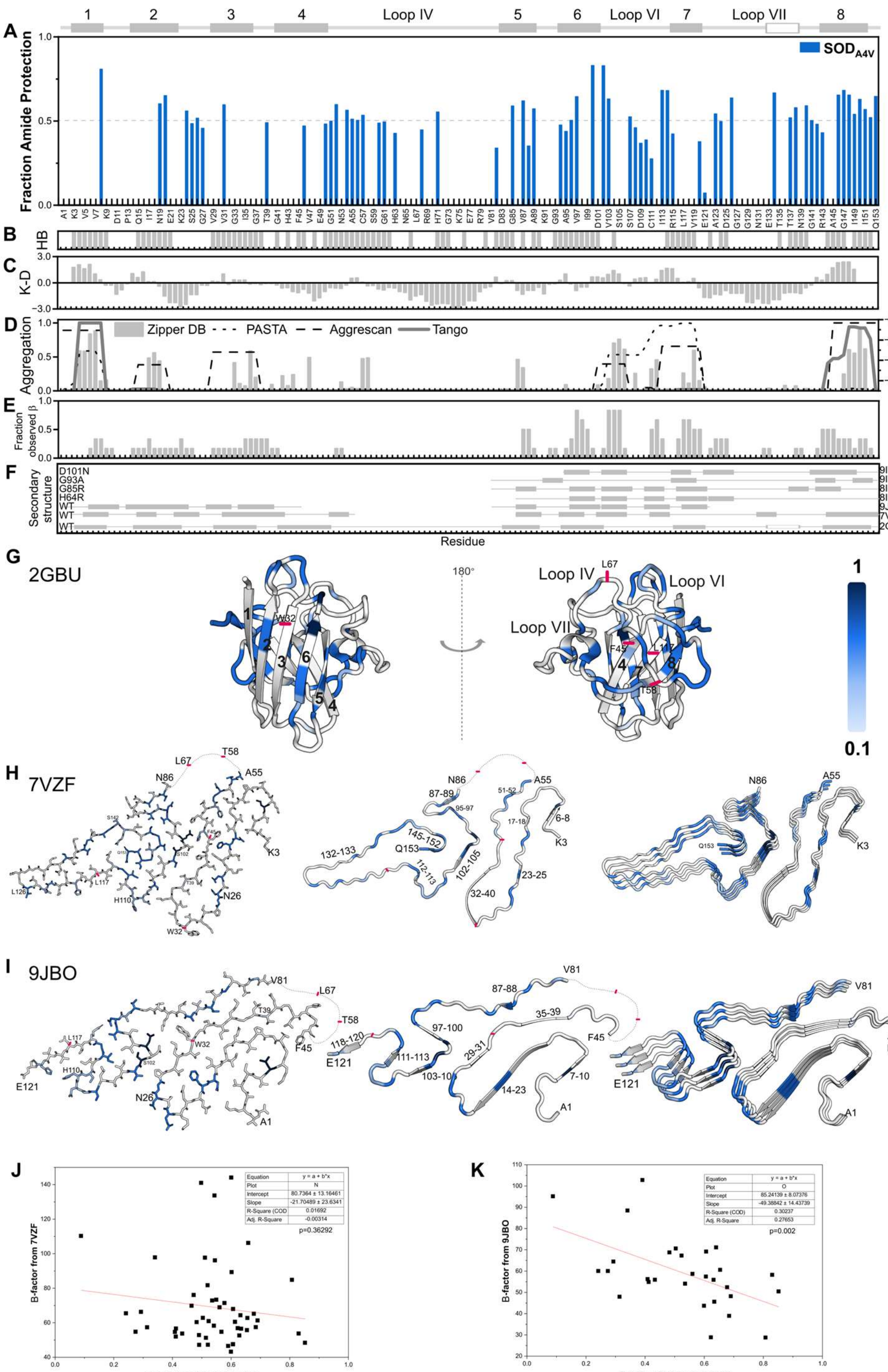

**Figure S5.** qHDX analysis of ALS-associated SOD1 A4V from Naser et al. 2022.[2] A) Fraction of amide protected against solvent exchange for SOD1 A4V. Bars are average values of three biological replicates. B) WT holo SOD1 native amide HDX protection.[29] Residues that are protected against exchange in the mature dimer are indicated by grey bars. C) Kyte & Doolittle hydrophobicity, positive values are hydrophobic, negative are hydrophilic.[9] D) Sequence-based aggregation predictors: TANGO,[13] AGGRESCAN,[12] and PASTA,[11] normalized from 0 (no aggregation propensity) to 1 (highest aggregation propensity); and ZipperDB.[10] E) Fraction observed β-conformation in available SOD1 amyloid fibril structures, formed *in vitro* from full-length protein as determined by cryo-EM: WT: 7VZF, 9JBO, and ALS-associated SOD1 mutants: H46R, 8IHV; G85R, 8IHU; G93A, 9IYD; and D101N, 9IYJ. Extensive portions of the polypeptide chain had no defined structure, as illustrated in F) which shows secondary structure in fibril structures for WT and mutant SOD1s, as well as the secondary structure from the crystal structure of reduced apoSOD1 dimer (PDB 2GBU). G) qHDX protection, mapped onto the crystal structure of reduced apoSOD1 dimer (PDB 2GBU), from 0.1 to 1, coloured as light to dark blue, respectively.[30] qHDX protection mapped onto cryo-EM structures of amyloid fibril of SOD1 prepared from full length WT SOD1 *in vitro* with shaking, starting with H) reduced apo protein, 7VZF,[31] or I) enzymatically active protein under reducing and metal depriving conditions, 9JBO.[32] White indicates no data for G-I. The amyloid structures are very different and many residues that show qHDX protection are not ordered in the structures. J) Correlation between the B-factor in the fibril (7VZF) and A4V protection is not significant. K) Correlation between the B-factor observed in the fibril (9JBO) and A4V protection is significant and provides additional support for the presence of local native-like features in amyloid, as also evidenced by notable correspondence between native and fibril β-strands, shown in F), G) and I).

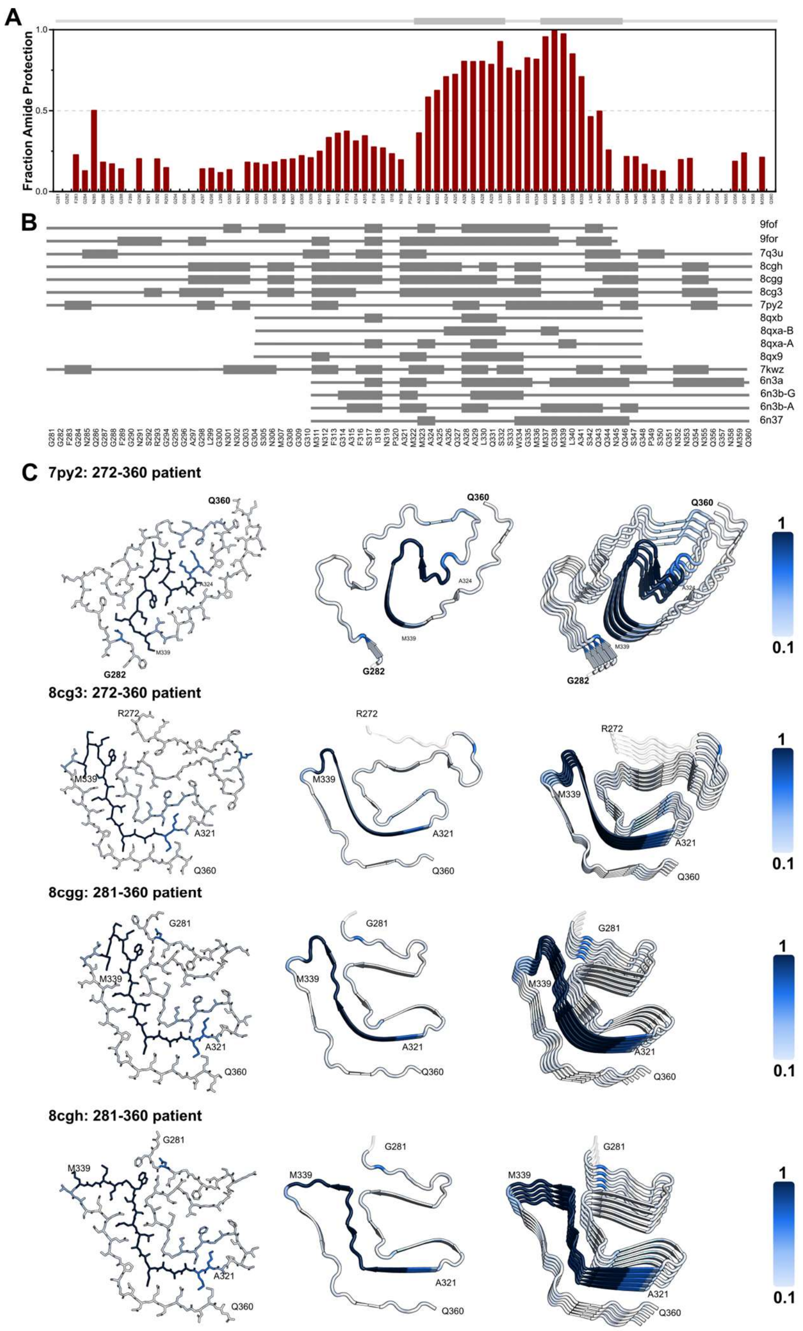

**Figure S6.** Comparison of qHDX results for TDP-43 LCD (281-360) IB to available structures of TDP-43$^{LCD}$. A) Fraction amide protection for TDP-43$^{LCD}$. Bars are average values of three biological replicates. Gaps indicate areas of no data. Grey line at top of panel represents secondary structure from the NMR solution structure (2N3X) with thin line for random coil and thick line for helices. B) Regions in β-conformation in cryo-EM structures of TDP are shown by thick grey bars. C) Cryo-EM structures of TDP-43 assemblies from patients: 8CG3, 8CGG, 8CGH, 9FOR and 9FOF or D) formed *in vitro*: 6N37, 6N3B, 6N3A, 7KWZ, 7Q3U, 8QX9, coloured according to qHDX protection. Fractional protection from 0.1 to 1 coloured as light to dark blue, respectively; unassigned residues are light grey, and structured regions of fibrils containing residues not included in the construct studied here are transparent. The residues with the strongest protection tend to be the most localized in the middle of the structure in the assemblies from patients.

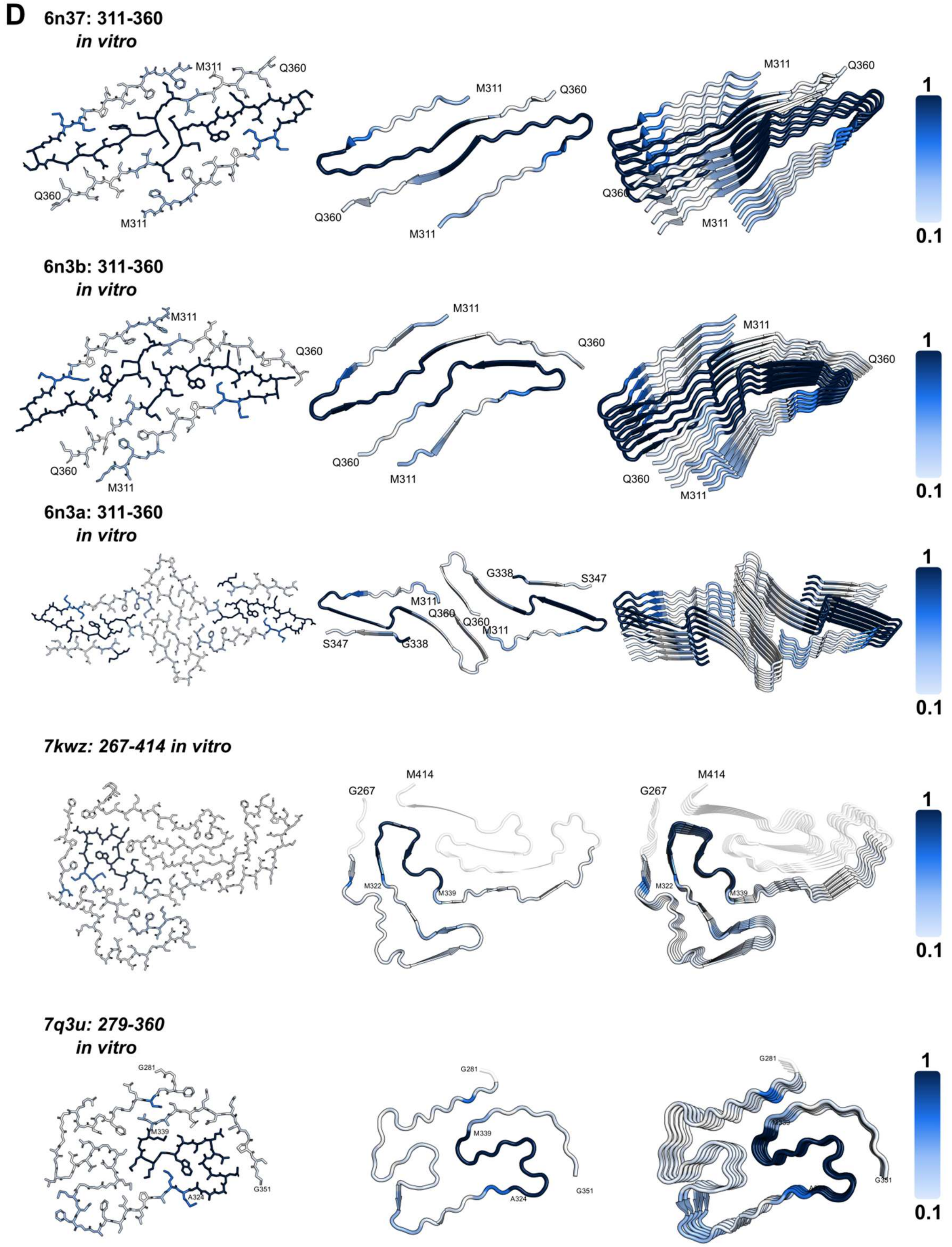

**Figure S6.** (continued)

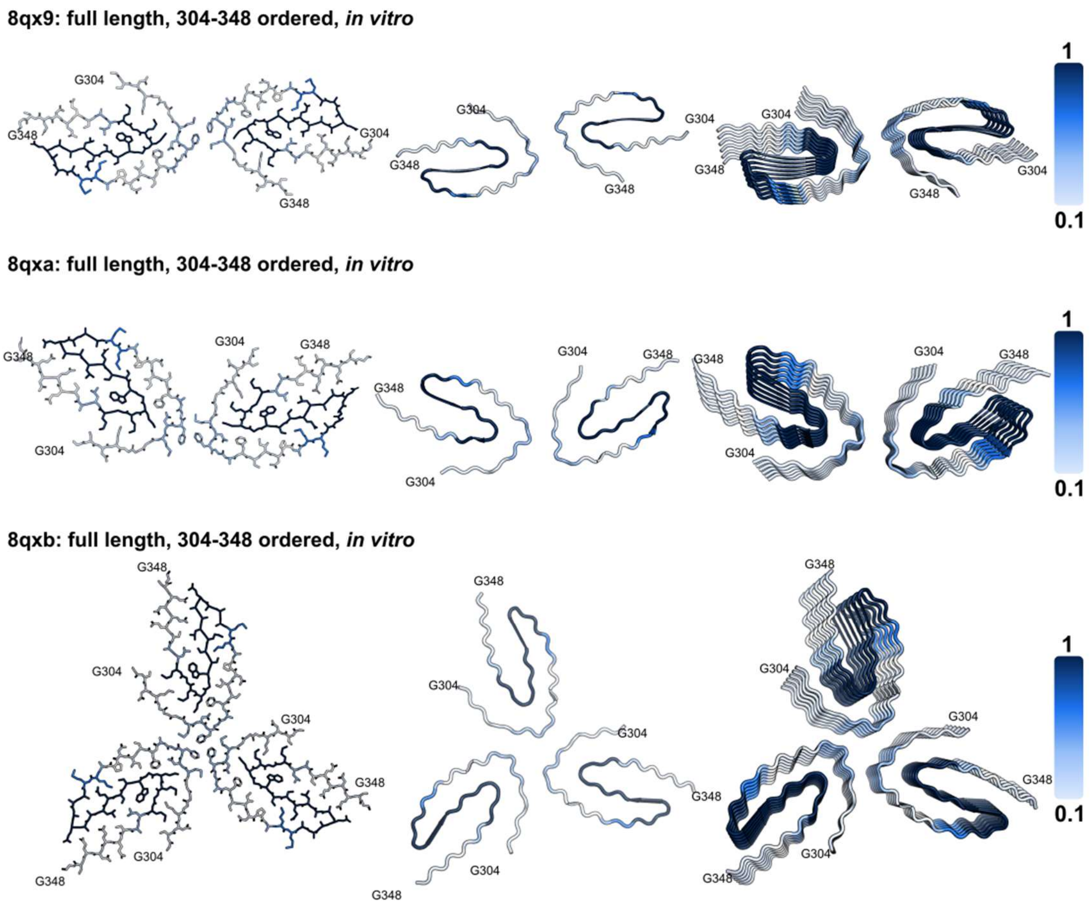

**Figure S6.** (continued)

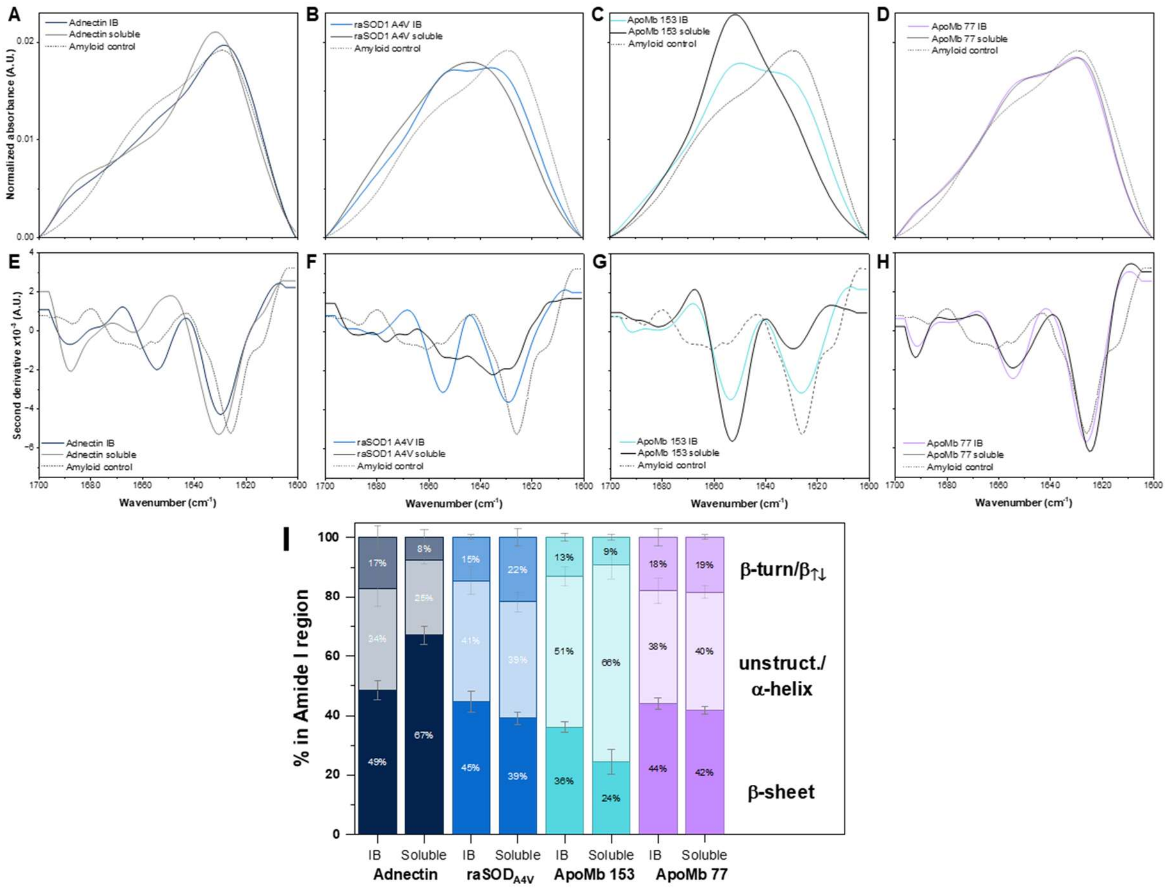

**Figure S7.** Comparison of IB and purified protein by FTIR.[1,2] A)-D) Normalized absorbance spectra of Adnectin, raSOD1 A4V, ApoMb153 and ApoMb77, respectively, for IB and purified soluble protein with E)-H) corresponding second derivative spectra. I) Secondary structure analysis of FTIR spectra. Secondary structure analysis for the proteins used three components: β-turn/antiparallel β-sheet (~1680 cm$^{-1}$), unstructured, alpha-helix (~1654 cm$^{-1}$) and β-sheet (~1630 cm$^{-1}$). Further details are given in Tables S13 and S14 and Figure S8.

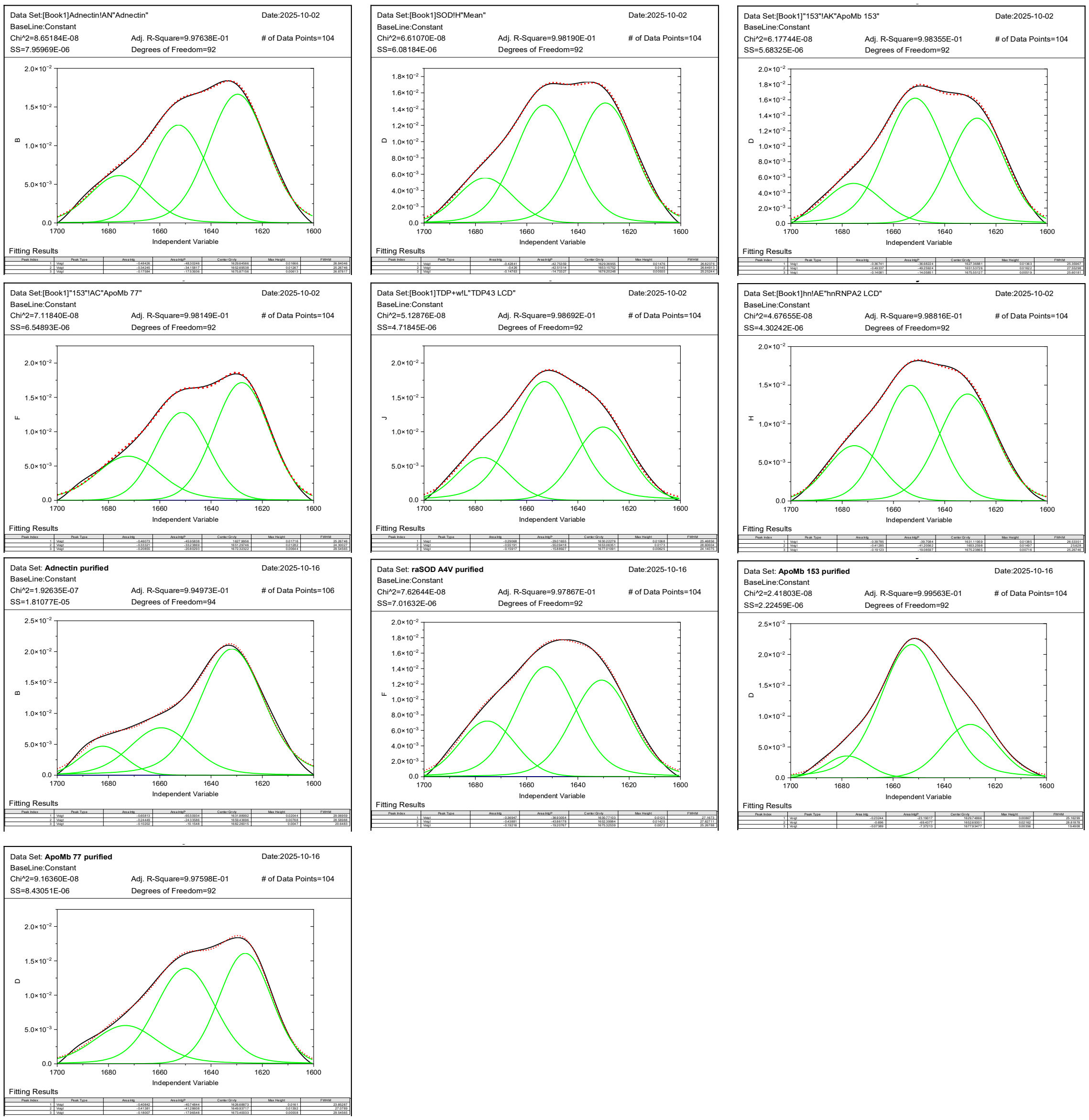

**Figure S8.** FTIR secondary structure analysis of IB and purified proteins. Spectra (black continuous lines) were fit (red dotted lines) using three component peaks (green continuous lines) with Voigt line shapes: β-turn/antiparallel β-sheet (~1680 cm$^{-1}$), unstructured, α-helix (~1654 cm$^{-1}$) and β-sheet (~1630 cm$^{-1}$). For a given spectrum, initial peak position estimates for the fitting were obtained based on the corresponding second derivative spectrum. The Gaussian width was limited to a maximum of 30 cm$^{-1}$ to keep the components within a physically reasonable range. The setting to three component peaks avoids overfitting (green lines). All accepted fits had an $R^2$ greater than 0.99. The fit results, including peak center (cm$^{-1}$), area (%) and Full Width at Half Maximum (FWHM, cm$^{-1}$) are given in Table S14.

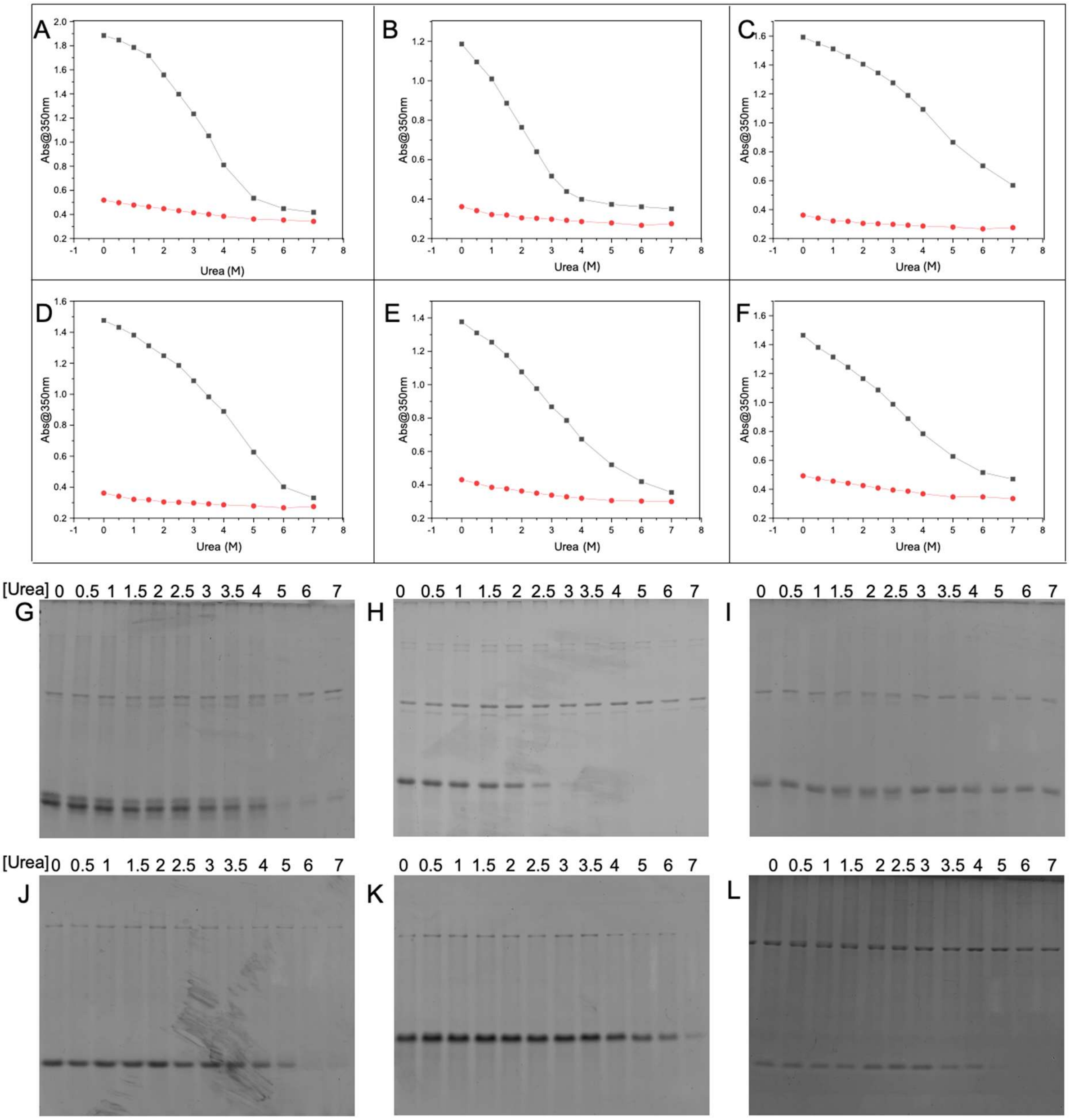

**Figure S9.** Urea solubilization of IBs monitored by SDS-PAGE compared with turbidity. A-F) Solubilization profiles measured after 3 hrs of sample incubation in urea. Panels show: A) Adnectin, B) raSOD1 A4V, C) ApoMb153, D) ApoMb77, E) hnRNPA2$^{LCD}$, and F) TDP-43$^{LCD}$. A-F) Black lines show average IB solubilization profile, red lines show the corresponding background scattering (see Methods 1.9). G-L) Insoluble pellet remaining after IB incubation for 24 hrs in urea was collected by centrifugation and visualized by SDS-PAGE. Only apoMb153 and hnRNPA2$^{LCD}$ gave measurable protein bands at high urea concentration. While relatively low scattering was observed for hnRNPA2$^{LCD}$, the sample nevertheless showed measurable protein, suggesting small species that scattered little were included in the pellet. G) Adnectin, H) raSOD1 A4V, I) ApoMb153, J) ApoMb 77, K) hnRNPA2$^{LCD}$, L) TDP-43$^{LCD}$.

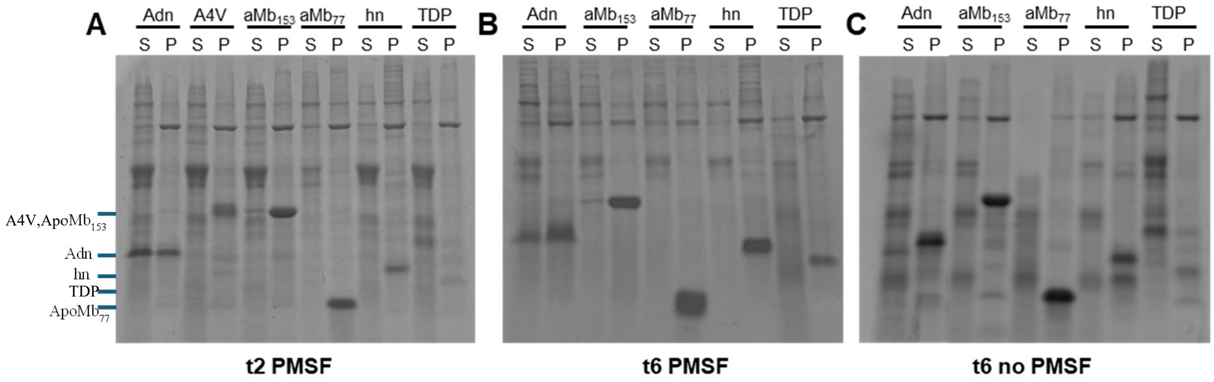

**Figure S10.** IB formation over time and in the presence or absence of PMSF, monitored by SDS-PAGE. A) Protein induction time of 2 hours (t2) with PMSF, B) 6 hours (t6) with PMSF and C) 6 hours (t6) without PMSF. S corresponds to the supernatant and P to the pellet (IB sample). Results are shown for Adnectin (Adn), SOD1 A4V (A4V), ApoMb 153 ($aMb_{153}$), ApoMb 77 ($aMb_{77}$), and LCDs of hnRNPA2 (hn) and TDP-43 (TDP). Note: Soluble expression of Adnectin (Adn) at shorter time (t2), consistent with Adnectin forming IB from the folded soluble state;[26,33] hnRNPA2$^{LCD}$ (hn) has a clear fragment without PMSF; sufficient TDP-43$^{LCD}$ (TDP) IBs could only be obtained with PMSF.

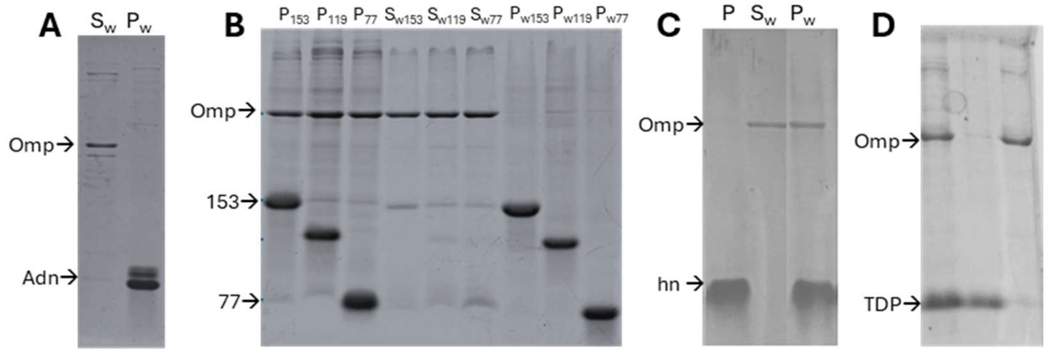

**Figure S11.** IB washing with 1% octyl glucoside, monitored by SDS-PAGE. Results are shown for A) Adnectin (Adn), B) ApoMb153 (153) and ApoMb77 (77), C) hnRNPA2$^{LCD}$ (hn) and D) TDP-43$^{LCD}$ (TDP). Gels contain the pellet IB sample before washing (P), Supernatant after washing ($S_w$) and the pellet IB washed ($P_w$). Further details in Methods 1.1.

### 3. Supplementary Results and Discussion

#### 3.1. SI R&D 1: ATR-FTIR of inclusion bodies, controls for background

To minimize FTIR contributions from species other than the target protein in the IB sample, the following background signal control and subtraction approach was applied. The relative contribution of background proteins was estimated by SDS-PAGE densitometry (Figure S12).

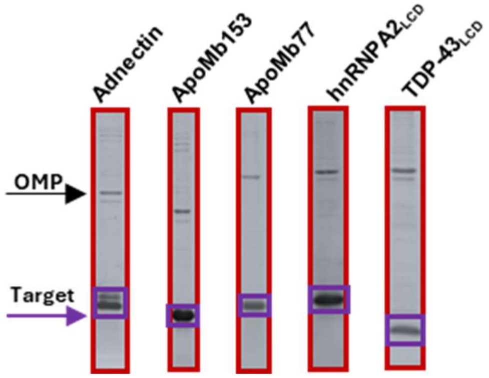

**Figure S12.** SDS-PAGE densitometry analysis for estimating background protein contributions in IB samples. The higher-molecular-weight band corresponds to an outer membrane protein (OMP). In the gel analysis, the red box represents the total protein signal in each lane, whereas the purple region corresponds to the overexpressed target protein.

The fraction of background protein signal, $F_B$, was calculated from gel densitometry (GelAnalyzer 23.11) as:

$$F_B = \frac{(\textbf{All proteins}) - (\textbf{Overexpressed protein})}{(\textbf{All proteins})} \qquad \text{Eq6}$$

Average $F_B$ values were determined from three independent growths.

| Cell background (protein) | Average $F_B$ | Stdev |
|---|---|---|
| BL21 DE3 pLysS tON (Adnectin) | ~0.28 | 0.05 |
| BL21 DE3 pLysS t4 (ApoMb153) | ~0.25 | 0.05 |
| BL21 DE3 pLysS t4 (ApoMb77) | ~0.29 | 0.05 |
| Rosetta DE3 t6 (hnRNPA2$^{LCD}$) | ~0.25 | 0.05 |
| Rosetta DE3 tON (TDP43$^{LCD}$) | ~0.52 | 0.04 |

FTIR spectra were measured for uninduced cell pellet samples (Table M1, Methods 1.1), i.e. tON pLysS for Adnectin, t4 pLysS for ApoMb153 and ApoMb77, t6 Rosetta for hnRNPA2 LCD, and tON Rosetta for TDP-43 LCD (Figure S13).

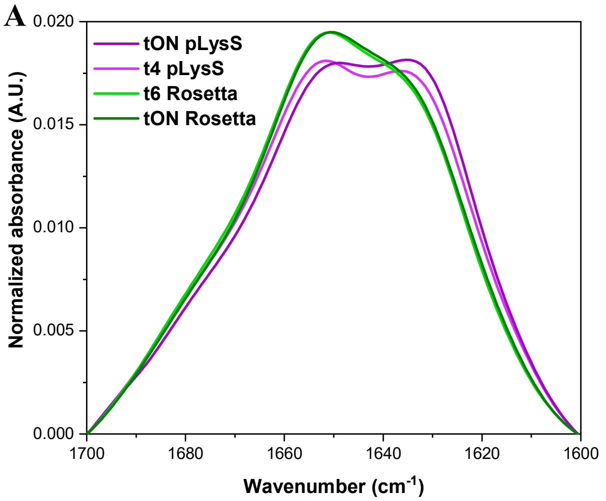

**Figure S13. ATR- FTIR spectra of uninduced cell pellet controls used for background correction.** Spectra were measured for uninduced cells pellet samples which were used as a control for each expression condition background: tON pLysS (Adnectin), t4 pLysS (ApoMb153 and ApoMb77), t6 Rosetta (hnRNPA2 LCD), and tON Rosetta (TDP-43 LCD). **A)** Normalized absorbance amide I spectra. **B)** Corresponding second-derivative spectra.

For these corrections, spectra normalized to the amide I area were used. The background corrected spectrum for IB samples was obtained by subtracting the spectrum of the corresponding uninduced cells, scaled by $F_B$ (Figure S14).

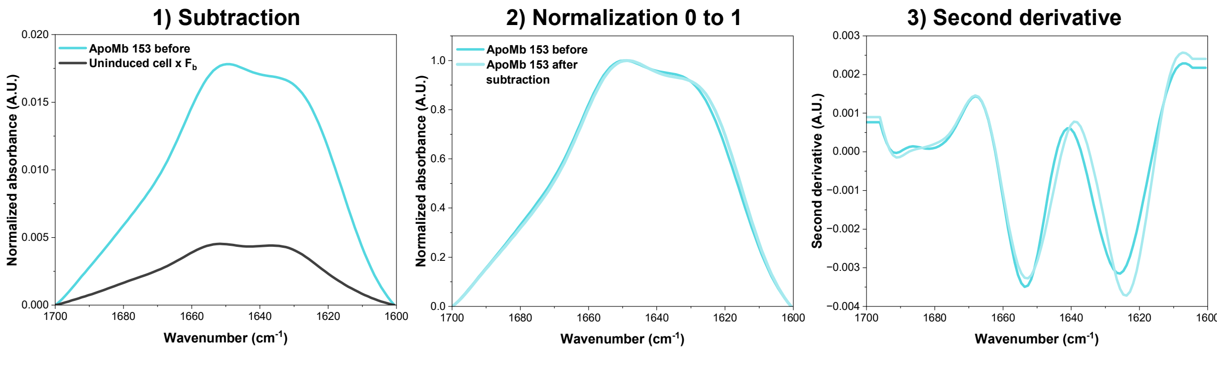

**Figure S14.** Subtraction of the cell background (uninduced cells pellet) from the IB sample. 1) ATR-FTIR spectra of the ApoMb153 IB sample and the corresponding uninduced cells pellet scaled by $\mathbf{F_B}$. This background was subtracted from the ApoMb153 IB spectrum to obtain the background corrected spectrum, which was normalized by area, 2), and is very similar to the uncorrected spectrum. 3) Second derivative spectra corresponding to the spectra in 2).

The background corrected spectra for all proteins are shown in Figure S15, together with spectra obtained for IB samples washed with 1% *n*-octyl glucoside in TEN buffer, which reduced the contribution from outer membrane protein (OMP) (see Methods 1.1, Figure S11). Overall, background correction had relatively little effect on the spectral shape. The largest effect was observed for TDP-43$^{LCD}$, consistent with its higher $F_B$ value (~50%). Importantly, even after background subtraction or detergent washing, the dominant spectral component remained in the unordered/$\alpha$-helical region.

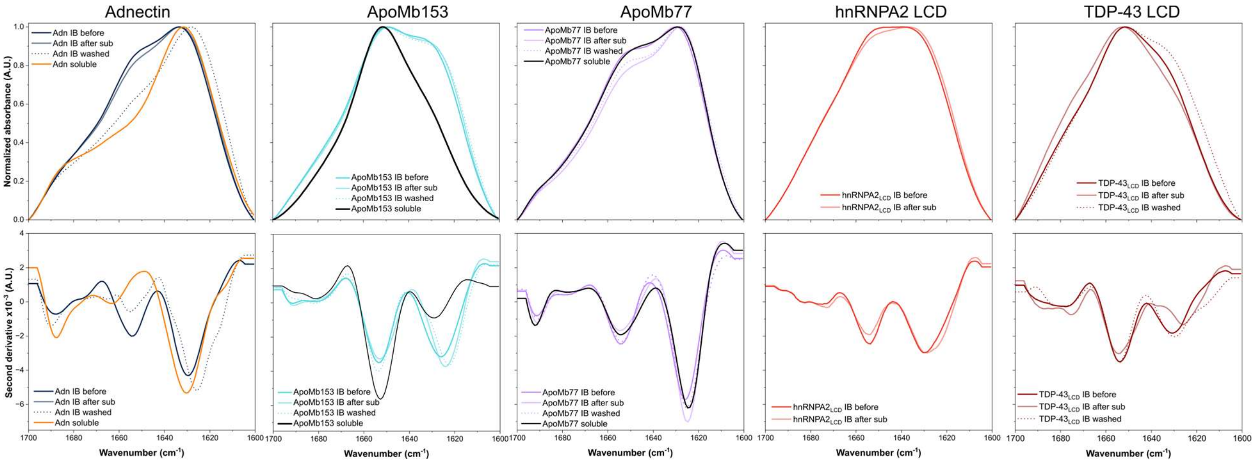

**Figure S15.** Effect of background subtraction and detergent washing on IB FTIR spectra. The top row shows the normalized spectra for IB samples and the corresponding soluble pure protein, and the bottom row shows the corresponding second derivative spectra.

Due to the additional experimental uncertainties associated with background correction, the qualitative analysis in the main text is based on the uncorrected FTIR spectra.

#### 3.2. SI R&D 2: Urea solubilization

The relationship between the equilibrium stability of pure protein measured by urea denaturation compared to apparent IB stability measured by solubilization in urea must be interpreted with caution because the two measurements differ in multiple ways. Fluorescence emission spectra of purified Adnectin in 0, 4, and 8.5 M urea (Figure S16) show that the protein is not fully denatured even in high urea concentrations. In contrast, urea-induced solubilization of Adnectin IBs monitored by turbidity yields an apparent $C_{\text{mid}}$ of ~3 M.

There are multiple possible explanations for this difference between purified Adnectin and Adnectin IBs. One possibility is that the protein molecules in the IBs adopt a less stable conformation, making them more susceptible to unfolding and release from the aggregate upon exposure to urea. It is also possible that fully folded monomers, partially folded species, or small soluble oligomers are released from the aggregate but are not detected, or are underestimated, by the turbidity assay. Accordingly, the apparent IB $C_{\text{mid}}$ measured by turbidity should not be interpreted as a direct measure of unfolding of the protein molecules in the aggregate.

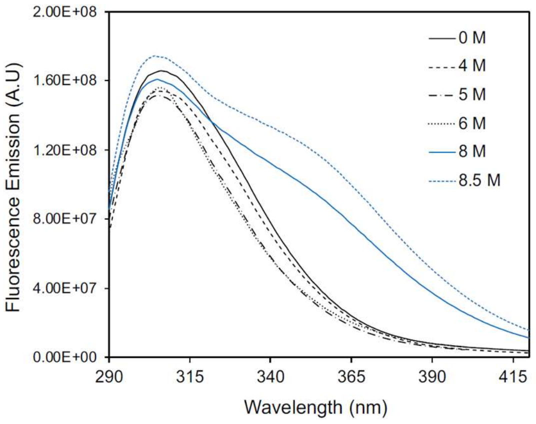

**Figure S16.** Fluorescence emission spectrum of purified Adnectin in 40 mM sodium citrate at pH 4.0. Excitation was at 277 nm with a slit width of 3 and 5 nm, for excitation and emission, respectively, on a Fluorolog 311 (Horiba Scientific).

#### 3.3. SI R&D 3: Lysis results

Because the different cell strains used to make IB samples were lysed using different numbers of freeze–thaw (FT) cycles (see Methods 1.1), control experiments were performed to assess whether FT cycle number influenced the results. IB samples of proteins initially prepared with more FT cycles (Table M1) were also examined after lysis with fewer cycles and gave very similar results (Figure S17).

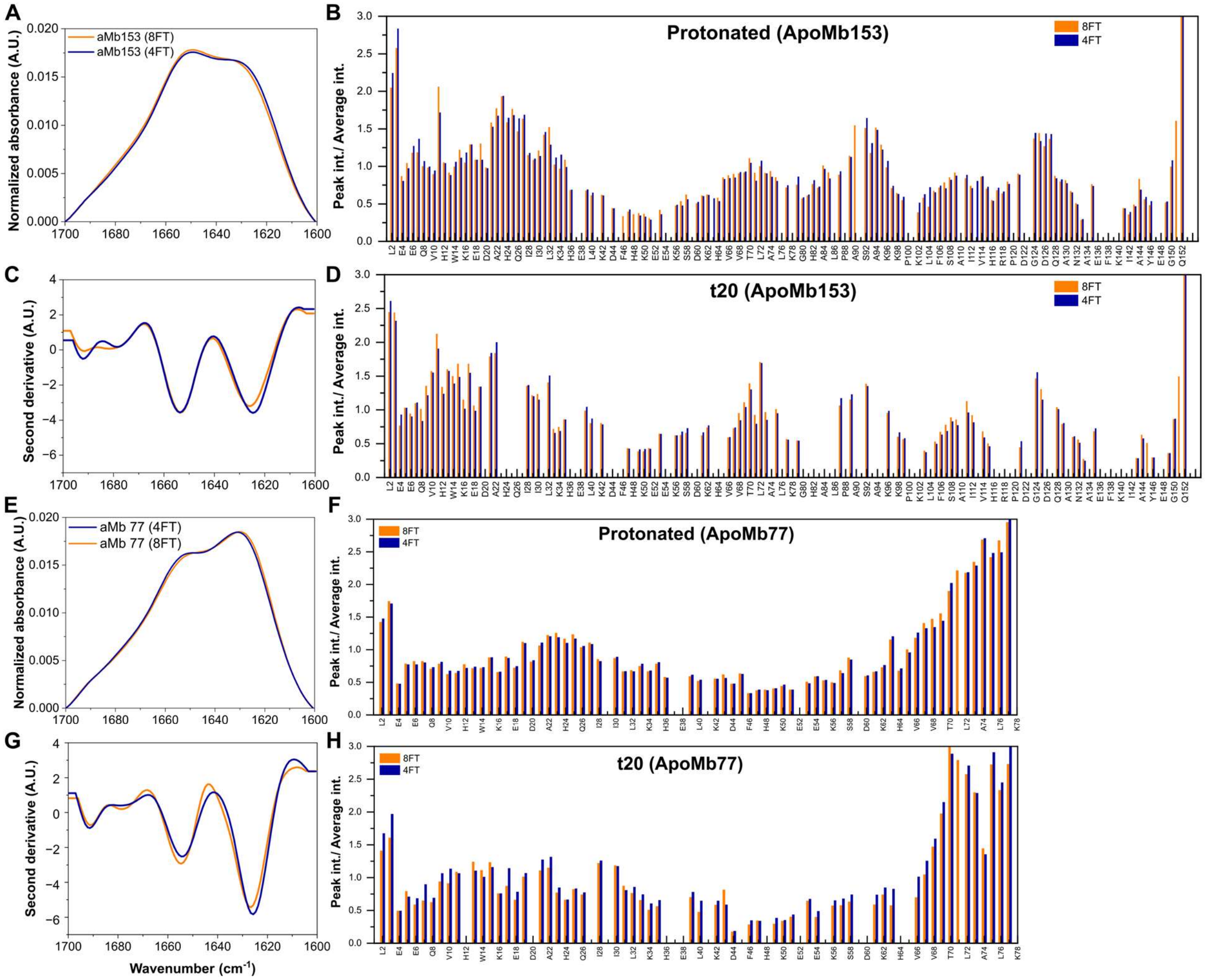

**Figure S17.** Comparison of samples prepared with different numbers of freeze–thaw (FT) cycles during lysis by FTIR and NMR. A, C, B, D) ApoMb 153 FTIR absorbance, FTIR second derivative, NMR protonated sample signal, and NMR exchanged sample signal at t20 (see Methods 1.1, 1.3). Corresponding sets of spectra are shown for ApoMb 77 (E, G, F, H), hnRNPA2$^{LCD}$ (I, K, J, L) and TDP-43$^{LCD}$ (M, O, N, P). The spectra for samples prepared with 4 freeze-thaw cycles (4FT) are very similar to those for samples prepared with 8 freeze-thaw cycles (8FT).

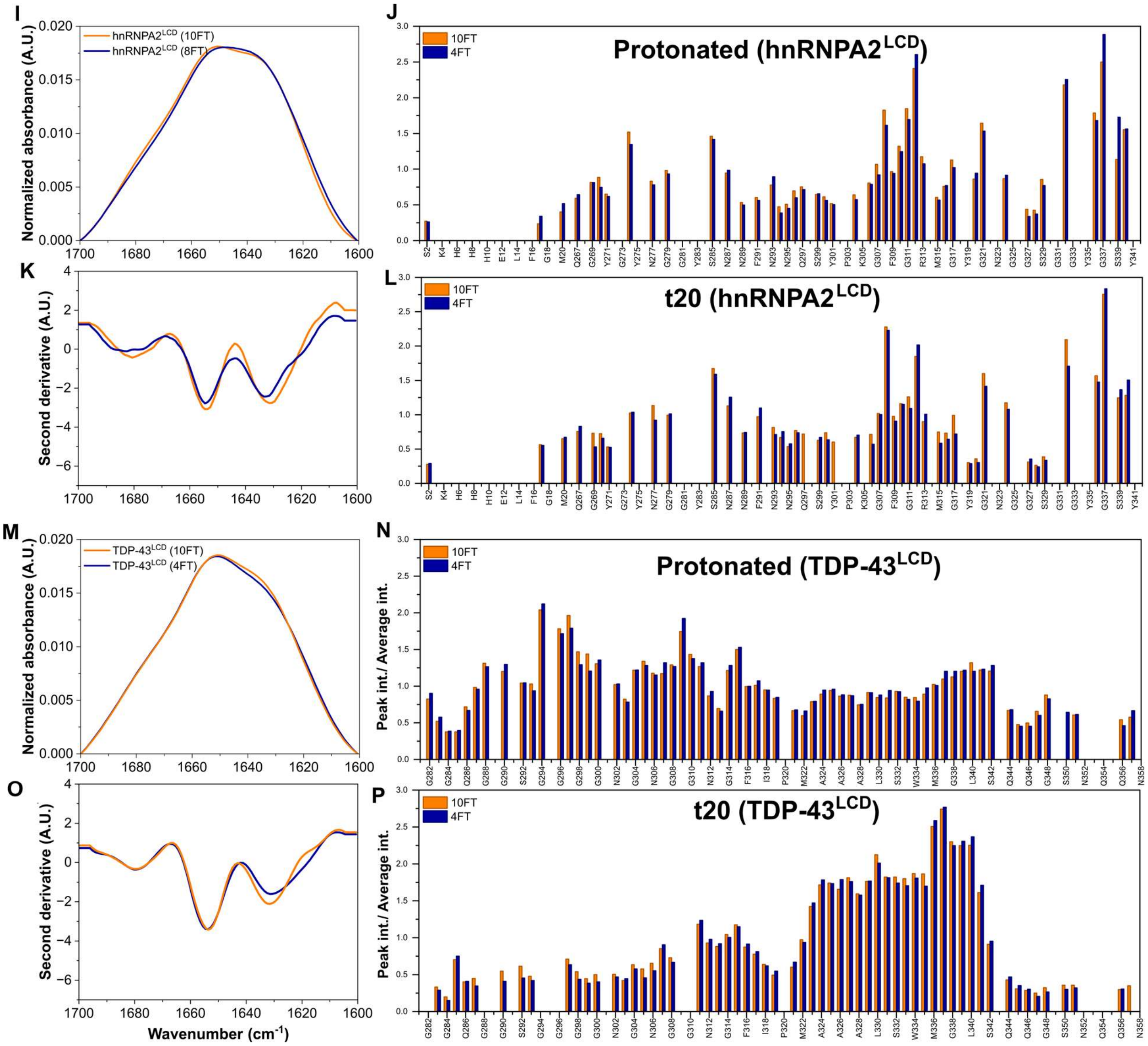

**Figure S17.** continued

### 4. Supplementary Tables

**Table S1.** Summary of biophysical properties of the studied proteins and IB characteristics. Sequence properties include protein length (#aa), affinity tag, hydrophobicity, and predicted isoelectric point (pI). Native-state properties include the predominant structure, and, where available, melting temperature ($\boldsymbol{T_m}$). Qualitative characteristics are shown using plus signs, where a greater number of plus signs indicates a larger relative extent of the indicated property within this protein set. Footnotes define the experimental basis and interpretation of each IB characteristic.

| | **protein** | **Adnectin** | **raSOD1$_{A4V}$** | **ApoMb$^{153}$** | **ApoMb$^{77}$** | **hnRNPA2$^{LCD}$** | **TDP-43$^{LCD}$** |
|---|---|---|---|---|---|---|---|
| **Theoretical** | #aa | 104 | 153 | 153 | 77 | 96 | 86 |
| | Tag | C-terminal His tag | - | - | - | N-terminal TEV tag | C-terminal His tag |
| | Hydrophobicity | ++++ | +++ | +++ | +++++ | - | + |
| | (GRAVY) (Expasy) | -0.311 | -0.393 | -0.367 | -0.157 | -1.38 | -0.71 |
| | pI (Expasy) | 8.22 | 5.54 | 9.01 | 6.56 | 8.19 | 9.79 |
| **Native (exp.)** | Structure | β-sheet | β-sheet | α-helix | Oligomer β-sheet | Disorder | Transient α-helix |
| | $T_m$ (°C) | 70~80[×] | 36.3* | 60[+] | - | - | - |
| **IB (experiments)** | Soluble expression at early time of induction[a] | ++ | - | + | - | - | - |
| | qHDX protection[b] | Extensive | Extensive | Extensive | More localized | Localized | Localized |
| | Proteolysis[c] | + | ++ | ++ | + | ++++ | +++++ |
| | Fraction protection (1D)[d] | 49% | 40% | 48% | 50% | 42% | 35% |
| | FTIR[e] | α/+β | Mix α/β | +α/β | α/+β | Mix α/β | +α/β |
| | Congo red[f] | + | + | ++ | + | ++ | + |
| | Urea ($C_{mid}$)[g] | +++ | + | +++++ | ++++ | ++ | ++ |
| | Urea ($m$)[g] | ++++ | +++++ | + | ++ | ++ | + |

**Native (experiments):** [×]From Trainor *et al.* 2016;[33] *From Vassall *et al.* 2011;[34] [+]From Griko *et al.* 1988[35]

**IB (experiments):**

[a]**Soluble expression at early induction** was analyzed by SDS–PAGE analysis of soluble and pellet fractions at early induction times (Figure S10). Adnectin was observed in both soluble and insoluble fractions at early induction and later mainly in the insoluble fraction, consistent with recruitment of folded soluble species into IBs. ApoMb153 showed limited early soluble expression, whereas SOD1A4V, ApoMb77, hnRNPA2$^{LCD}$, and TDP-43$^{LCD}$ were mainly recovered in the pellet.

[b]**qHDX protection** refers to residue-resolved backbone amide protection against exchange with $D_2O$. Adnectin (Figure 2), raSOD1A4V (Figure S5), and ApoMb153 (Figure 3) show extensive protection consistent with substantial local native-like structure; ApoMb77 (Figure 3) shows a more restricted pattern associated with hydrophobic regions; hnRNPA2$^{LCD}$ and TDP-43$^{LCD}$ show more localized protection, with hnRNPA2$^{LCD}$ (Figure 4) protection toward the N-terminal tag region and TDP-43$^{LCD}$ (Figure 5) protection concentrated in the conserved A321–M339 region.

[c]**Proteolysis** summarizes the susceptibility of IB-associated proteins to cellular proteases during sample preparation, assessed from samples prepared without PMSF by MS and SDS–PAGE (Figure S4, Figure S10). Proteolysis is interpreted together with qHDX as a probe of accessible and flexible regions. Adnectin has cleavages in regions relatively exposed in the native-like monomer and/or domain-swapped structures, supporting a native-like contribution to the IB architecture (Figures 2, S3, S4). Similarly, SOD1$^{A4V}$ shows most cleavages in loop regions (Figure S5). ApoMb153 cleavages occur mainly in regions less stable in the native state, consistent with local native-like structure in the IB (Figure 3). ApoMb77 lacks cleavage in helix E, suggesting that this hydrophobic region contributes to intermolecular aggregation. The LCDs show extensive proteolysis by SDS–PAGE in the absence of PMSF (Figure S10), but cleavage is reduced in the protected regions of hnRNPA2$^{LCD}$ (N-terminal tag) and TDP-43$^{LCD}$ (A324–M337).

[d]**Fraction protection (1D)** was calculated from the amide region of 1D $^{1}$H NMR spectra and provides a global estimate of the fraction of amides protected against exchange for the whole IB sample (Figure 6B). The average fraction of protected amides decreases in the order ApoMb77, Adnectin, ApoMb153, hnRNPA2$^{LCD}$, SOD1$^{A4V}$ and TDP-43$^{LCD}$, reflecting differences in monomer stability, as native SOD1$^{A4V}$ has a substantially lower $T_m$ than Adnectin and ApoMb153. The LCD IBs are toward the lower end of this range, especially for TDP-43LCD, consistent with lower overall protection, greater disorder/accessibility, and reduced stability.

[e]**FTIR** summarizes ATR-FTIR amide I analysis (Figures 6D–F, Figure S7; Table S13, Table S14). The notation describes the unstructured/α-helical signal near ~1654 cm$^{-1}$ (α), and β-sheet near ~1630 cm$^{-1}$ (β). Adnectin and SOD1$^{A4V}$ retain prominent β/antiparallel β features consistent with native-like β-structure; ApoMb153 retains a stronger α/unstructured signal consistent with partial native-like helical content; ApoMb77 is more β-enriched; hnRNPA2$^{LCD}$ shows a mixed α/β signal; and TDP-43$^{LCD}$ has the strongest α/unstructured contribution with a weaker β-sheet signal, consistent with a mostly disordered assembly containing a smaller ordered core.

[f]**Congo red** reports qualitative changes in Congo red absorbance upon binding to IBs (Figure 6C). A larger number of plus signs indicates a larger relative Congo red response within this protein set, not an absolute amyloid content. Overall, all IBs show smaller CR shifts than the amyloid control. hnRNPA2$^{LCD}$ shows the largest CR response, consistent with some amyloid-like/β-structured contribution, whereas Adnectin, SOD1A4V, ApoMb77, and TDP-43$^{LCD}$ show weaker responses; ApoMb153 shows an intermediate response.

[g]**Urea solubilization** was monitored by turbidity (Figures 6G–I, Figure S9). SOD1$^{A4V}$ has the lowest apparent stability, whereas ApoMb153 and ApoMb77 are more resistant to solubilization. Adnectin and SOD1$^{A4V}$ show relatively high *m*-values, consistent with more homogeneous assemblies, whereas ApoMb, hnRNPA2$^{LCD}$, and TDP-43$^{LCD}$ show lower *m*-values, consistent with greater heterogeneity. The LCD IBs show comparatively low stability and high heterogeneity, consistent with loosely packed/disordered assemblies containing smaller locally ordered cores.

**Table S2.** Resonance assignments of Adnectin in 95% DMSO/ 5% $H_2O$ at pH 5.5

| Residue | Assignment (ppm) | | | | |
|---|---|---|---|---|---|
| | $^{1}$H | $^{15}$N | CO | $C_\alpha$ | $C_\beta$ |
| 1 VAL | 8.445 | 116.376 | 171.054 | 58.044 | 31.163 |
| 2 SER | 8.187 | 116.216 | 170.513 | 55.305 | 62.007 |
| 3 ASP | 8.267 | 119.352 | 170.665 | 49.972 | 36.280 |
| 4 VAL | 7.687 | 116.406 | 170.200 | 56.242 | 30.370 |
| 6 ARG | 8.152 | 117.523 | 172.016 | 52.567 | 29.145 |
| 7 ASP | 8.126 | 116.656 | 170.838 | 50.044 | 36.135 |
| 8 LEU | 7.821 | 118.009 | 172.384 | 51.774 | 40.532 |
| 9 GLU | 7.930 | 117.203 | 171.411 | 52.350 | 27.487 |
| 10 VAL | 7.691 | 115.728 | 171.335 | 58.044 | 30.730 |
| 11 VAL | 7.821 | 117.704 | 171.011 | 58.044 | 30.658 |
| 12 ALA | 7.975 | 123.147 | 172.168 | 48.531 | 18.047 |
| 13 ALA | 7.953 | 119.672 | 172.589 | 48.459 | 18.191 |

| | | | | | |
|---|---|---|---|---|---|
| 14 THR | 7.871 | 112.989 | 169.900 | 57.035 | 67.052 |
| 16 THR | 8.014 | 112.379 | 171.378 | 60.494 | 66.620 |
| 17 SER | 7.909 | 115.093 | 171.400 | 57.035 | 61.359 |
| 18 LEU | 7.911 | 120.244 | 173.292 | 52.423 | 40.027 |
| 19 LEU | 7.860 | 118.431 | | 52.423 | 40.171 |
| 21 SER | 7.934 | 115.855 | 171.595 | 56.098 | 61.719 |
| 22 TRP | 8.214 | 120.792 | 173.011 | 56.026 | 27.343 |
| 23 SER | 8.096 | 113.153 | 171.840 | 56.963 | 61.359 |
| 24 ALA | 8.082 | 122.210 | 173.432 | 50.477 | 17.614 |
| 25 ARG | 7.926 | 116.151 | | 54.080 | 28.641 |
| 27 LYS | 7.924 | 118.301 | | 52.423 | 30.730 |
| 29 ALA | 8.036 | 122.759 | 174.092 | 49.612 | 17.614 |
| 30 ARG | 7.939 | 116.041 | | 54.008 | 28.568 |
| 34 ILE | 7.911 | 116.965 | 171.584 | 57.755 | 36.712 |
| 35 THR | 7.709 | 113.858 | 170.405 | 58.260 | 66.908 |
| 36 TYR | 7.895 | 118.575 | | 54.945 | 36.640 |
| 37 GLY | 8.171 | 106.200 | 169.357 | 42.477 | |
| 38 GLU | 8.009 | 117.089 | 171.897 | 52.423 | 27.632 |
| 39 THR | 7.805 | 111.323 | | 58.764 | 67.052 |
| 40 GLY | 8.127 | 107.155 | | 42.621 | |
| 41 GLY | 8.123 | 105.399 | 169.076 | 42.477 | |
| 42 ASN | 8.076 | 116.746 | 171.400 | 50.044 | 37.577 |
| 43 SER | 7.937 | 113.817 | 169.778 | 53.720 | 61.791 |
| 45 VAL | 7.773 | 114.328 | 171.681 | 58.188 | 30.658 |
| 46 GLN | 8.031 | 120.358 | 171.508 | 52.495 | 27.848 |
| 47 GLU | 7.871 | 117.466 | 171.346 | 52.495 | 27.920 |
| 48 PHE | 7.973 | 116.912 | 171.573 | 54.152 | 37.577 |
| 49 THR | 8.036 | 112.049 | 170.243 | 58.404 | 67.124 |
| 50 VAL | 7.753 | 118.417 | | 56.026 | 30.514 |
| 52 LYS | 8.088 | 117.412 | 172.005 | 52.495 | 31.595 |
| 53 ASN | 8.148 | 117.34 | 171.616 | 50.188 | 37.000 |
| 54 VAL | 7.638 | 114.001 | 171.281 | 58.188 | 30.586 |
| 55 TYR | 8.041 | 118.826 | 172.038 | 54.873 | 36.496 |
| 56 THR | 7.808 | 111.231 | | 58.620 | 67.052 |
| 57 ALA | 7.916 | 122.256 | 172.935 | 48.747 | 18.407 |
| 58 THR | 7.839 | 110.147 | 170.600 | 58.476 | 67.052 |
| 59 ILE | 7.734 | 116.896 | 171.562 | 57.755 | 36.784 |
| 60 SER | 7.985 | 115.953 | 170.978 | 55.954 | 61.863 |
| 61 GLY | 8.049 | 107.476 | 170.211 | 42.621 | |
| 62 LEU | 7.775 | 117.666 | 172.395 | 51.125 | 40.892 |
| 63 LYS | 8.081 | 119.375 | 170.492 | 51.702 | 30.442 |
| 65 GLY | 8.204 | 105.593 | 169.422 | 42.477 | |
| 66 VAL | 7.621 | 113.255 | 171.335 | 57.900 | 30.947 |
| 67 ASP | 8.256 | 119.314 | 170.719 | 50.044 | 36.135 |

| | | | | | |
|---|---|---|---|---|---|
| 68 TYR | 7.654 | 115.831 | 171.216 | 54.440 | 36.712 |
| 69 THR | 7.924 | 111.885 | 169.140 | 58.404 | 67.052 |
| 70 ILE | 7.790 | 117.673 | 171.540 | 57.683 | 37.144 |
| 71 THR | 7.906 | 115.034 | 170.373 | 58.692 | 67.052 |
| 72 VAL | 7.615 | 116.224 | 171.173 | 58.044 | 30.586 |
| 73 TYR | 7.896 | 118.929 | 171.303 | 54.440 | 36.784 |
| 74 ALA | 7.993 | 120.701 | 172.773 | 48.747 | 18.335 |
| 75 VAL | 7.887 | 114.114 | 171.422 | 58.188 | 30.586 |
| 76 THR | 7.677 | 112.560 | 170.243 | 58.044 | 67.124 |
| 77 LEU | 8.104 | 118.514 | 171.086 | 50.116 | 36.352 |
| 78 LEU | 7.817 | 116.799 | | 55.017 | 36.640 |
| 82 CYS | 8.325 | 119.695 | 168.816 | 53.792 | 25.758 |
| 84 ILE | 7.885 | 115.723 | 171.573 | 57.611 | 37.144 |
| 85 SER | 7.958 | 116.344 | 170.622 | 55.377 | 61.719 |
| 86 ILE | 7.790 | 117.722 | 171.216 | 57.611 | 37.072 |
| 87 ASN | 8.068 | 119.607 | | 50.549 | 37.289 |
| 89 ARG | 8.120 | 117.838 | 172.005 | 52.783 | 29.001 |
| 90 THR | 7.764 | 111.445 | 170.449 | 58.692 | 66.908 |
| 91 GLU | 7.908 | 118.908 | 171.595 | 52.350 | 27.560 |
| 92 ILE | 7.810 | 116.665 | 171.465 | 57.467 | 36.712 |
| 93 ASP | 8.239 | 120.610 | 170.740 | 49.900 | 36.352 |
| 94 LYS | 7.884 | 118.453 | | 50.765 | 30.658 |
| 96 SER | 8.203 | 112.867 | 171.032 | 56.098 | 61.575 |

**Table S3.** Resonance assignments of ApoMb153 in 95% DMSO/ 5% $H_2O$ at pH 5.5

| Residues | Assignment (ppm) | | | | |
|---|---|---|---|---|---|
| | $^{1}H$ | $^{15}N$ | CO | $C_\alpha$ | $C_\beta$ |
| 0 MET | | | | | |
| 1 VAL | 8.517 | 120.093 | 170.996 | 58.488 | 30.836 |
| 2 LEU | 8.210 | 122.480 | 172.675 | 51.089 | 40.805 |
| 3 SER | 7.973 | 113.274 | 170.997 | 55.227 | 61.811 |
| 4 GLU | 8.288 | 119.485 | 172.586 | 52.780 | 27.066 |
| 5 GLY | 8.294 | 106.037 | 169.971 | 42.567 | |
| 6 GLU | 7.990 | 117.772 | 172.557 | 52.904 | 27.235 |
| 7 TRP | 8.161 | 119.457 | 172.732 | 54.735 | 27.359 |
| 8 GLN | 8.237 | 117.855 | 172.46 | 53.686 | 27.521 |
| 9 LEU | 7.939 | 119.075 | 173.135 | 52.44 | 40.429 |
| 10 VAL | 7.814 | 116.258 | 172.169 | 59.241 | 30.509 |
| 11 LEU | 8.072 | 120.993 | 173.24 | 52.459 | 40.489 |
| 12 HIS | 8.212 | 116.972 | 170.978 | 52.408 | 26.926 |
| 13 VAL | 7.862 | 115.943 | 172.000 | 58.550 | 30.648 |
| 14 TRP | 8.205 | 120.878 | 172.32 | 54.330 | 27.360 |

| | | | | | |
|---|---|---|---|---|---|
| 15 ALA | 8.116 | 120.891 | 173.391 | 49.153 | 17.998 |
| 16 LYS | 7.989 | 117.065 | 172.507 | 53.965 | 31.054 |
| 17 VAL | 7.802 | 115.313 | 172.035 | 58.641 | 30.927 |
| 18 GLU | 8.073 | 119.461 | 171.700 | 52.654 | 27.278 |
| 19 ALA | 7.968 | 120.238 | 173.179 | 48.792 | 18.124 |
| 20 ASP | 8.269 | 116.527 | 171.135 | 50.171 | 35.921 |
| 21 VAL | 7.747 | 113.845 | 171.318 | 58.078 | 30.956 |
| 22 ALA | 8.098 | 122.927 | 173.35 | 48.984 | 18.091 |
| 23 GLY | 8.130 | 104.618 | 169.679 | 42.540 | |
| 24 HIS | 8.159 | 115.078 | 170.912 | 52.194 | 27.278 |
| 25 GLY | 8.415 | 107.116 | 169.517 | 42.500 | |
| 26 GLN | 8.188 | 117.562 | 171.986 | 52.515 | 28.204 |
| 27 ASP | 8.406 | 118.673 | 171.023 | 50.102 | 36.181 |
| 28 ILE | 7.720 | 116.135 | 171.447 | 57.620 | 36.645 |
| 29 LEU | 8.063 | 121.899 | 172.501 | 51.728 | 40.600 |
| 30 ILE | 7.766 | 116.951 | 171.582 | 57.62 | 36.738 |
| 31 ARG | 8.017 | 121.035 | 171.641 | 52.44 | 28.983 |
| 32 LEU | 7.926 | 119.267 | 172.578 | 51.454 | 40.914 |
| 33 PHE | 7.971 | 116.677 | 171.593 | 54.182 | 37.380 |
| 34 LYS | 7.932 | 118.075 | 171.981 | 52.515 | 31.541 |
| 35 SER | 8.035 | 114.279 | 170.253 | 55.578 | 61.889 |
| 36 HIS | 8.281 | 117.430 | 168.861 | 50.263 | 27.033 |
| 37 PRO | | | 172.823 | 60.218 | 29.685 |
| 38 GLU | 8.515 | 118.158 | 171.916 | 52.979 | 27.364 |
| 39 THR | 7.748 | 112.470 | 170.531 | 58.918 | 66.92 |
| 40 LEU | 8.032 | 120.792 | 172.961 | 52.367 | 40.513 |
| 41 GLU | 8.043 | 117.846 | 171.868 | 52.718 | 27.235 |
| 42 LYS | 7.877 | 118.267 | 172.103 | 52.718 | 31.328 |
| 43 PHE | 8.038 | 117.080 | 171.786 | 54.061 | 37.139 |
| 44 ASP | 8.422 | 117.657 | 171.465 | 50.213 | 36.088 |
| 45 ARG | 8.057 | 118.056 | 172.575 | 52.730 | 29.153 |
| 46 PHE | 8.013 | 116.584 | 172.000 | 54.412 | 37.357 |
| 47 LYS | 8.220 | 118.274 | 172.394 | 53.174 | 31.371 |
| 48 HIS | 8.303 | 116.770 | 170.711 | 52.857 | 26.808 |
| 49 LEU | 8.177 | 119.752 | 172.861 | 51.970 | 40.815 |
| 50 LYS | 8.271 | 119.309 | 172.760 | 52.170 | 31.018 |
| 51 THR | 7.818 | 111.779 | 170.724 | 58.415 | 67.042 |
| 52 GLU | 8.159 | 119.437 | 172.516 | 53.205 | 27.187 |
| 53 ALA | 8.204 | 121.043 | 173.814 | 49.751 | 17.724 |
| 54 GLU | 7.993 | 116.232 | 172.863 | 52.604 | 28.139 |
| 55 MET | 8.042 | 117.75 | 172.544 | 52.837 | 29.301 |
| 56 LYS | 8.075 | 118.339 | 172.591 | 52.745 | 31.200 |
| 57 ALA | 8.060 | 120.787 | 173.748 | 49.075 | 17.999 |

| | | | | | |
|---|---|---|---|---|---|
| 58 SER | 7.989 | 112.325 | 171.353 | 56.592 | 61.451 |
| 59 GLU | 7.971 | 117.982 | 172.225 | 52.470 | 27.258 |
| 60 ASP | 8.203 | 117.275 | 171.578 | 50.447 | 35.967 |
| 61 LEU | 7.855 | 118.557 | 172.968 | 52.010 | 40.426 |
| 62 LYS | 7.849 | 117.540 | 172.501 | 53.133 | 30.864 |
| 63 LYS | 7.965 | 118.175 | 172.223 | 53.146 | 30.993 |
| 64 HIS | 8.125 | 115.308 | 170.689 | 52.010 | 27.370 |
| 65 GLY | 8.302 | 106.599 | 169.391 | 42.356 | |
| 66 VAL | 8.058 | 115.537 | 171.592 | 57.986 | 30.864 |
| 67 THR | 8.087 | 115.852 | 170.542 | 58.975 | 66.812 |
| 68 VAL | 7.828 | 117.411 | 171.534 | 58.234 | 30.825 |
| 69 LEU | 8.147 | 121.590 | 173.043 | 51.725 | 40.450 |
| 70 THR | 7.799 | 111.152 | 170.929 | 58.505 | 67.008 |
| 71 ALA | 8.212 | 122.575 | 173.748 | 49.800 | 17.349 |
| 72 LEU | 8.051 | 116.91 | 173.607 | 52.459 | 40.267 |
| 73 GLY | 8.128 | 105.931 | 169.921 | 42.375 | |
| 74 ALA | 7.969 | 120.558 | 173.597 | 49.434 | 17.899 |
| 75 ILE | 7.901 | 115.921 | 172.320 | 58.63 | 36.105 |
| 76 LEU | 8.027 | 121.09 | 173.231 | 52.378 | 40.334 |
| 77 LYS | 7.932 | 118.066 | 172.55 | 53.475 | 31.048 |
| 78 LYS | | | 172.522 | 53.200 | 31.108 |
| 79 LYS | 8.011 | 118.036 | 172.760 | 53.100 | 31.100 |
| 80 GLY | 8.212 | 106.208 | 169.573 | 42.438 | |
| 81 HIS | 8.211 | 115.652 | 170.605 | 52.128 | 27.246 |
| 82 HIS | 8.349 | 116.724 | 170.574 | 52.238 | 27.458 |
| 83 GLU | 8.349 | 118.716 | 171.625 | 52.528 | 27.246 |
| 84 ALA | 8.272 | 122.188 | 172.739 | 48.690 | 18.118 |
| 85 GLU | 8.101 | 116.655 | 171.493 | 52.128 | 27.433 |
| 86 LEU | 7.909 | 119.315 | 172.369 | 51.183 | 40.793 |
| 87 LYS | 8.127 | 119.145 | 170.472 | 52.642 | 30.641 |
| 88 PRO | | | 172.335 | 59.733 | 29.393 |
| 89 LEU | 8.050 | 117.701 | 172.628 | 52.654 | 40.562 |
| 90 ALA | 7.963 | 120.148 | 172.865 | 48.769 | 17.982 |
| 91 GLN | 7.995 | 116.270 | 171.951 | 52.503 | 28.309 |
| 92 SER | 8.014 | 113.902 | 170.703 | 55.541 | 61.748 |
| 93 HIS | 8.288 | 117.225 | 170.162 | 52.029 | 27.262 |
| 94 ALA | 8.239 | 121.603 | 173.248 | 49.000 | 18.000 |
| 95 THR | 7.989 | 111.178 | 170.479 | 58.568 | 66.945 |
| 96 LYS | 8.026 | 119.525 | 172.108 | 52.656 | 31.275 |
| 97 HIS | 8.273 | 116.220 | 170.242 | 52.066 | 27.308 |
| 98 LYS | 8.231 | 118.624 | 172.126 | 52.629 | 31.435 |
| 99 ILE | 8.182 | 119.440 | 170.507 | 55.117 | 36.417 |
| 101 ILE | | | 171.958 | 57.959 | 36.417 |

| | | | | | |
|---|---|---|---|---|---|
| 102 LYS | 8.097 | 120.632 | 172.112 | 53.113 | 31.324 |
| 103 TYR | 7.818 | 116.839 | 171.781 | 54.557 | 36.645 |
| 104 LEU | 8.028 | 118.664 | 172.684 | 51.826 | 40.610 |
| 105 GLU | 7.951 | 116.807 | 171.693 | 52.650 | 27.716 |
| 106 PHE | 7.956 | 117.350 | 171.526 | 54.209 | 37.250 |
| 107 ILE | 8.044 | 117.145 | 171.791 | 57.802 | 36.564 |
| 108 SER | 8.063 | 116.655 | 170.954 | 55.950 | 61.000 |
| 109 GLU | 8.028 | 118.975 | 171.587 | 52.515 | 27.450 |
| 110 ALA | 8.062 | 121.124 | 172.975 | 48.884 | 17.809 |
| 111 ILE | 7.802 | 115.801 | 171.850 | 57.793 | 36.411 |
| 112 ILE | 7.832 | 119.313 | 171.826 | 57.793 | 36.411 |
| 113 HIS | 8.269 | 119.462 | 170.715 | 52.330 | 27.179 |
| 114 VAL | 7.887 | 116.347 | 171.765 | 58.641 | 30.588 |
| 115 LEU | 8.161 | 121.661 | 172.675 | 51.916 | 40.475 |
| 116 HIS | 8.197 | 116.053 | 170.452 | 51.889 | 27.340 |
| 117 SER | 8.078 | 114.046 | 170.684 | 55.442 | 61.881 |
| 118 ARG | 8.285 | 119.839 | 171.735 | 52.791 | 28.934 |
| 119 HIS | 8.311 | 116.856 | 169.074 | 50.171 | 26.635 |
| 120 PRO | | | 172.837 | 60.381 | 29.747 |
| 121 GLY | 8.449 | 105.782 | 169.243 | 42.188 | |
| 122 ASN | 8.036 | 116.614 | 171.531 | 49.878 | 37.231 |
| 123 PHE | 8.200 | 117.638 | 172.141 | 52.470 | 38.106 |
| 124 GLY | 8.335 | 106.264 | 169.615 | 42.688 | |
| 125 ALA | 8.116 | 120.330 | 173.392 | 49.127 | 18.056 |
| 126 ASP | 8.288 | 115.763 | 171.386 | 49.878 | 35.873 |
| 127 ALA | 7.906 | 120.591 | 173.549 | 49.628 | 17.680 |
| 128 GLN | 8.154 | 116.929 | 173.018 | 53.629 | 27.371 |
| 129 GLY | 8.210 | 106.606 | 170.371 | 42.751 | |
| 130 ALA | 8.109 | 121.328 | 174.304 | 49.878 | 17.68 |
| 131 MET | 8.158 | 116.354 | 172.963 | 53.565 | 31.024 |
| 132 ASN | 8.233 | 118.093 | 173.341 | 51.459 | 36.792 |
| 133 LYS | 8.270 | 120.079 | 173.970 | 53.849 | 29.000 |
| 134 ALA | 8.108 | 120.593 | 173.388 | 49.16 | 17.999 |
| 135 LEU | 7.912 | 116.871 | 172.003 | 52.192 | 40.748 |
| 140 LYS | | | 173.841 | 52.470 | 30.278 |
| 141 ASP | 8.341 | 118.184 | 172.919 | 51.826 | 35.461 |
| 142 ILE | 7.769 | 117.167 | 171.574 | 57.434 | 36.656 |
| 143 ALA | 8.067 | 122.141 | 174.694 | 50.999 | 16.797 |
| 144 ALA | 8.052 | 118.442 | 174.438 | 50.447 | 16.889 |
| 145 LYS | 7.775 | 116.120 | 173.388 | 52.768 | 30.613 |
| 146 TYR | 7.790 | 115.482 | 172.999 | 56.675 | 36.233 |
| 148 GLU | | | 172.014 | 53.304 | 26.761 |
| 149 LEU | 7.662 | 117.085 | 172.863 | 51.819 | 40.682 |

| | | | | | |
|---|---|---|---|---|---|
| 150 GLY | 7.965 | 104.773 | 169.252 | 42.353 | |
| 151 TYR | 7.889 | 116.403 | 171.541 | 54.975 | 36.831 |
| 152 GLN | 8.194 | 117.644 | 172.122 | 52.608 | 28.200 |
| 153 GLY | 8.017 | 105.654 | 171.534 | 41.008 | |

**Table S4**. Resonance assignments of ApoMb77 in 95% DMSO/ 5% $H_2O$ at pH 5.5

| Residue | **Assignment (ppm)** | | | | |
|---|---|---|---|---|---|
| | $^{1}H$ | $^{15}N$ | CO | $C_\alpha$ | $C_\beta$ |
| 0 MET | | | | | |
| 1 VAL | 8.507 | 120.128 | 171.013 | 58.54 | 30.852 |
| 2 LEU | 8.203 | 122.487 | 172.742 | 51.086 | 40.812 |
| 3 SER | 7.974 | 113.325 | 171.088 | 55.22 | 61.680 |
| 4 GLU | 8.328 | 119.695 | 172.761 | 53.278 | 27.093 |
| 5 GLY | 8.319 | 106.038 | 170.139 | 42.691 | |
| 6 GLU | 7.986 | 117.935 | 172.752 | 53.033 | 27.136 |
| 7 TRP | 8.177 | 119.507 | 172.865 | 54.914 | 27.449 |
| 8 GLN | 8.233 | 117.752 | 172.639 | 53.099 | 27.470 |
| 9 LEU | 7.908 | 119.099 | 173.306 | 52.652 | 40.680 |
| 10 VAL | 7.825 | 116.540 | 172.648 | 59.479 | 30.961 |
| 11 LEU | 8.063 | 120.820 | 173.381 | 52.367 | 40.374 |
| 12 HIS | 8.208 | 116.861 | 171.079 | 52.514 | 26.822 |
| 13 VAL | 7.860 | 116.002 | 172.207 | 57.987 | 30.750 |
| 14 TRP | 8.206 | 120.847 | 172.847 | 54.594 | 27.324 |
| 15 ALA | 8.119 | 120.863 | 173.428 | 49.208 | 17.918 |
| 16 LYS | 7.969 | 117.019 | 172.63 | 53.742 | 31.023 |
| 17 VAL | 7.799 | 115.433 | 172.178 | 58.800 | 30.476 |
| 18 GLU | 8.083 | 119.431 | 171.84 | 52.719 | 27.261 |
| 19 ALA | 7.962 | 120.218 | 173.278 | 48.832 | 18.106 |
| 20 ASP | 8.246 | 116.455 | 171.229 | 50.148 | 35.852 |
| 21 VAL | 7.562 | 114.016 | 171.427 | 58.112 | 30.710 |
| 22 ALA | 8.094 | 122.911 | 173.457 | 49.020 | 17.900 |
| 23 GLY | 8.129 | 104.622 | 169.735 | 42.498 | |
| 24 HIS | 8.154 | 115.062 | 170.938 | 52.218 | 27.136 |
| 25 GLY | 8.406 | 107.116 | 169.575 | 42.498 | |
| 26 GLN | 8.183 | 117.562 | 172.019 | 52.594 | 28.39 |
| 27 ASP | 8.401 | 118.663 | 171.098 | 50.023 | 35.977 |
| 28 ILE | 7.730 | 116.263 | 171.530 | 57.673 | 36.729 |
| 29 LEU | 8.056 | 121.944 | 172.601 | 51.653 | 40.555 |
| 30 ILE | 7.767 | 117.017 | 171.652 | 57.610 | 36.729 |
| 31 ARG | 8.014 | 121.03 | 171.718 | 52.594 | 28.954 |
| 32 LEU | 7.910 | 119.202 | 172.639 | 51.649 | 40.680 |
| 33 PHE | 7.973 | 116.732 | 171.633 | 54.218 | 37.429 |
| 34 LYS | 8.116 | 118.112 | 172.019 | 52.406 | 30.961 |

| | | | | | |
|---|---|---|---|---|---|
| 35 SER | 8.016 | 114.237 | 170.271 | 55.721 | 61.798 |
| 36 HIS | 8.285 | 117.356 | 168.805 | 50.209 | 26.592 |
| 37 PRO | | | 172.874 | 60.044 | 29.599 |
| 38 GLU | 8.486 | 118.188 | 171.991 | 52.777 | 27.281 |
| 39 THR | 7.758 | 112.569 | 170.600 | 58.853 | 66.891 |
| 40 LEU | 8.035 | 120.856 | 173.090 | 52.367 | 40.499 |
| 41 GLU | 8.051 | 117.940 | 171.791 | 52.589 | 26.987 |
| 42 LYS | 7.883 | 118.300 | 172.188 | 52.589 | 31.323 |
| 43 PHE | 8.023 | 117.079 | 171.776 | 54.225 | 37.370 |
| 44 ASP | 8.418 | 117.712 | 171.530 | 50.271 | 36.176 |
| 46 PHE | 7.990 | 116.554 | 171.991 | 54.412 | 37.357 |
| 47 LYS | 8.211 | 118.264 | 172.517 | 53.247 | 31.149 |
| 48 HIS | 8.295 | 116.752 | 170.788 | 52.280 | 27.261 |
| 49 LEU | 8.173 | 119.773 | 173.278 | 51.817 | 40.812 |
| 50 LYS | 8.267 | 119.235 | 172.808 | 52.965 | 31.165 |
| 51 THR | 7.802 | 111.765 | 170.445 | 58.352 | 66.872 |
| 52 GLU | 8.188 | 119.612 | 172.667 | 53.206 | 27.033 |
| 53 ALA | 8.213 | 121.033 | 173.983 | 49.804 | 17.604 |
| 54 GLU | 7.978 | 116.286 | 172.968 | 52.917 | 26.947 |
| 56 LYS | 8.076 | 118.376 | 172.752 | 51.920 | 30.726 |
| 57 ALA | 8.056 | 120.721 | 173.861 | 49.25 | 17.822 |
| 58 SER | 7.982 | 112.296 | 171.464 | 56.473 | 61.547 |
| 59 GLU | 7.967 | 118.028 | 172.216 | 53.101 | 27.094 |
| 60 ASP | 8.195 | 117.205 | 171.652 | 50.459 | 35.926 |
| 61 LEU | 7.841 | 118.582 | 172.043 | 52.151 | 40.436 |
| 62 LYS | 7.849 | 117.592 | 172.583 | 53.216 | 31.040 |
| 63 LYS | 7.958 | 117.949 | 172.225 | 53.158 | 30.914 |
| 64 HIS | 8.111 | 115.192 | 170.637 | 52.088 | 27.281 |
| 65 GLY | 8.288 | 106.597 | 169.369 | 42.316 | |
| 66 VAL | 8.010 | 115.457 | 171.539 | 57.976 | 30.961 |
| 67 THR | 8.078 | 116.044 | 170.431 | 58.900 | 66.872 |
| 68 VAL | 7.807 | 117.385 | 171.436 | 57.976 | 30.977 |
| 69 LEU | 8.174 | 121.977 | 172.733 | 51.649 | 40.561 |
| 70 THR | 7.715 | 111.014 | 170.534 | 58.164 | 67.122 |
| 71 ALA | 8.100 | 122.14 | 173.005 | 49.144 | 17.759 |
| 72 LEU | 7.907 | 116.692 | 173.000 | 51.900 | 40.561 |
| 73 GLY | 8.093 | 105.527 | 169.143 | 42.316 | |
| 74 ALA | 7.905 | 120.236 | 172.695 | 48.643 | 18.26 |
| 75 ILE | 7.989 | 115.486 | 171.352 | 57.475 | 36.74 |
| 76 LEU | 7.954 | 121.803 | 172.423 | 51.211 | 40.937 |
| 77 LYS | 8.075 | 118.109 | 173.832 | 51.900 | 30.726 |

**Table S5**. Resonance assignments of hnRNPA2$^{LCD}$ in 95% DMSO/ 5% $H_2O$ at pH 5.5

| Residue | Assignment (ppm) | | | | |
|---|---|---|---|---|---|
| | $^{1}H$ | $^{15}N$ | CO | Cα | Cβ |
| 247 SER | 8.101 | 114.171 | 170.884 | 55.894 | 62.065 |
| 262 GLN | 8.057 | 118.439 | 172.059 | 52.771 | 28.177 |
| 263 GLY | 8.058 | 106.485 | 169.33 | 42.598 | |
| 264 HIS | 8.187 | 115.241 | 170.538 | 52.187 | 27.487 |
| 265 MET | 8.318 | 118.484 | 171.743 | 52.580 | 31.895 |
| 266 ASN | 8.377 | 118.436 | 171.43 | 50.412 | 37.007 |
| 267 GLN | 8.128 | 117.853 | 172.102 | 52.961 | 27.814 |
| 268 GLY | 8.244 | 106.037 | 169.812 | 42.447 | |
| 269 GLY | 8.156 | 106.084 | 169.084 | 42.485 | |
| 270 GLY | 7.987 | 106.261 | 170.023 | 42.738 | |
| 271 TYR | 8.080 | 115.517 | 172.101 | 55.237 | 36.304 |
| 274 GLY | 8.119 | 105.455 | 169.455 | 42.225 | |
| 275 TYR | 7.993 | 116.738 | 171.66 | 54.821 | 37.222 |
| 276 ASP | 8.416 | 117.225 | 170.915 | 49.852 | 36.470 |
| 277 ASN | 7.999 | 116.415 | 171.126 | 50.018 | 36.977 |
| 278 TYR | 7.992 | 117.36 | 171.908 | 55.225 | 36.792 |
| 279 GLY | 8.259 | 106.399 | 169.805 | 42.535 | |
| 283 TYR | 8.078 | 117.537 | 171.961 | 55.264 | 36.802 |
| 284 GLY | 8.237 | 106.038 | 169.604 | 42.491 | |
| 285 SER | 8.009 | 113.409 | 171.015 | 55.917 | 62.017 |
| 286 GLY | 8.254 | 107.565 | 169.226 | 42.488 | |
| 287 ASN | 8.025 | 116.909 | 171.257 | 53.082 | 39.193 |
| 288 TYR | 7.939 | 117.116 | 171.459 | 54.964 | 37.215 |
| 289 ASN | 8.219 | 116.642 | 171.588 | 50.102 | 37.299 |
| 290 ASP | 8.184 | 117.537 | 171.007 | 49.995 | 36.093 |
| 291 PHE | 8.011 | 116.015 | 171.743 | 54.868 | 37.281 |
| 292 GLY | 8.085 | 105.453 | 169.033 | 42.336 | |
| 293 ASN | 8.068 | 116.864 | 171.586 | 50.203 | 37.466 |
| 294 TYR | 8.116 | 117.628 | 171.742 | 55.219 | 37.222 |
| 295 ASN | 8.218 | 116.854 | 171.227 | 50.650 | 36.268 |
| 296 GLN | 7.798 | 116.639 | 171.519 | 52.519 | 28.257 |
| 297 GLN | 8.062 | 119.023 | 170.491 | 51.507 | 27.152 |
| 299 SER | 8.077 | 112.602 | 170.609 | 55.464 | 62.089 |
| 300 ASN | 8.131 | 118.525 | 171.056 | 50.269 | 37.138 |
| 301 TYR | 7.954 | 116.415 | 171.727 | 54.725 | 37.019 |
| 302 GLY | 8.092 | 105.917 | 167.435 | 42.247 | |
| 304 MET | 8.203 | 116.776 | 171.704 | 52.652 | 31.847 |
| 305 LYS | 7.925 | 117.986 | 172.101 | 52.652 | 31.585 |
| 306 SER | 8.048 | 113.902 | 170.886 | 55.881 | 62.059 |
| 307 GLY | 8.133 | 107.208 | 169.118 | 42.393 | |

| | | | | | |
|---|---|---|---|---|---|
| 308 GLN | 8.121 | 117.225 | 171.38 | 50.107 | 27.222 |
| 309 PHE | 8.169 | 117.763 | 171.932 | 54.522 | 37.126 |
| 310 GLY | 8.370 | 106.082 | 169.884 | 42.469 | |
| 311 GLY | 8.008 | 105.274 | 169.710 | 42.469 | |
| 312 SER | 8.046 | 113.588 | 170.774 | 55.774 | 61.958 |
| 313 ARG | 8.183 | 119.470 | 171.720 | 52.647 | 29.151 |
| 314 ASN | 8.135 | 117.451 | 171.446 | 50.216 | 37.112 |
| 315 MET | 8.086 | 117.853 | 172.057 | 52.520 | 32.889 |
| 316 GLY | 8.235 | 105.501 | 169.649 | 42.170 | |
| 317 GLY | 7.981 | 104.913 | 168.282 | 42.083 | |
| 319 TYR | 8.358 | 118.258 | 172.039 | 54.618 | 37.007 |
| 320 GLY | 8.286 | 106.530 | 169.742 | 42.469 | |
| 321 GLY | 8.149 | 105.633 | 169.24 | 42.292 | |
| 322 GLY | 8.181 | 105.732 | 169.227 | 42.225 | |
| 323 ASN | 8.115 | 117.269 | 171.120 | 50.281 | 37.200 |
| 324 TYR | 7.953 | 116.686 | 171.751 | 54.773 | 37.019 |
| 325 GLY | 8.093 | 105.472 | 167.915 | 42.292 | |
| 327 GLY | 8.425 | 106.706 | 169.578 | 42.297 | |
| 328 GLY | 8.161 | 105.363 | 169.641 | 42.325 | |
| 329 SER | 8.122 | 113.633 | 171.111 | 55.990 | 61.850 |
| 330 GLY | 8.298 | 107.929 | 169.766 | 42.702 | |
| 331 GLY | 8.061 | 105.542 | 169.567 | 42.535 | |
| 332 SER | 8.070 | 113.585 | 171.117 | 55.797 | 61.922 |
| 333 GLY | 8.293 | 107.925 | 169.758 | 42.618 | |
| 334 GLY | 8.031 | 105.363 | 169.274 | 42.386 | |
| 335 TYR | 8.108 | 117.359 | 172.273 | 54.281 | 37.123 |
| 336 GLY | 8.358 | 107.073 | 169.789 | 42.488 | |
| 337 GLY | 8.080 | 105.680 | 169.221 | 42.225 | |
| 338 ARG | 8.070 | 117.672 | 171.953 | 52.607 | 30.868 |
| 339 SER | 8.129 | 114.527 | 170.35 | 55.69 | 62.005 |
| 340 ARG | 8.028 | 118.887 | 171.804 | 52.592 | 29.273 |
| 341 TYR | 8.164 | 117.537 | 173.258 | 54.378 | 36.315 |

**Table S6.** Resonance assignments of TDP-$43^{LCD}$ in 95% DMSO/ 5% $H_2O$ at pH 5.5

| Residue | Assignment (ppm) | | | | |
|---|---|---|---|---|---|
| | $^1H$ | $^{15}N$ | CO | $C_\alpha$ | $C_\beta$ |
| 281 GLY | | | 166.729 | 40.330 | |
| 282 GLY | 8.556 | 107.376 | 168.853 | 42.082 | |
| 283 PHE | 8.393 | 117.372 | 172.143 | 54.638 | 37.616 |
| 284 GLY | 8.420 | 106.808 | 169.633 | 42.374 | |
| 285 ASN | 8.207 | 117.114 | 171.570 | 50.300 | 37.160 |
| 286 GLN | 8.198 | 118.057 | 172.246 | 52.844 | 27.441 |
| 287 GLY | 8.223 | 105.670 | 169.652 | 42.499 | |

| | | | | | |
|---|---|---|---|---|---|
| 288 GLY | 7.991 | 105.232 | 169.380 | 42.207 | |
| 289 PHE | 8.175 | 117.012 | 172.148 | 54.619 | 37.536 |
| 290 GLY | 8.388 | 106.806 | 169.215 | 42.374 | |
| 291 ASN | 8.164 | 116.960 | 171.823 | 49.771 | 37.441 |
| 292 SER | 8.128 | 114.378 | 170.761 | 55.956 | 61.640 |
| 293 ARG | 8.169 | 118.870 | 172.293 | 52.761 | 28.651 |
| 294 GLY | 8.097 | 105.620 | 170.085 | 42.331 | |
| 295 GLY | 8.099 | 105.635 | 169.821 | 42.331 | |
| 296 GLY | 8.182 | 105.954 | 169.286 | 42.164 | |
| 297 ALA | 8.131 | 120.817 | 173.172 | 48.809 | 18.091 |
| 298 GLY | 8.205 | 105.123 | 169.247 | 42.291 | |
| 299 LEU | 7.962 | 118.186 | 173.059 | 51.509 | 40.784 |
| 300 GLY | 8.266 | 106.225 | 169.384 | 42.249 | |
| 301 ASN | 8.129 | 116.873 | 171.504 | 50.356 | 37.232 |
| 302 ASN | 8.234 | 117.529 | 171.48 | 50.216 | 36.785 |
| 303 GLN | 8.040 | 116.879 | 172.143 | 52.594 | 27.441 |
| 304 GLY | 8.121 | 105.594 | 169.408 | 42.248 | |
| 305 SER | 7.954 | 112.870 | 170.611 | 55.371 | 62.058 |
| 306 ASN | 8.328 | 119.539 | 171.541 | 50.300 | 36.868 |
| 307 MET | 8.052 | 117.070 | 172.086 | 52.552 | 31.237 |
| 308 GLY | 8.197 | 105.801 | 169.798 | 42.416 | |
| 309 GLY | 8.112 | 105.448 | 169.793 | 42.416 | |
| 310 GLY | 8.147 | 105.745 | 169.375 | 42.416 | |
| 311 MET | 8.056 | 116.679 | 171.353 | 52.302 | 32.196 |
| 312 ASN | 8.194 | 117.661 | 171.386 | 50.189 | 37.065 |
| 313 PHE | 8.183 | 117.800 | 171.922 | 54.765 | 37.056 |
| 314 GLY | 8.334 | 106.02 | 169.135 | 42.541 | |
| 315 ALA | 7.914 | 120.291 | 172.585 | 48.684 | 18.091 |
| 316 PHE | 8.020 | 115.061 | 171.560 | 54.326 | 37.441 |
| 317 SER | 8.097 | 113.668 | 170.122 | 55.538 | 61.891 |
| 318 ILE | 7.704 | 116.526 | 171.194 | 56.583 | 37.483 |
| 319 ASN | 8.406 | 120.833 | 171.297 | 47.932 | 37.274 |
| 321 ALA | 8.066 | 118.339 | 174.352 | 50.592 | 16.805 |
| 322 MET | 7.653 | 114.376 | 172.989 | 52.863 | 30.713 |
| 323 MET | 7.662 | 116.721 | 172.472 | 53.281 | 31.005 |
| 324 ALA | 8.007 | 120.600 | 173.860 | 49.771 | 17.422 |
| 325 ALA | 7.957 | 119.191 | 173.736 | 49.771 | 17.422 |
| 326 ALA | 7.951 | 118.952 | 173.865 | 49.771 | 17.422 |
| 327 GLN | 7.958 | 115.838 | 172.622 | 53.783 | 27.202 |
| 328 ALA | 8.001 | 120.240 | 173.543 | 49.436 | 17.589 |
| 329 ALA | 7.935 | 119.039 | 173.518 | 49.687 | 17.589 |
| 330 LEU | 7.811 | 116.926 | 173.055 | 52.237 | 40.367 |
| 331 GLN | 7.911 | 116.870 | 172.143 | 52.863 | 27.662 |
| 332 SER | 7.879 | 113.044 | 171.024 | 55.538 | 61.807 |

| | | | | | |
|---|---|---|---|---|---|
| 333 SER | 8.058 | 115.407 | 170.912 | 55.538 | 61.807 |
| 334 TRP | 8.085 | 119.737 | 172.603 | 54.513 | 27.233 |
| 335 GLY | 8.204 | 106.300 | 169.718 | 42.750 | |
| 336 MET | 7.993 | 116.847 | 172.021 | 52.469 | 31.507 |
| 337 MET | 8.102 | 117.400 | 172.138 | 52.469 | 31.507 |
| 338 GLY | 8.147 | 106.130 | 169.380 | 42.374 | |
| 339 MET | 7.948 | 116.400 | 171.664 | 52.362 | 32.134 |
| 340 LEU | 8.033 | 118.864 | 172.641 | 51.275 | 40.539 |
| 341 ALA | 8.069 | 120.830 | 173.069 | 48.768 | 18.007 |
| 342 SER | 7.925 | 111.932 | 170.743 | 55.538 | 61.724 |
| 343 GLN | 8.021 | 118.886 | 171.842 | 52.529 | 27.817 |
| 344 GLN | 8.035 | 117.573 | 171.577 | 52.529 | 27.817 |
| 345 ASN | 8.138 | 117.832 | 171.391 | 50.258 | 37.202 |
| 346 GLN | 8.004 | 117.624 | 171.786 | 52.696 | 27.954 |
| 347 SER | 8.048 | 113.750 | 170.658 | 55.789 | 61.891 |
| 348 GLY | 7.964 | 106.366 | 167.754 | 41.830 | |
| 350 SER | 8.195 | 113.294 | 171.137 | 55.873 | 61.640 |
| 351 GLY | 8.087 | 107.260 | 169.474 | 42.540 | |
| 352 ASN | 8.167 | 117.000 | 171.532 | 50.189 | 37.483 |
| 354 GLN | | 117.529 | 171.814 | 52.761 | 27.483 |
| 355 ASN | 8.122 | 117.004 | 171.701 | 50.197 | 36.898 |
| 356 GLN | 8.071 | 117.926 | 172.556 | 53.178 | 27.400 |
| 357 GLY | 8.290 | 105.964 | 169.384 | 42.249 | |
| 358 ASN | 8.135 | 116.999 | 171.988 | 50.091 | 36.827 |
| 359 MET | 8.054 | 117.396 | 172.068 | 52.613 | 31.465 |

**Table S7.** Average qHDX of Adnectin IBs.

| Fraction Amide Protection | | | | | |
|---|---|---|---|---|---|
| **Residue** | **Average** | **error** | **Residue** | **Average** | **error** |
| S2 | 0.51 | 0.03 | V54 | 0.46 | 0.06 |
| V4 | 0.71 | 0.21 | Y55 | 0.47 | 0.05 |
| D7 | 0.53 | 0.02 | A57 | 0.62 | 0.09 |
| V10 | 0.47 | 0.04 | T58 | 0.44 | 0.02 |
| A12 | 0.41 | 0.06 | I59 | 0.51 | 0.06 |
| A13 | 0.45 | 0.06 | S60 | 0.43 | 0.13 |
| T14 | 0.40 | 0.05 | G61 | 0.42 | 0.10 |
| T16 | 0.43 | 0.08 | K63 | 0.52 | 0.03 |
| L18 | 0.62 | 0.19 | G65 | 0.46 | 0.02 |
| S23 | 0.59 | 0.08 | V66 | 0.56 | 0.13 |
| A24 | 0.68 | 0.06 | Y68 | 0.55 | 0.11 |
| K27 | 0.44 | 0.07 | T69 | 0.57 | 0.07 |
| A29 | 0.65 | 0.08 | V72 | 0.78 | 0.13 |
| T35 | 0.61 | 0.16 | A74 | 0.71 | 0.10 |
| G37 | 0.47 | 0.03 | V75 | 0.67 | 0.12 |
| E38 | 0.53 | 0.06 | L77 | 0.62 | 0.13 |
| G40 | 0.45 | 0.11 | Y81 | 0.57 | 0.06 |
| G41 | 0.45 | 0.07 | G82 | 0.44 | 0.19 |
| N42 | 0.62 | 0.09 | I84 | 0.50 | 0.03 |
| S43 | 0.56 | 0.14 | N87 | 0.52 | 0.04 |
| Q46 | 0.38 | 0.04 | R89 | 0.45 | 0.11 |
| E47 | 0.52 | 0.10 | I92 | 0.58 | 0.09 |
| F48 | 0.57 | 0.11 | | | |
| T49 | 0.44 | 0.08 | | | |
| V50 | 0.56 | 0.08 | | | |

**Table S8.** Average qHDX raSOD1$^{A4V}$ IBs

| **Fraction Amide Protection** | | | | | |
|---|---|---|---|---|---|
| **Residue** | **Average** | **error** | **Residue** | **Average** | **error** |
| L8 | 0.81 | 0.00 | E100 | 0.83 | |
| N19 | 0.60 | 0.03 | S102 | 0.83 | 0.03 |
| F20 | 0.65 | 0.05 | V103 | 0.63 | 0.05 |
| E24 | 0.56 | 0.10 | S107 | 0.53 | 0.11 |
| S25 | 0.48 | | G108 | 0.46 | 0.04 |
| N26 | 0.52 | 0.06 | D109 | 0.37 | 0.06 |
| G27 | 0.46 | 0.11 | H110 | 0.39 | 0.03 |
| V31 | 0.60 | | C111 | 0.28 | 0.05 |
| T39 | 0.49 | 0.08 | I113 | 0.68 | 0.07 |
| H46 | 0.47 | 0.03 | G114 | 0.68 | 0.01 |
| F50 | 0.48 | 0.10 | R115 | 0.42 | 0.08 |
| G51 | 0.50 | 0.06 | H120 | 0.38 | 0.05 |
| D52 | 0.60 | 0.08 | E121 | 0.07 | 0.02 |
| T54 | 0.56 | 0.02 | A123 | 0.54 | |
| A55 | 0.51 | 0.09 | D124 | 0.50 | |
| G56 | 0.50 | 0.07 | L126 | 0.64 | 0.05 |
| C57 | 0.53 | 0.01 | S134 | 0.67 | 0.01 |
| A60 | 0.49 | 0.07 | T137 | 0.52 | 0.01 |
| G61 | 0.50 | 0.10 | G138 | 0.58 | 0.05 |
| H63 | 0.43 | 0.01 | A140 | 0.59 | 0.01 |
| S68 | 0.45 | 0.07 | G141 | 0.50 | 0.02 |
| H71 | 0.55 | 0.03 | S142 | 0.48 | 0.01 |
| G82 | 0.34 | 0.07 | R143 | 0.43 | 0.05 |
| G85 | 0.59 | 0.07 | C146 | 0.65 | 0.06 |
| V87 | 0.62 | 0.08 | G147 | 0.68 | 0.08 |
| T88 | 0.35 | 0.08 | V148 | 0.65 | 0.07 |
| A89 | 0.57 | 0.05 | I149 | 0.54 | 0.07 |
| V94 | 0.48 | 0.07 | G150 | 0.63 | 0.06 |
| A95 | 0.44 | 0.06 | I151 | 0.57 | 0.07 |
| D96 | 0.50 | 0.00 | A152 | 0.52 | 0.01 |
| V97 | 0.65 | 0.01 | Q153 | 0.65 | 0.11 |

**Table S9.** Average qHDX of ApoMb153 IBs

| Fraction Amide Protection | | | | | |
|---|---|---|---|---|---|
| **Residue** | **Average** | **error** | **Residue** | **Average** | **error** |
| L2 | 0.62 | 0.08 | T67 | 0.56 | 0.06 |
| S3 | 0.54 | 0.05 | V68 | 0.61 | 0.09 |
| E4 | 0.54 | 0.10 | L69 | 0.63 | 0.05 |
| G5 | 0.58 | 0.06 | T70 | 0.66 | 0.08 |
| E6 | 0.48 | 0.04 | A71 | 0.65 | 0.17 |
| W7 | 0.49 | 0.07 | G73 | 0.69 | 0.07 |
| Q8 | 0.57 | 0.08 | I75 | 0.61 | 0.04 |
| L9 | 0.77 | 0.11 | K77 | 0.46 | 0.02 |
| V10 | 0.92 | 0.06 | K79 | 0.44 | 0.07 |
| L11 | 0.89 | 0.00 | E85 | 0.81 | 0.17 |
| V13 | 0.91 | 0.06 | K87 | 0.68 | 0.12 |
| W14 | 0.89 | 0.11 | L89 | 0.58 | 0.08 |
| A15 | 0.82 | 0.11 | S92 | 0.64 | 0.07 |
| K16 | 0.63 | 0.09 | K96 | 0.66 | 0.07 |
| V17 | 0.69 | 0.03 | K98 | 0.62 | 0.07 |
| E18 | 0.54 | 0.05 | I99 | 0.58 | 0.12 |
| A19 | 0.65 | 0.12 | Y103 | 0.37 | 0.04 |
| V21 | 0.62 | 0.04 | E105 | 0.40 | 0.07 |
| A22 | 0.63 | 0.08 | F106 | 0.53 | 0.07 |
| I28 | 0.57 | 0.11 | S108 | 0.60 | 0.04 |
| L29 | 0.58 | 0.04 | E109 | 0.52 | 0.05 |
| I30 | 0.53 | 0.04 | I111 | 0.64 | 0.16 |
| R31 | 0.42 | 0.02 | I112 | 0.74 | 0.09 |
| L32 | 0.50 | 0.08 | V114 | 0.44 | 0.06 |
| F33 | 0.38 | 0.05 | L115 | 0.50 | 0.07 |
| K34 | 0.44 | 0.08 | G124 | 0.68 | 0.09 |
| S35 | 0.53 | 0.05 | A125 | 0.82 | 0.13 |
| T39 | 0.83 | 0.11 | Q128 | 0.67 | 0.05 |
| L40 | 0.68 | 0.08 | G129 | 0.66 | 0.05 |
| K42 | 0.74 | 0.05 | M131 | 0.50 | 0.04 |
| K47 | 0.70 | 0.04 | N132 | 0.62 | 0.08 |
| L49 | 0.62 | 0.07 | K133 | 0.56 | 0.05 |
| K50 | 0.72 | 0.10 | L135 | 0.51 | 0.03 |
| T51 | 0.84 | 0.10 | I142 | 0.41 | 0.03 |
| A53 | 0.91 | 0.11 | A143 | 0.31 | 0.04 |
| K56 | 0.72 | 0.02 | A144 | 0.42 | 0.07 |
| A57 | 0.70 | 0.09 | Y146 | 0.33 | 0.04 |
| S58 | 0.68 | 0.09 | L149 | 0.39 | 0.02 |
| L61 | 0.62 | 0.04 | G150 | 0.56 | 0.09 |
| K62 | 0.69 | 0.06 | Q152 | 0.68 | 0.07 |
| V66 | 0.43 | 0.02 | | | |

**Table S10.** Average qHDX of ApoMb77 IBs.

| Fraction Amide Protection | | | | | |
|---|---|---|---|---|---|
| **Residue** | **Average** | **error** | **Residue** | **Average** | **error** |
| L2 | 0.52 | 0.06 | T39 | 0.57 | 0.03 |
| S3 | 0.55 | 0.11 | L40 | 0.54 | 0.04 |
| E4 | 0.45 | 0.02 | K42 | 0.65 | 0.13 |
| G5 | 0.56 | 0.07 | F46 | 0.47 | 0.02 |
| E6 | 0.41 | 0.09 | K47 | 0.34 | 0.02 |
| W7 | 0.50 | 0.11 | L49 | 0.36 | 0.05 |
| Q8 | 0.58 | 0.07 | T51 | 0.45 | 0.06 |
| L9 | 0.83 | 0.18 | E54 | 0.33 | 0.02 |
| V10 | 0.91 | 0.07 | K56 | 0.51 | 0.07 |
| L11 | 1.01 | 0.06 | A57 | 0.48 | 0.04 |
| V13 | 1.07 | 0.08 | S58 | 0.44 | 0.09 |
| W14 | 0.98 | 0.05 | L61 | 0.57 | 0.03 |
| A15 | 1.06 | 0.10 | K62 | 0.51 | 0.08 |
| V17 | 0.64 | 0.12 | K63 | 0.31 | 0.03 |
| E18 | 0.59 | 0.07 | V66 | 0.38 | 0.04 |
| A19 | 0.59 | 0.05 | T67 | 0.46 | 0.05 |
| V21 | 0.55 | 0.04 | V68 | 0.66 | 0.05 |
| A22 | 0.57 | 0.08 | L69 | 0.66 | 0.15 |
| I28 | 0.88 | 0.06 | T70 | 0.82 | 0.06 |
| I30 | 0.84 | 0.13 | A71 | 0.63 | 0.04 |
| R31 | 0.79 | 0.05 | L72 | 0.80 | 0.12 |
| L32 | 0.80 | 0.10 | G73 | 0.68 | 0.05 |
| F33 | 0.53 | 0.05 | I75 | 0.67 | 0.06 |
| K34 | 0.48 | 0.05 | L76 | 0.55 | 0.06 |
| S35 | 0.45 | 0.07 | K77 | 0.52 | 0.05 |

**Table S11.** Average qHDX hnRNPA2$^{LCD}$ IBs

| Fraction Amide Protection | | | | | |
|---|---|---|---|---|---|
| **Residue** | **Average** | **error** | **Residue** | **Average** | **error** |
| S2 | 0.36 | 0.09 | K305 | 0.37 | 0.09 |
| Q17 | 0.98 | 0.05 | S306 | 0.34 | 0.09 |
| M20 | 0.72 | 0.13 | G307 | 0.32 | 0.09 |
| Q267 | 0.59 | 0.15 | N308 | 0.43 | 0.06 |
| G269 | 0.19 | 0.05 | F309 | 0.34 | 0.11 |
| G270 | 0.23 | 0.08 | G310 | 0.27 | 0.07 |
| Y271 | 0.28 | 0.09 | G311 | 0.23 | 0.05 |
| G274 | 0.28 | 0.00 | S312 | 0.23 | 0.09 |
| Y275 | 0.31 | 0.00 | R313 | 0.27 | 0.09 |
| N277 | 0.47 | 0.08 | M315 | 0.49 | 0.03 |
| G279 | 0.39 | 0.00 | G316 | 0.38 | 0.01 |
| Y283 | 0.46 | 0.01 | G317 | 0.28 | 0.08 |
| S285 | 0.35 | 0.07 | Y319 | 0.32 | 0.00 |
| N287 | 0.47 | 0.01 | G320 | 0.30 | 0.07 |
| N289 | 0.63 | 0.10 | G321 | 0.24 | 0.03 |
| F291 | 0.56 | 0.29 | G322 | 0.34 | 0.22 |
| N293 | 0.37 | 0.10 | Y324 | 0.53 | 0.03 |
| Y294 | 0.60 | 0.05 | G327 | 0.23 | 0.08 |
| N295 | 0.43 | 0.04 | G328 | 0.18 | 0.02 |
| Q296 | 0.37 | 0.06 | S329 | 0.29 | 0.04 |
| Q297 | 0.39 | 0.11 | G331 | 0.24 | 0.07 |
| S299 | 0.34 | 0.07 | S332 | 0.26 | 0.06 |
| N300 | 0.40 | 0.14 | G334 | 0.24 | 0.05 |
| Y301 | 0.38 | 0.09 | G336 | 0.28 | 0.06 |
| G302 | 0.42 | 0.03 | S339 | 0.30 | 0.08 |
| M304 | 0.39 | 0.07 | R340 | 0.30 | 0.11 |

**Table S12. Average qHDX TDP-43$^{LCD}$ IBs**

| Fraction Amide Protection | | | | | |
|---|---|---|---|---|---|
| **Residue** | **Average** | **error** | **Residue** | **Average** | **error** |
| F283 | 0.23 | 0.03 | M322 | 0.58 | 0.02 |
| G284 | 0.13 | 0.03 | M323 | 0.62 | 0.03 |
| N285 | 0.50 | 0.02 | A324 | 0.71 | 0.06 |
| Q286 | 0.18 | 0.03 | A325 | 0.72 | 0.08 |
| G287 | 0.17 | 0.05 | A326 | 0.80 | 0.05 |
| G288 | 0.14 | 0.02 | Q327 | 0.80 | 0.09 |
| G290 | 0.20 | 0.08 | A328 | 0.80 | 0.06 |
| S292 | 0.20 | 0.06 | A329 | 0.78 | 0.02 |
| R293 | 0.15 | 0.02 | L330 | 0.93 | 0.04 |
| A297 | 0.14 | 0.00 | Q331 | 0.76 | 0.08 |
| G298 | 0.14 | 0.06 | S332 | 0.75 | 0.09 |
| L299 | 0.12 | 0.01 | S333 | 0.82 | 0.09 |
| G300 | 0.13 | 0.05 | W334 | 0.82 | 0.06 |
| N302 | 0.18 | 0.00 | G335 | 0.95 | 0.08 |
| Q303 | 0.18 | 0.04 | M336 | 0.99 | 0.03 |
| G304 | 0.17 | 0.02 | M337 | 0.97 | 0.05 |
| S305 | 0.18 | 0.05 | G338 | 0.85 | 0.08 |
| N306 | 0.20 | 0.03 | M339 | 0.71 | 0.05 |
| M307 | 0.20 | 0.06 | L340 | 0.46 | 0.04 |
| G308 | 0.22 | 0.03 | A341 | 0.50 | 0.03 |
| G309 | 0.21 | 0.01 | S342 | 0.26 | 0.02 |
| G310 | 0.25 | 0.04 | Q344 | 0.22 | 0.05 |
| M311 | 0.33 | 0.03 | N345 | 0.21 | 0.08 |
| N312 | 0.36 | 0.05 | Q346 | 0.17 | 0.03 |
| F313 | 0.37 | 0.06 | S347 | 0.13 | 0.02 |
| G314 | 0.31 | 0.01 | G348 | 0.13 | 0.02 |
| A315 | 0.34 | 0.03 | S350 | 0.20 | 0.04 |
| F316 | 0.28 | 0.04 | G351 | 0.20 | 0.02 |
| S317 | 0.27 | 0.03 | Q356 | 0.19 | 0.04 |
| I318 | 0.23 | 0.01 | G357 | 0.24 | 0.03 |
| N319 | 0.20 | 0.02 | M359 | 0.21 | 0.07 |
| A321 | 0.36 | 0.03 | | | |

**Table S13.** FTIR second derivative peaks for IB and purified proteins.

| | | Antiparallel β-sheet ($cm^{-1}$) | β-turn ($cm^{-1}$) | Unst./α ($cm^{-1}$) | β-sheet ($cm^{-1}$) |
|---|---|---|---|---|---|
| IB | Adnectin | 1689 | 1677 | 1654 | 1630 |
| | raSOD1$_{A4V}$ | 1692 | 1680 | 1654 | 1629 |
| | ApoMb153 | 1695 | 1680 | 1654 | 1624 |
| | ApoMb77 | 1695 | 1677 | 1654 | 1624 |
| | hnRNPA2$_{LCD}$ | - | 1681 | 1654 | 1630 |
| | TDP43$_{LCD}$ | - | 1677 | 1654 | 1630 |
| Purified | Adnectin | 1688 | | | 1630 |
| | raSOD1$_{A4V}$ | 1687 | | 1651 | 1630 |
| | ApoMb153 | | 1681 | 1653 | 1629 |
| | ApoMb77 | 1691 | - | 1654 | 1624 |

**Table S14.** Results from curve fitting of ATR-FTIR spectra.

| | Protein | β-turn/β-antiparallel | | | α-helix/unstructured | | | β-sheet | | |
|---|---|---|---|---|---|---|---|---|---|---|
| | | Center ($cm^{-1}$) | area (%) | FWHM ($cm^{-1}$) | Center ($cm^{-1}$) | area (%) | FWHM ($cm^{-1}$) | Center ($cm^{-1}$) | area (%) | FWHM ($cm^{-1}$) |
| IB | Adnectin | 1675.9±2.0 | 17.9±4.1 | 26.3±2.1 | 1653.1±0.5 | 35.3±6.4 | 25.3±2.0 | 1630.2±0.6 | 50.2±3.3 | 27.3±0.6 |
| | raSOD1$_{A4V}$ | 1676.4±0.3 | 15.1±0.8 | 25.0±0.4 | 1653.5±0.7 | 42.0±4.7 | 26.5±1.2 | 1629.6±0.5 | 45.9±3.6 | 27.0±0.4 |
| | ApoMb 153 | 1675.9±0.5 | 13.6±1.3 | 25.3±0.7 | 1651.7±0.3 | 52.1±3.2 | 27.9±0.9 | 1627.4±0.5 | 37.1±1.8 | 25.4±0.3 |
| | ApoMb 77 | 1674.2±1.2 | 18.5±2.9 | 28.2±1.7 | 1651.6±0.8 | 38.9±4.4 | 25.8±1.2 | 1627.8±0.5 | 45.0±2.0 | 24.9±0.6 |
| | hnRNPA2 | 1674.2±1.4 | 18.2±4.2 | 25.8±1.7 | 1651.9±0.5 | 46.3±7.0 | 27.8±1.7 | 1629.0±0.3 | 39.0±3.5 | 26.2±0.7 |
| | TDP43 | 1677.1±0.3 | 16.6±1.0 | 24.1±0.3 | 1653.4±0.1 | 54.9±2.8 | 28.0±0.6 | 1630.7±0.7 | 31.5±2.6 | 25.9±0.4 |
| Purified | Adnectin | 1683.6±1.5 | 7.8±22.6 | 19.0±1.9 | 1661.6±2.7 | 25.4±1.3 | 28.9±0.4 | 1632.3±0.7 | 67.1±3.0 | 29.0±0.5 |
| | raSOD1$_{A4V}$ | 1674.4±1.0 | 22.0±3.0 | 26.2±1.1 | 1652.0±0.8 | 39.0±3.4 | 26.6±0.9 | 1631.1±0.6 | 39.2±2.1 | 27.5±0.6 |
| | ApoMb 153 | 1677.0±0.5 | 9.4±1.0 | 20.8±0.7 | 1652.6±0.5 | 66.2±4.6 | 27.6±0.7 | 1630.1±0.6 | 24.5±4.1 | 25.5±0.8 |
| | ApoMb 77 | 1673.2±0.3 | 18.6±0.8 | 29.5±0.0 | 1650.1±0.2 | 39.9±2.1 | 26.4±0.9 | 1626.8±0.1 | 41.8±1.3 | 24.0±0.2 |

The amide I band was fit using three Voigt components: β-turn/antiparallel β-sheet (~1680 $cm^{-1}$), unstructured/α-helix (~1654 $cm^{-1}$) and β-sheet (~1630 $cm^{-1}$). The fitted spectra are shown in Figure S8.
